# Co-transcriptional RNA strand displacement circuits

**DOI:** 10.1101/2021.07.20.450530

**Authors:** Samuel W. Schaffter, Elizabeth A. Strychalski

## Abstract

Engineered molecular circuits that process information in biological systems could address emerging human health and biomanufacturing needs. However, such circuits can be difficult to rationally design and scale. DNA-based strand displacement reactions have demonstrated the largest and most computationally powerful molecular circuits to date but are limited in biological systems due to the difficulty in genetically encoding components. Here, we develop scalable co-transcriptional RNA strand displacement (ctRSD) circuits that are rationally programmed *via* base pairing interactions. ctRSD addresses the limitations of DNA-based strand displacement circuits by isothermally producing circuit components *via* transcription. We demonstrate the programmability of ctRSD *in vitro* by implementing logic and amplification elements, and multi-layer signaling cascades. Further, we show ctRSD kinetics are accurately predicted by a simple model of coupled transcription and strand displacement, enabling model-driven design. We envision ctRSD will enable rational design of powerful molecular circuits that operate in biological systems, including living cells.

## Introduction

A major goal of synthetic biology is developing programmable molecular circuits that can be rationally engineered to process information in biological systems. Such circuits have the potential to address emerging challenges in human health and disease (*1*), agriculture (*2*), and biomanufacturing (*3*). To meet these diverse needs, molecular circuits must be scalable, modular, and rationally programmable to execute operations like logic, signal amplification, and multi-layer cascades. Further, circuits capable of a wide range of computations beyond Boolean logic, such as molecular pattern recognition (*4*), could greatly expand existing capabilities. A key challenge to developing such circuits is identifying molecular components that not only meet the above criteria, but also behave predictably to enable model-driven design.

The predictable and programmable Watson-Crick base pairing interactions of nucleic acids has led to their adoption as versatile components for molecular circuit programming. In particular, *in vitro* circuits based on toehold-mediated strand displacement (TMSD) reactions have demonstrated sophisticated digital computations and mathematical operations (*5*), molecular pattern recognition (*4, 6*), signal cascades (*7*) and amplifiers (*5, 8, 9*), and complex dynamics (*9–11*). In TMSD reactions, a single-stranded input binds to a double-stranded nucleic acid gate *via* a single-stranded toehold domain and displaces an output strand with a new exposed toehold that can facilitate further TMSD reactions (Fig. 1A). Interactions between inputs and gates are programmed through sequence complementarity, and the combinatorial nucleic acid sequence space allows TMSD reaction networks to be scaled up to >100 components (*4*). Additionally, reaction kinetics can be tuned over six orders of magnitude by simply changing the length of the toehold (*12*). These properties have enabled predictive models of TMSD circuit behavior that allow circuit design abstraction (*13*).

**Fig. 1.**
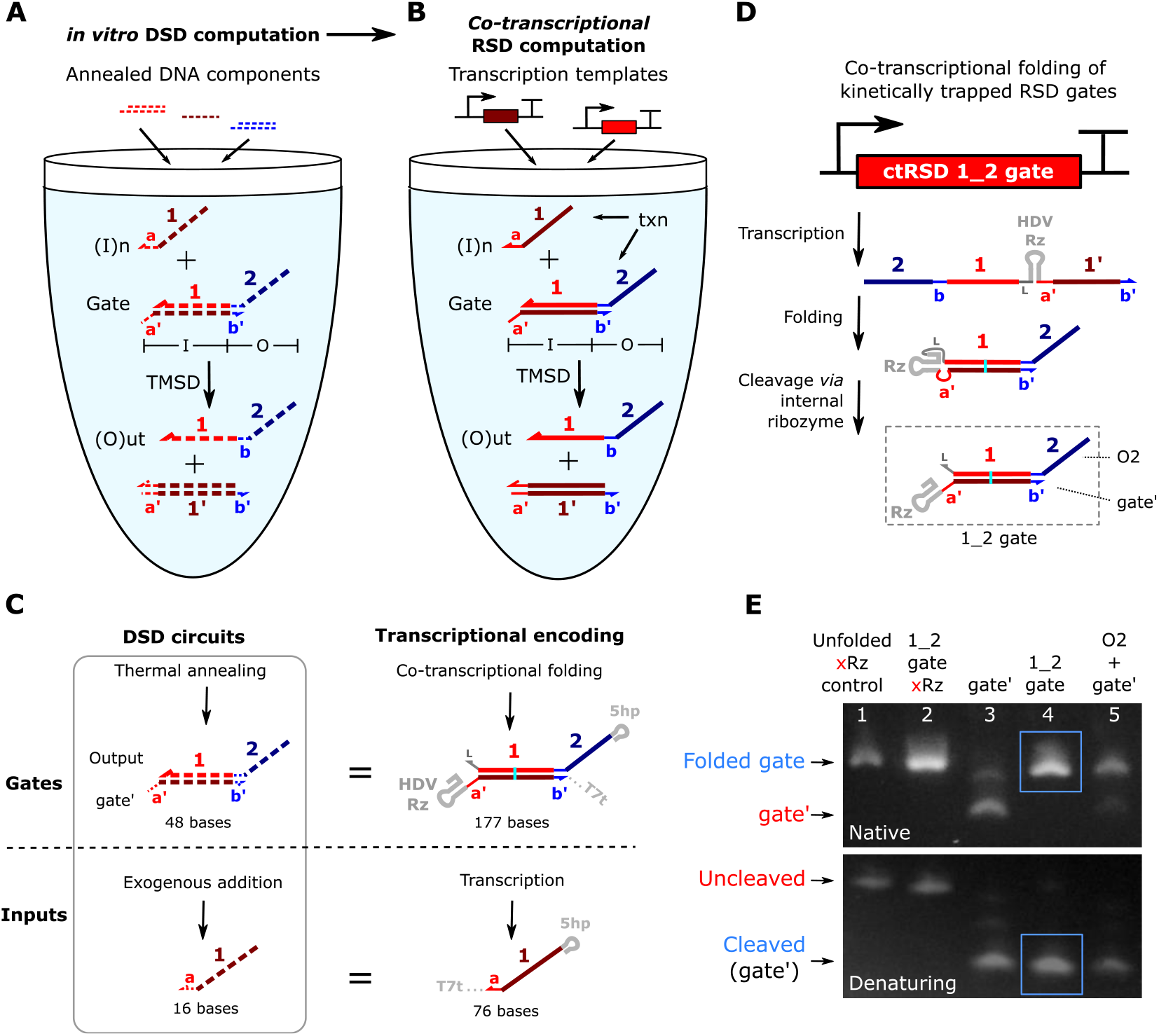
Co-transcriptional RNA strand displacement (ctRSD) design. (**A**) In DNA strand displacement (DSD), pre-annealed DNA gates are mixed to build a circuit. Strand exchange between the input and gate releases an output. (**B**) In ctRSD, designed transcription templates produce the RNA components that make up a circuit. DNA and RNA are represented with dashed and solid lines, respectively. Bold letters and numbers represent sequence identity. A prime (‘) denotes complementarity. The I and O below the gate represent input and output domains, respectively. (**C**) Transcriptional encoding of ctRSD components. All RNAs possess a 5’ hairpin (5hp) and a 3’ terminator (T7t). For simplicity, these motifs are omitted elsewhere. The cyan line represents a G-U wobble pairing. The gate contains a self-cleaving ribozyme (HDV Rz) to enable co-transcriptional folding of kinetically trapped gates (D). See Supplementary Fig. 1 for schematics with sequences. (**D**) In ctRSD, gates fold into RNA hairpins that self-cleave to produce a reactive dsRNA products. Input and output domains define gate names (*e.g*. “1_2 gate”). (**E**) Gel electrophoresis demonstrating gate folding and cleavage (Lane 4, blue box). Lane 1: a transcript that is the same length as the gate but does not fold into a hairpin or cleave (xRz). Lane 2: the 1_2 gate without cleavage (xRz). Lane 3: the gate’ strand (Rz, a’-, 1’-, and b’-domains) alone. Lane 5: separate transcription of the output (O2) and gate’ strands. The 46 base single-stranded O2 strand stained poorly for visualization (*53*). See Supplementary Fig. 2 for control transcript designs.

Because TMSD circuits are composed of nucleic acids, they have great potential for integration with biological systems. However, these circuits have primarily been implemented *in vitro* using DNA components that are not easily genetically encoded (*14, 15*). This restricts their applications in synthetic biology, particularly *in vivo* (*16*). A key challenge to operating TMSD circuits in biological systems is developing a method to isothermally prepare all circuit components in a single reaction. Typically, TMSD components are thermally annealed separately to prevent spurious reactions between gates and then mixed to make a circuit (*4–6*). Thus, these circuits currently cannot be continuously produced in the same place they are operated. Although TMSD circuits can be prepared and then added to biological samples (*17*) or transfected into cells (*18*) at fixed concentrations, these implementations are only single use (*19–21*) and circuit lifetime is limited by component degradation. A method to continuously produce TMSD circuits *in situ* could greatly expand their capabilities. Genetically encoded RNA-based circuits that utilize strand displacement have been developed (*22–27*) and other transcription-based circuits have achieved some of the capabilities of TMSD circuits (*28, 29*). However, these systems have yet to demonstrate the predictive design and scale up seen in state-of-the-art DNA-based circuits.

Here, we develop scalable and programmable co-transcriptional RNA strand displacement (ctRSD) circuits. In ctRSD, circuit components isothermally self-assemble during transcription and execute programmed computations in the same reaction. We validate ctRSD circuit performance *in vitro* by building circuits that execute logic, signal amplification, and multi-layer cascades. We demonstrate the scalability and modularity of the ctRSD by successfully implementing 13 ctRSD gates in 8 different circuit topologies. We find ctRSD kinetics are well predicted by a simple model of coupled transcription and strand displacement that assumes uniform kinetic behavior across gates, facilitating predictive circuit engineering. Further, ctRSD circuits are designed so that state-of-the-art DNA-based circuits capable of neural network computations and pattern recognition (*4, 6*) could be directly adopted. ctRSD should enable the power of TMSD circuits to be realized in biological systems for smart diagnostics or sensors (*6, 30, 31*). Ultimately, ctRSD circuits could be genetically encoded and continuously operated inside living cells.

## Results

### Design of co-transcriptional RNA strand displacement (ctRSD) components

To develop ctRSD, we sought a system in which modular and programmable strand displacement circuit components could be isothermally produced *via* transcription. In TMSD circuits, modularity is achieved by designing toehold exchange gates that allow any input sequence to be converted into any output sequence through a gate (*4, 5, 7, 12*). For example, in Fig. 1A, the input domain is composed of the a’-toehold and the 1:1’-domain duplex, both of which are complementary to the input, I1. The output domain is composed of the sequestered b-toehold and the 2-domain overhang, neither of which share complementarity with I1. Thus, input and output domain sequences are completely independent. We adopted an analogous modular gate design for ctRSD (Fig. 1B). In these toehold exchange gates, the b-toehold of the output is sequestered in a duplex, kinetically precluding a reaction downstream unless the gate input is present. In DNA-based circuits, the toehold exchange gates are thermally annealed in separate test tubes to kinetically trap the outputs before circuit components are mixed. In ctRSD circuits, the RNA toehold exchange gates must isothermally assemble into kinetically trapped intermediates in a single pot during transcription. Simply transcribing the two gate strands separately and allowing them to hybridize to form a gate is not a viable option as the output strand of the gate can also react with downstream gates, introducing significant leak (Supplementary Fig. 7).

To transcriptionally encode kinetically trapped RNA toehold exchange gates, we inserted a self-cleaving RNA ribozyme motif between the two strands of the gate (Fig. 1C). This motif allows us to encode RNA gates as single transcripts that fold into hairpins and then cleave to yield reactive gates (Fig. 1D). Co-transcriptional folding is at least one order of magnitude faster than transcription (*32*), so the RNA gates should fold before they have time to react downstream. The self-cleaving ribozyme also ensures 1:1 stoichiometry between the gate strands, further reducing the potential for leaks (*33*). Inclusion of the ribozyme motif is critical, as the co-transcriptionally folded RNA gate exhibited >7-fold lower downstream leak rate than transcribing the two strands of the RNA gate separately (Supplementary Fig. 8). A 5’ hairpin motif and a 3’ hairpin terminator for T7 RNAP were also appended to gates and inputs (Fig. 1C). The 5’ hairpin contains the T7 RNAP consensus initiation sequence to facilitate efficient and uniform transcription across components (*34, 35*). The terminator hairpin reduces unwanted products associated with runoff transcription (*36, 37*) and enables incorporation into plasmids.

Fig. 1C shows the final selection for our ctRSD gate design, however, there are many alternative implementations that would embody the same general features (Supplementary Section 2). To optimize gate performance, we analyzed four considerations when selecting the final design: 1) directionality of the single-stranded toehold, 2) domain sequence identity, 3) domain transcription order, and 4) self-cleaving ribozyme choice. We designed the ctRSD gates with 5’ toeholds because a 5’ toehold on an RNA gate allows the invading strand to participate in co-axial base stacking, increasing the binding strength compared to a 3’ toehold (*38, 39*). We restricted the gate output sequences to cytosine (C), adenine (A), or uracil (U) bases. This sequence constraint reduces unwanted secondary structure or dimerization of single-stranded components (*4, 5, 7*). A G-U wobble pair was also introduced in the middle of the hybridized portion of the gate to reduce DNA template synthesis errors (*40*) and to drive the forward ctRSD reaction with inputs that convert the G-U wobble to a G-C pair (*41*). The 5’ end of the output strand of the gate was selected as the starting point for transcription so that the first sequence produced would only possess C, A, and U bases, preventing co-transcriptional folding into undesired secondary structure. This transcription order ensures that the G, A, and U restricted sequence of the strand that hybridizes to the output strand (*i.e*. the gate’ strand) is transcribed after its complementary sequence to promote folding of the RNA gate stem over alternative structures with G-U wobbles (Supplementary Section 2.2). For the self-cleaving ribozyme, we selected a variant of the hepatitis delta virus (HDV) ribozyme (*42*) (Supplementary Section 2.3). This ribozyme has no upstream or downstream sequence constraints, has a very stable fold (*43*), and has been reported to cleave itself with a rate constant of nearly 1 s^−1^ *in vivo* (*44*).

We used native and denaturing agarose gel electrophoresis to confirm the ctRSD gate fold and cleave as designed. On a native gel, the ctRSD gate (lane 4, Fig. 1E) was the same size as a control sample in which the two strands of the gate were transcribed from separate templates (lane 5, Fig. 1E), indicating full length gate production and folding. On a denaturing gel, the primary product from the ctRSD gate (lane 4, Fig. 1E) migrated faster than the uncleaved control transcript (lane 2, Fig. 1E) and was the same size as the gate’ strand alone (lane 3, Fig. 1E), indicating ribozyme cleavage. Importantly, the cleavage reaction is efficient and fast; we observed >90 % cleavage in less than 15 min with an estimated cleavage rate constant of 0.25 min^−1^ (Supplementary Fig. 12).

### Experimental characterization and modeling of ctRSD

We next sought to characterize the ctRSD reaction in which a gate and its corresponding input are co-transcribed and react *via* strand displacement to release an output strand (Fig. 1B). The I1:gate’ product of the strand displacement reaction is a higher molecular weight than the unreacted gate, so we first analyzed the reaction with native gel electrophoresis (Fig. 2A). Increasing concentrations of I1 template increased the percentage of I1:gate’ product on the gel, with a 2:1 mixture of the I1 and 1_2 gate templates yielding 100 % product (lane 3 to lane 7, Fig. 2B). Assuming the transcription rates of I1 and the 1_2 gate are approximately equal, the fraction of I1:gate’ produced with increasing I1 template concentration provides information about the thermodynamics of the ctRSD reaction. We found the unreacted 1_2 gate percentages across input concentrations in experiments were within ≈12 % of the thermodynamic predictions from NUPACK 3.2.2 (*45*) (Fig. 2B and Supplementary Section 3).

**Fig. 2.**
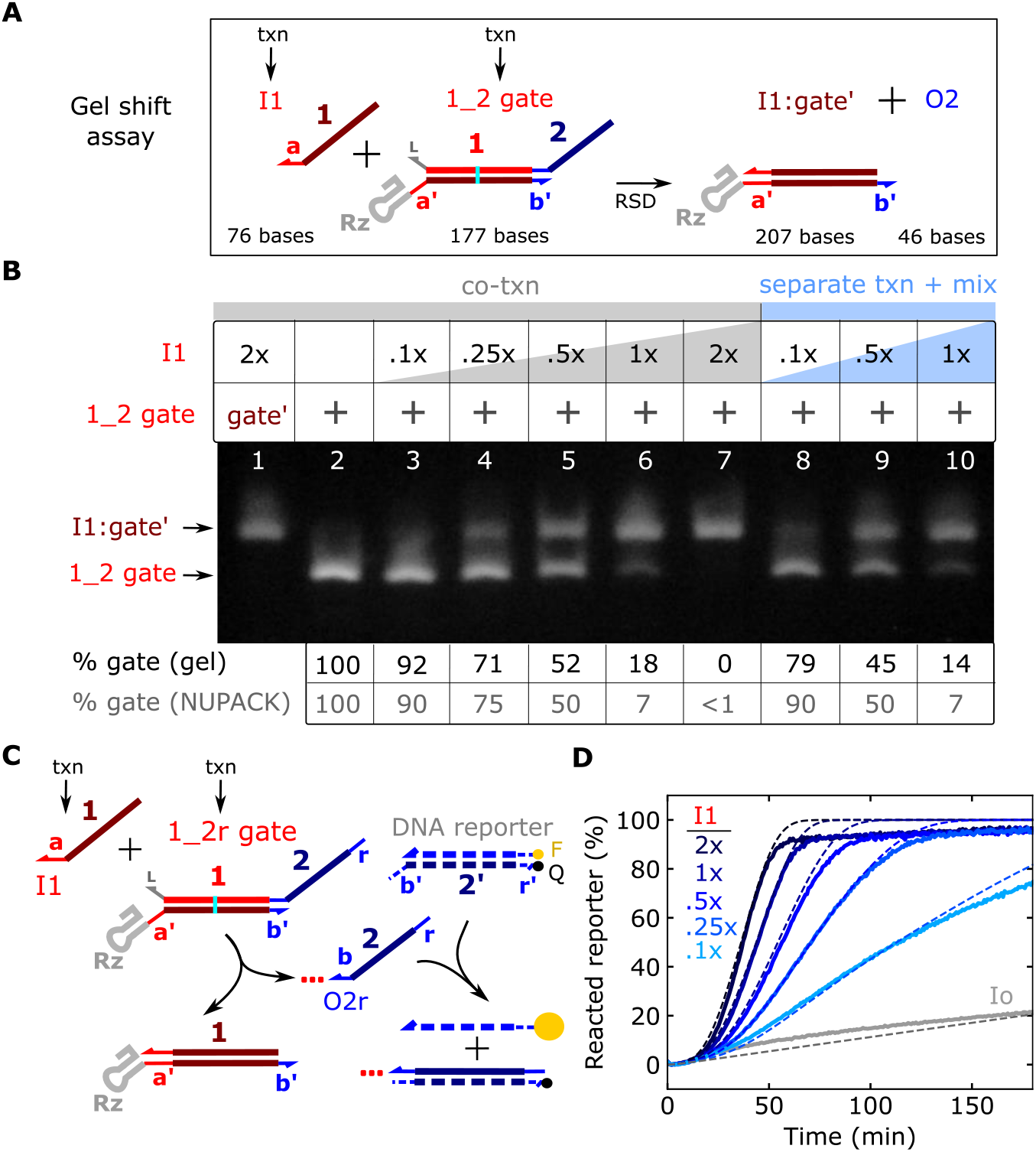
Characterization of the ctRSD reaction. (**A**) The ctRSD reaction schematic. The Il:gate’ complex is 30 bases longer than the gate. (**B**) Native RNA gel electrophoresis results demonstrating ctRSD. Lane 1: Il:gate’ complex. Lane 2: 1_2 gate. Lane 3 to lane 7: 25 nmol/L of 1_2 gate template was co-transcribed with 2.5 nmol/L (0.1x) to 50 nmol/L (2x) of I1 template. The 46 base output strand of the gate (O2) was not visible (*53*). Electrophoresis was conducted 2 h DNase I addition. Lane 8 to lane 10: I1 and 1_2 gate templates were transcribed separately for 30 min, DNase I was added for 30 min, then samples were mixed in equal volumes and incubated at 37 °C for 2 h before electrophoresis. The table below the gel shows the percentage of 1_2 gate in each lane agrees with NUPACK predictions. (**C**) Schematic of the fluorescent DNA reporter assay to track O2r production. The red dotted line trailing O2r represents the upstream portion of the output strand not involved in downstream reactions. (**D**) Experimental (solid lines) and simulated (dashed lines) DNA reporter signal during co-transcription of the 1_2r gate and different I1 template concentrations. The gray lines indicate the 1_2r gate co-transcribed with a randomized input sequence (Io) that does interact with the 1_2r gate. DNA template and T7 RNAP concentrations are tabulated in Supplement Table 4. See Supplementary Section 4 for simulation details.

The results in lane 3 to lane 7 in Fig. 2B were obtained from simultaneous transcription of I1 and the 1_2 gate, so the observed reaction between the two transcripts could result from I1 binding to the 1_2 gate prior to folding, rather than strand displacement. To rule out this potential reaction pathway, we transcribed the I1 and the 1_2 gate RNAs separately and then mixed them together. Separate transcription followed by mixing yielded similar results to co-transcription (lane 8 to lane 10, Fig. 2B), suggesting I1 and the 1_2 gate react *via* the designed strand displacement mechanism.

To explore ctRSD kinetics, we co-transcribed the input and gate templates alongside a DNA reporter complex designed to release a fluorescent signal upon reaction with the gate output strand (Fig. 2C). We opted to use a DNA-based reporter, rather than an RNA aptamer-based reporter (*46, 47*), because the DNA reporter is easily calibrated to output concentration for modeling (*4, 5, 7*). To be stable at 37 °C, we designed the reporter with a 16 base duplex. The 5’ end of the 1_2 gate was extended to include the full complement of the reporter (1_2r gate) to ensure an irreversible reaction. We fixed the 1_2r gate template concentration and varied the I1 template concentration. To ensure the same transcriptional load for comparison, a template that produced an unreactive input (Io) was added to maintain the same total input template concentration across samples (Methods). As expected from mass action kinetics, increasing concentrations of the I1 template resulted in faster ctRSD reaction kinetics (Fig. 2D). A gate with a mutant ribozyme that cannot cleave resulted in >3-fold slower output production (Supplementary Fig. 15). We also found transcription of the 1_2r gate with Io alone resulted in ≈20 % of the maximum DNA reporter signal, indicating a slow leak reaction (Fig. 2D). The magnitude of this leak depended on T7 RNAP and total template concentrations (Supplementary Fig. 17).

We next investigated whether a mass action kinetic model of coupled transcription, ribozyme cleavage, and RNA strand displacement (Supplementary Section 4.1) could recapitulate the ctRSD kinetics observed in experiments. For model parameters, we used the ribozyme cleavage rate that we measured (Supplementary Fig. 12) and estimated order of magnitude strand displacement rate constants consistent with previous literature (Supplementary Section 4.2). We calibrated the transcription rate constant for each experiment with a control sample (Methods). Our initial model did not include any terms to describe the leak observed when the 1_2r gate was transcribed without the correct input and thus could not capture that effect (Supplementary Fig. 18).

To investigate the source of the leak, we evaluated how well incorporating plausible leak pathways into the model recapitulated the experimental leak kinetics. We first evaluated a leak pathway in which the cleaved 1_2r gate could directly react with the DNA reporter *via* a 0 base toehold (*33*). In simulations, this model exhibited a lag time before the leak was observed, inconsistent with experiments (Supplementary Fig. 18). We next introduced a leak pathway in which the 1_2r gate could react with the DNA reporter prior to folding. In simulations, this leak pathway closely recapitulated the observed leak kinetics using a folding rate constant consistent with T7 RNAP transcription (Supplementary Fig. 18). To experimentally investigate the presence of this leak pathway, we transcribed the 1_2r gate in the absence of DNA reporter, heat denatured T7 RNAP, and then added the DNA reporter to the solution containing the folded 1_2r gate. If the leak pathway involved the unfolded 1_2r gate, no signal should be observed upon reporter addition. We found that reporter addition resulted in an instantaneous jump in fluorescence, and the magnitude of the leak signal increased with increasing 1_2r gate transcription time (Supplementary Fig. 19). From these results, we reasoned the leak is not due to a reaction with the 1_2r gate prior to folding, but rather due to the presence of a 1_2r gate product that is highly reactive. Such an unintended side product could be the result of premature termination or gate misfolding events that leave the b-toehold of the gate exposed to rapidly react with the DNA reporter. We modeled this leak reaction by assuming the 1_2r gate template directly produced output at a leak transcription rate. In the model, a leak transcription rate of 3 % the gate transcription rate recapitulated the experimental kinetics (Supplementary Fig. 18). We included this leak term in all subsequent simulations. With the inclusion of this leak term, the kinetic model exhibited good agreement with experimental ctRSD kinetics (Fig. 2D).

Using the same design as the 1_2r gate, we created three more ctRSD gate sequences with corresponding inputs. We reused the same input toehold sequence across gates to facilitate similar strand displacement kinetics (*4, 7*). These gate sequences cleaved with similar efficiency as the 1_2r gate (Supplementary Fig. 20) and exhibited nearly identical ctRSD kinetics as the 1_2r gate (Fig. 3A). Importantly, I1, I3, I4, I5 only reacted with their designed gate (Fig. 3B), demonstrating orthogonality.

**Fig. 3.**
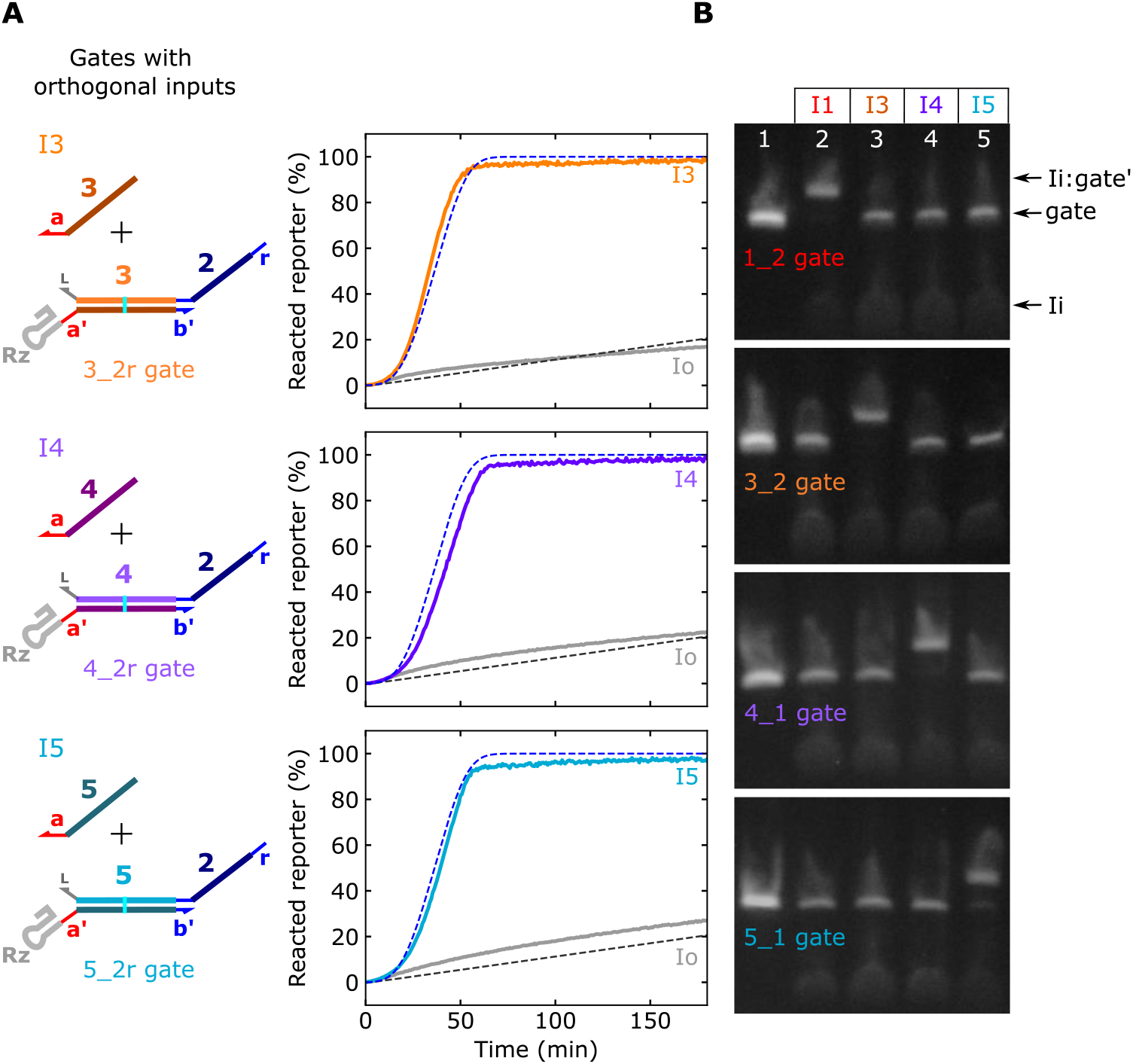
Orthogonal ctRSD input and gate sequences. (**A**) Fluorescent DNA reporter signal during co-transcription of 25 nmol/L gates with orthogonal input domains and 50 nmol/L of the designed input template or the Io template. The dashed lines show the results of the model for the 1_2r gate from Fig. 2D. (**B**) Native gel electrophoresis results demonstrating orthogonality of the four gate and input sequences. In each gel, 25 nmol/L of a single ctRSD gate was co-transcribed with no input (lane 1) or 50 nmol/L of the (I1, I3, I4, or I5) template. Electrophoresis was conducted 2 h after degradation of DNA templates with DNase I. The 1_2 gate and 3_2 gate samples were analyzed on the same gel. The 4_1 gate and 5_1 gate samples were analyzed on the same gel. Both gel images were taken with the same setting and were otherwise unmodified. See Supplementary Section 1 for schematics with sequences.

### ctRSD logic and signal amplification elements

We next investigated whether ctRSD components could be programed to execute logic (*5, 48*), signal amplification (*6, 8*), and multi-layer cascades (*7*). To assess the predictability of ctRSD circuit design, for each circuit we built we evaluated how well our kinetic model predicted behavior. Our model assumes all ctRSD components are transcribed at the same rate and all gates cleave at the same rate. Further, we assume ctRSD components with the same toehold sequence have the same strand displacement rate constants (Supplementary Section 4).

We began by designing OR and AND logic elements. The OR element was composed of two gates that react with different inputs but release the same output (Fig. 4A). We confirmed OR functionality with native gel electrophoresis (Fig. 4B) and the DNA reporter assay (Fig. 4C). Importantly, OR element kinetics closely matched model predictions (Fig. 4C). The AND element was a gate composed of two input domains separated by an internal loop (Fig. 4D). In this design, I3 reacts with the gate to expose the toehold for I1 in the internal loop. We tested AND gates with internal loops composed of (3, 4, 5, or 6) bases of the a’-toehold. The 5 base and 6 base variants resulted in complete gate reaction with 2x input template (Supplementary Fig. 21). To reduce the chance of the gate reacting with I1 alone, we chose the 5 base internal loop design. Native gel electrophoresis confirmed the AND gate reacted with I3 and I3+I1 but not with I1 alone (Fig. 4E). Similar results were observed with the DNA reporter assay, and the kinetics of output release aligned with model predictions (Fig. 4F). A second AND gate with I4 and I5 as inputs behaved similarly (Supplementary Fig. 22). Our simulations suggested the AND gates exhibited 6 % leak transcription compared to 3 % for the single input gates. This could be because AND gates are twice the length of single gates (Supplementary Section 4.1).

**Fig. 4.**
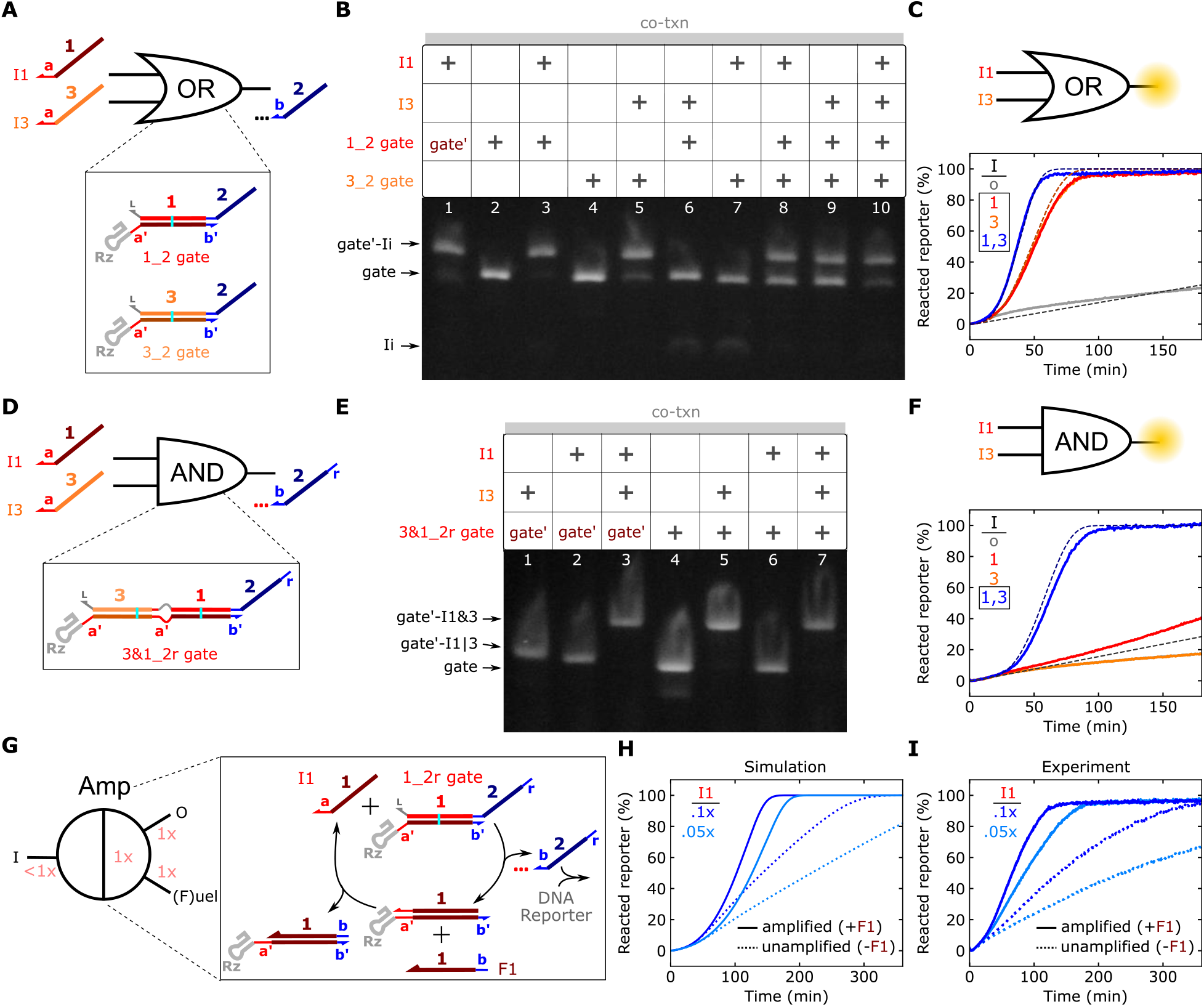
Characterization of ctRSD logic and catalytic amplification elements. (**A**) A ctRSD OR circuit element. (**B**) Native gel electrophoresis results for the OR element. Electrophoresis was conducted 0.5 h after DNase I addition. The gate’ strand is from the 1_2 gate. (**C**) Experimental (solid lines) and simulated (dashed lines) reporter signal during co-transcription of the OR element with different inputs. The trajectories for I1 alone and I3 alone overlap. The 1_2r and 3_2r gates were used in this experiment. (**D**) A ctRSD AND circuit element. See Supplementary Section 1 for schematics with sequences. (**E**) Native RNA gel electrophoresis results for the AND element. Electrophoresis was conducted 1 h after DNase I addition. The gate’ is from the 3&1_2r gate. (**F**) Experimental (solid lines) and simulated (dashed lines) DNA reporter signal during co-transcription of the AND element with different inputs. The trajectories for Io alone and I3 alone overlap. (**G**) A ctRSD catalytic amplification element. (**H** and **I**) Simulated (H) and experimental (I) DNA reporter signal during co-transcription of the 1_2r gate and I1 with (solid lines) and without (dashed lines) the F1 template (1x). For the gel results, gate and input templates were 25 nmol/L and 50 nmol/L, respectively. DNA template and T7 RNAP concentrations are tabulated in Supplement Table 4.

A powerful component in strand displacement circuits is the seesaw element, which facilitates signal amplification in larger circuits (*4, 5*). In a seesaw element, a single-stranded fuel component reacts with a I:gate’ complex to displace the input, thus allowing multiple rounds of catalytic signal release (Fig. 4G). In DNA-based circuits, which have fixed gate and input concentrations, a seesaw element enables a gate to react completely even when the input is at a lower concentration than the gate. In ctRSD, output release will eventually saturate the DNA reporter signal regardless of the input concentration. However, simulations indicated a seesaw element should decrease the time required to reach reporter saturation for low input template concentrations (Fig. 4H). When the input template was 0.05x or 0.1x the concentration of the gate template, inclusion of the fuel strand template (amplified, Fig. 4I) reduced the time to reach reporter saturation ≈3-fold and ≈4-fold, respectively, compared to samples with the input template but without the fuel template (unamplified, Fig. 4I).

### Multi-layer ctRSD cascades

Strand displacement circuits capable of complex digital logic (*5*), pattern recognition (*4*), or temporal signal release (*7*) require cascades of multi-layer signal transduction, so we next investigated whether we could program ctRSD cascades. We began by designing circuits with one to four ctRSD reaction layers in which the input and gate of the highest layer produce an output that triggers the next layer until the reporting reaction is triggered (Fig. 5A). All four multi-layer cascades exhibited kinetics in good agreement with model predictions (Fig. 5B). We next integrated ctRSD logic elements into a four-input OR circuit (Fig. 5C), a cascade of two AND gates (Fig. 5D), and two permutations of AND+OR cascades (Fig. 5, E and F). These cascades successfully executed the designed logic operations, and the experimental kinetics generally agreed with model predictions. However, there were two minor deviations in experimental kinetics compared to model predictions.

**Fig. 5.**
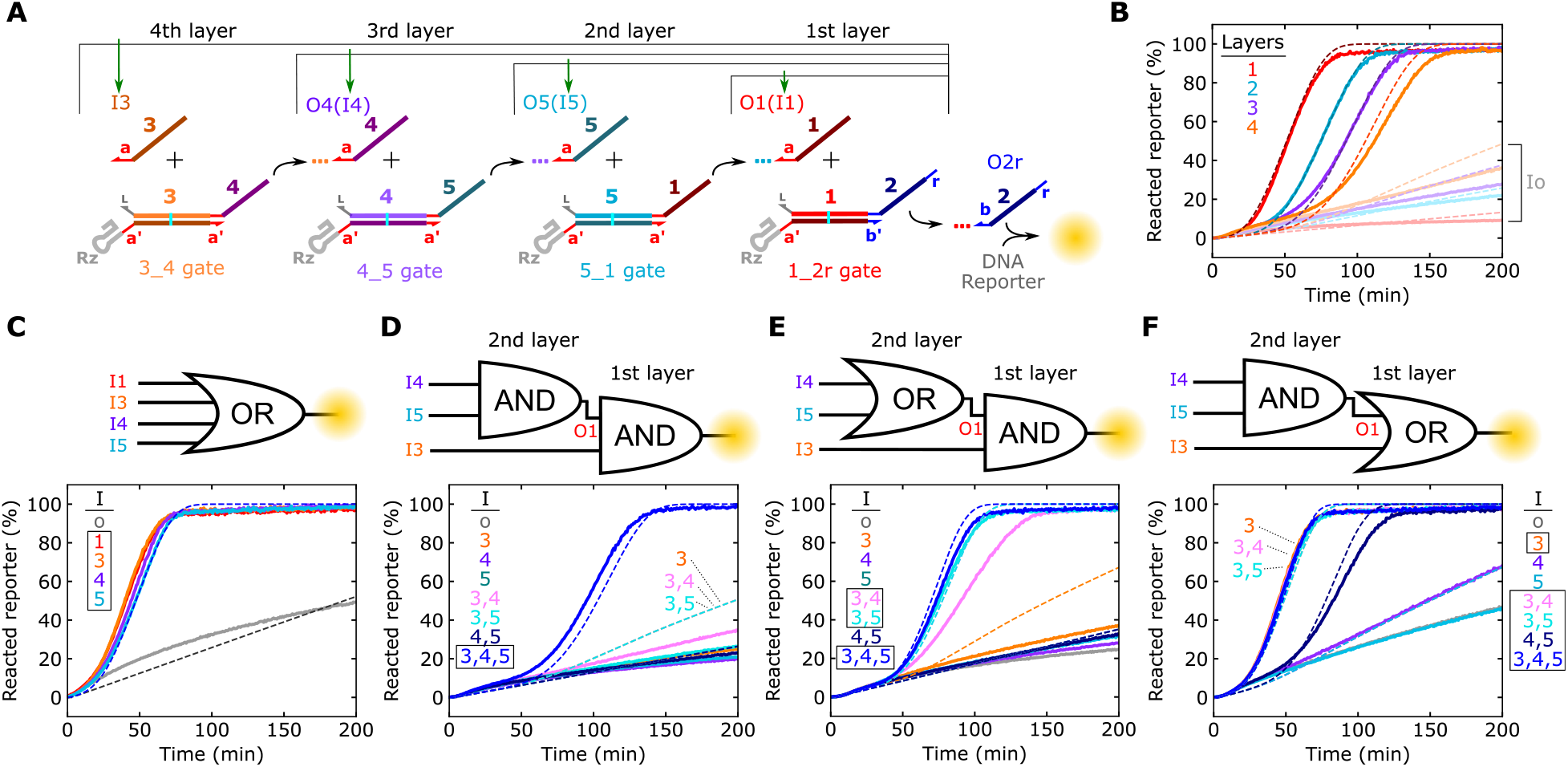
Characterization of ctRSD cascades. (**A**) Schematic of one-to four-layer cascades. Green arrows indicate the sole input template included for each cascade layer. The colored dotted lines trailing outputs represent the upstream portion of the output strand not involved in downstream reactions. (**B**) Experimental (solid lines) and simulated (dashed lines) DNA reporter signal for each layered cascade in (A). Faded lines represent each cascade with the Io template rather than the correct input template. (**C** to **F**) Experimental (solid lines) and simulated (dashed lines) reporter signal for each of the logic circuits depicted above the plots. Boxes in (C) to (F) denote the sets of inputs that should result in output release. Overlapping kinetic trajectories are labeled in the plots. In (F), the simulation results for Io, I4, and I5 all overlap with the experimental results for I4. DNA template and T7 RNAP concentrations are tabulated in Supplement Table 4.

In the first deviation from the model, the two cascades in which the first layer was the 3&1_2r gate exhibited less leak than predicted when only I3 was present (Fig. 5, D and E). I3 opens the 3&1_2r gate to react with any leak products from the upstream layer in the cascades. Presumably, the 3&1_2r gate and upstream leak products reacted less than anticipated. Our model assumes leak products react with the same rate constant as their corresponding output products, but leak products are likely misfolded gates that are bulkier than single-stranded outputs. For a I3:3&1_2r complex, the region upstream of the toehold the leak product reacts with is a duplex. Thus, steric hinderance between the I3:3&1_2r complex and a leak product could result in lower leak than predicted in simulations (Supplementary Fig. 23). Similar steric hinderance between the ctRSD gate ribozyme and an upstream leak product could explain why the observed leak in multi-layer ctRSD cascades was less than predicted (Supplementary Fig. 23 and Fig. 5B). In support of this hypothesis, we found the rate constant for a ctRSD reaction using an input with a hairpin directly adjacent to its toehold was nearly 100-fold lower than with a single-stranded input (Supplementary Fig. 23).

In the second deviation from the model, the I3+I4 reaction in the OR+AND cascade (Fig. 5E) was slower than predicted. This could be due to a slower ctRSD reaction for the 4_1 gate. The 4_1 gate itself appears to fold, cleave, and react with I4 similarly to other gates (Fig. 3B and Supplementary Fig. 20), so the difference in kinetics is not due to the gate misfolding. While all gates reuse the same toehold sequence, the kinetics of the branch migration process can vary over an order of magnitude depending on the sequence (*49*). The initial branch migration sequence of the 4_1 gate contains a weak UA tract (Supplementary Fig. 3) that could result in slower strand displacement kinetics (*49*). This mechanism is consistent with the 4_2r gate reaction being slower than gate reactions with the other three input sequences (Fig. 3B) and the four-layer ctRSD cascade being slower than predicted (Fig. 5B). Consistent with this hypothesis, reducing the I4 RNA strand displacement rate constant 2.5-fold aligned the model predictions more closely to experimental results (Supplementary Fig. 24). Although these hypotheses regarding model deviations are plausible, we present analyses using the model that assumes uniform gate performance.

### Varying ctRSD toehold lengths

In toehold-mediated strand displacement, kinetics can be precisely controlled by varying toehold length and sequence (*12*). Such kinetic control has been demonstrated for both DNA (*12*) and RNA strand displacement (*38, 39*). In the ctRSD platform, toehold length could also influence gate folding or ribozyme cleavage kinetics. Further, in our gate designs, the bulky ribozyme is directly adjacent to the toehold and could sterically hinder input binding. Thus, extending the gate toehold alone could influence kinetics by introducing a single-stranded spacer between the ribozyme and the sequence the input binds.

To explore the influence of toehold length on ctRSD, we analyzed 1_2r gates with (6, 8, 10, or 12) base toeholds. These gates cleaved with similar efficiency (Supplementary Fig. 25) and exhibited similar leak in the DNA reporter assay (Supplementary Fig. 26), indicating proper folding and cleavage. To explore the influence of toehold length and spacer length on kinetics, we designed I1 variants possessing (4, 6, 8, or 10) base toeholds and combinatorially transcribed each input alongside a 1_2r gate possessing either a (6, 8, 10, or 12) base toehold (Supplementary Fig. 27). Increasing toehold length without spacers increased the rate of ctRSD. Inclusion of spacers adjacent to the ribozyme increased ctRSD kinetics for inputs with (4, 6, or 8) base toeholds. With sufficiently long spacers the reaction rate constants for all input toehold lengths aligned with predictions from DNA-based circuits (*12*), and (6, 8, or 10) base input toehold rate constants approached the theoretical maximum (Supplementary Section 7). Thus, ctRSD kinetics can be tuned by changing toehold length, and a single-stranded spacer between the ribozyme and the toehold should be included to reduce steric hinderance.

## Discussion

Here, we developed scalable co-transcriptional RNA strand displacement circuits that were rationally programmed to execute logic, signal amplification, and multi-layer cascades. Integral to the development of ctRSD was encoding RNA gates that co-transcriptionally folded into kinetically trapped intermediates, allowing all circuit components to be produced where they execute computations. We demonstrated the scalability and modularity of ctRSD by implementing 11 single input gates and 2 AND gates in 8 different circuit topologies, all of which exhibited kinetics in agreement with our model that assumed uniform kinetic parameters. Taken together, these results indicate the robustness of our ctRSD gate design choices. Although other designs were not investigated experimentally, we believe three design choices contributed to the scalability and modularity of ctRSD: 1) selecting the stable and cleavage sequence agnostic HDV ribozyme, 2) restricting the input and output sequences to C, A, or U bases, and 3) transcribing the output strand of the gates first. These choices likely reduced the chances of misfolding during transcription and facilitated proper ribozyme function across gate sequences.

We implemented the ctRSD gates with the same modular toehold exchange design (Fig. 1, A and B) and C, A, U sequence constraints employed in state-of-the-art DNA-based circuits. In DNA computing, these designs have enabled circuits composed of >100 components that execute complex digital (*5*) and neural network (*4*) computations. Thus, ctRSD is poised to achieve the same scalability and functionality as the most advanced DNA-based TMSD circuits, while offering improved component purity and stability at comparable costs (Supplementary Section 8).

Our design choices also introduce practical limitations. The C, A, U sequence constraint restricts the use of cellular RNAs composed of all four bases as inputs. Simply redesigning gates with a four letter code could make it difficult to predictably design sequences that fold correctly (*50*). To address this limitation, we envision building upstream ctRSD translation gates that modularly convert RNA inputs with a four-letter code into outputs with a three-letter code that are processed in ctRSD circuits with our prescribed design rules. In this manner, the same robust information processing circuits may be used, and translation gates with four-letter codes that function correctly could be identified by testing sequences spanning a cellular RNA of interest.

Another limitation of our design is the bulky HDV ribozyme motif left on the gates after cleavage. We found this motif influenced ctRSD kinetics unless a single-stranded spacer between the ribozyme and the toehold binding sequence was inserted (Supplementary Section 7). Recently, a scheme was reported for transcriptionally encoding strand displacement circuits that used a dual hammerhead ribozyme motif that excised itself after folding (*29*), and a similar multi-ribozyme strategy could be applied to ctRSD gates to remove the HDV ribozyme motif during gate production. However, in contrast to the ctRSD circuits presented here, gate performance in this alternative scheme varied with sequence and toeholds switched from 5’ to 3’ between circuit layers, reducing modularity and composability. Ultimately, merging ideas from both these implementations offers routes for further optimizing ctRSD.

We envision ctRSD enabling many new applications in nucleic acid computing and synthetic biology. For example, the inclusion of RNases in the ctRSD platform would allow continuous circuit turnover. Circuits could then respond multiple times to changing input signals, overcoming a current limitation in DNA computing (*11, 19*). Additionally, regulating input production with allosteric transcription factors allows ctRSD circuits that process non-nucleic acid inputs to be readily developed for smart diagnostics (*30, 31*). Finally, the ability to transcriptionally encode strand displacement components in DNA plasmids allows nucleic acid computing to be employed in a number of new environments where DNA computing is limited due to degradation (*16*), *e.g*. in blood samples (*17*), cell-free lysates, or inside living cells (*18*). *In vivo*, fluorescent RNA aptamers (*46, 47*) or RNA regulators that transduce RNA signals into fluorescent protein production (*24*) could track ctRSD dynamics. Further, ctRSD outputs could regulate protein expression through existing RNA technologies (*22, 23, 27*), allowing ctRSD circuits to control cellular function. Together, these examples indicate the potential for ctRSD to serve as a versatile, enabling technology across many synthetic biology platforms.

## Materials and Methods

### DNA and materials

DNA transcription templates were ordered as gBlock gene fragments from Integrated DNA Technologies (IDT), amplified *via* polymerase chain reaction (PCR) with Phusion High-Fidelity PCR Master Mix (Cat #: F531L) from ThermoFisher Scientific, and purified using Qiagen PCR clean-up kits. All DNA oligo primers were ordered from IDT with standard desalting. For *in vitro* transcription experiments T7 RNA polymerase (RNAP) and ribonucleotide triphosphates (NTPs) were ordered from ThermoFisher Scientific (Cat #: R0481). DNase I (Cat #: M0303S) was purchased from New England Biolabs (NEB). 4 % agarose EX E-gels were purchased from ThermoFisher Scientific (Cat #: G401004). All chemicals were purchased from Sigma Aldrich.

### Transcription template preparation

All transcription templates were prepared by PCR of 0.2 ng of gBlock DNA with Phusion High-Fidelity PCR Master Mix and 0.5 μmol/L of forward and reverse primers. PCR was conducted for 30 cycles with a 30 s 98 °C denaturing step, a 30 s 60 °C primer annealing step, and a 30 s 72 °C extension step. A 3 min 72 °C final extension step was executed at the end of the program. Following PCR amplification, the samples were purified with Qiagen PCR clean-up kits and eluted in Qiagen Buffer EB (10 mmol/L Tris-HCl, pH 8.5).

### RNA agarose gel electrophoresis

4 % agarose EX E-gels were used for all RNA gel electrophoresis experiments. These gels are pre-stained with SYBR Gold for fluorescence imaging. Electrophoresis was conducted on a Egel powerbase, and all E-gels were imaged using the E-gel power snap camera (ThermoFisher Scientific, Cat #: G8200). Unless otherwise stated, to prepare RNA for gel electrophoresis, DNA templates were transcribed at 37 °C for 30 min in transcription conditions (see *Characterization of RNA strand displacement with in vitro transcription*) with 0.6 U/μL T7 RNAP. To stop transcription, CaCl2 (final concentration (1 to 1.5) mmol/L) and DNase I (final concentration (0.1 to 0.2) U/μL) were added to degrade the DNA templates. After DNase I addition, the samples were left at 37 °C for (0.5 to 2) h (see Figure captions), and subsequently analyzed with gel electrophoresis. For native gels, the gels were sandwiched between icepacks to keep the gels cool during electrophoresis and were run for (45 to 60) min prior to imaging. Integrated band intensities were quantified in gel images using the Gel Analysis Tool in ImageJ as previously described (*51*). For denaturing gels, prior to electrophoresis, a solution of 100 % formamide, 36 mmol/L EDTA was mixed 1:1 by volume with the samples and the samples were heated to 90 °C for 5 min. The samples were then immediately loaded on gels for electrophoresis and run for (20 to 30) min before imaging. Gel images were not post processed, any brightness and contrast adjustments were executed during image acquisition and were thus applied uniformly to the images to aid visualization.

### Characterization of RNA strand displacement with a fluorescence DNA reporter

The *in vitro* transcription reactions with DNA reporter complexes were conducted in transcription buffer prepared in house (40 mmol/L Tris-HCl - pH 7.9, 6 mmol/L MgCl2, 10 mmol/L dithiothreitol (DTT), 10 mmol/L NaCl, and 10 mmol/L spermidine) supplemented with 2 mmol/L final concentration of each NTP type (ATP, UTP, CTP, GTP). All transcription reactions were conducted at 37 °C. Unless otherwise stated, 500 nmol/L of DNA reporter was used. For *in vitro* transcription reactions, all components other than T7 RNAP were mixed and tracked in the plate reader for 15 min to 60 min prior to adding T7 RNAP. Addition of T7 RNAP, followed by mixing, corresponded to *t* = 0 min in *in vitro* transcription experiments. The time to mix T7 RNAP into all samples for an experiment was less than one min. In our experiments, the T7 RNAP concentration varied depending on the total concentration of DNA templates present. To compare the response of a given ctRSD circuit to different input template concentrations or a different number of input templates, the same total template concentration was used across all reactions to ensure the same transcriptional load across samples. An input template (Io) that produces an RNA that does not interact with the gates was added to maintain the template concentration across samples. Supplementary Table 4 contains the concentrations of DNA templates (including Io) and T7 RNAP used in each experiment.

### Transcription rate calibration and sample variability

In our experiments, the transcription rate depended on the concentration of T7 RNAP and the total concentration of DNA templates (Supplementary Fig. 28). Further, variability of T7 RNAP activity (*52*) across manufacturer lots was expected to be the primary source of variation in our experiments. To calibrate for these effects, we developed a transcription rate reference sample (Supplementary Fig. 28). This reference sample tracked transcription with a template that constitutively expressed the 1_2r strand and contained the same T7 RNAP lot and concentration as the experimental samples on a given day. Additionally, the Io template was added so the total template concentration equaled that of the experimental samples. The reference sample calibrated the first order transcription rate constant chosen for simulations (Supplementary Fig. 29), thus accounting for variation in T7 RNAP activity when assessing how well experimental results agreed with model predictions. To estimate the variability in ctRSD reaction measurements introduced during sample preparation, we conducted reactions between the 1_2r gate and either I1 or Io in triplicate in the DNA reporter assay. Each reaction was prepared independently using the same transcription template, NTP, buffer, and T7 RNAP stocks. These replicates exhibited a standard deviation of < 1.5 % from the mean value at each time point (Supplementary Fig. 30). A variability of < 5 % standard deviation was observed for the AND gate cascade in Fig. 5D (Supplementary Fig. 30). Additionally, reactions between the 1_2r gate and either I1 or Io performed on different days exhibited < 3% standard deviation (Supplementary Fig. 31). We therefore assumed a conservative variability of < 5 % generalized to ctRSD circuits. For the small circuits studied here, we do not expect this level of variability to influence our conclusions and, unless otherwise stated, DNA reporter experiments were conducted with a single experimental replicate.

### Fluorescence data acquisition and normalization

BioTek Synergy Neo2 plate readers were used track *in vitro* transcription reactions. Reactions were typically conducted in 70 μL volumes in Greiner μClear 96-well plates (Cat #: 655096) read from the bottom. The DNA reporter complex was labeled with a HEX dye which was tracked with excitation: 524 nm (20 nm bandwidth), emission: 565 nm (20 nm bandwidth), and a gain of 85. Fluorescence readings were taken every 46 s. In a typical experiment, fluorescence readings were taken for (25 to 45) min before T7 RNAP was added to initiate the reactions. At the end of most experiments, an excess (2.5 μmol/L) of a DNA version of the O2r strand was added to each sample to obtain an internal maximum DNA reporter fluorescence value. Fluorescence data was then normalized as:

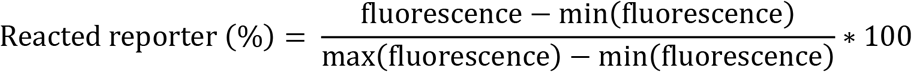

If the DNA O2r strand was not added, a control well in which the ctRSD reaction had saturated the reporter signal served as a max value for normalization.

## Supporting information

Fluorescence data

## Acknowledgments

The authors thank John T. Elliot, Julius B. Lucks, Pepijn G. Moerman, Moshe Rubanov for insightful comments on the manuscript and Nina Y. Alperovich, Jaeyoung K. Jung, Abe D. Pressman, Drew S. Tack, Jayan Rammohan, Eugenia F. Romantseva, David J. Ross, Olga B. Vasilyeva for helpful discussions throughout this work.

## Funding

National Research Council Postdoctoral Fellowship (SWS)

## Author contributions

SWS conceived, designed, and performed research. SWS conducted data analysis, developed simulation code, and executed simulations. EAS supervised the research. SWS and EAS wrote the manuscript.

## Competing interests

SWS has filed an invention disclosure pertaining to the technology presented in this manuscript. The authors declare no other competing interests.

## Data availability

The raw and normalized fluorescence data is supplied in Supplementary Data File 2. Any other relevant data are available from the authors upon reasonable request.

## Code availability

The simulation code is posted on GitHub: https://github.com/usnistgov/ctRSD-simulator

## Disclaimer

Certain commercial entities, equipment, or materials may be identified in this document to describe an experimental procedure or concept adequately. Such identification is not intended to imply recommendation or endorsement by the National Institute of Standards and Technology, nor is it intended to imply that the entities, materials, or equipment are necessarily the best available for the purpose. Official contribution of the National Institute of Standards and Technology; not subject to copyright in the United States.

## Notes

https://github.com/usnistgov/ctRSD-simulator

### Competing Interest Statement

The authors have declared no competing interest.

