## Supplementary Materials for "Co-transcriptional RNA strand displacement circuits"

Table of contents

**1. Sequences, schematics, and control transcripts ... 3**
**2. ctRSD design considerations ... 14**
2.1 Separate transcription of gate strands results in downstream leak ... 15
2.2 The co-transcriptional folding pathway ... 17
2.3 Self-cleaving ribozyme selection ... 19
**3. Equilibrium analysis with NUPACK ... 21**
**4. Modeling ctRSD reactions ... 22**
4.1 Model assumptions and reactions ... 22
4.2 Kinetic parameters used to model co-transcriptional RNA strand displacement ... 26
4.3 Modeling and characterizing leak ... 28
**5. Design of additional ctRSD sequence domains and circuit elements ... 33**
**6. Analysis of deviations between experiments and simulations ... 36**
**7. 1_2r gates with different toehold lengths ... 39**
**8. Potential advantages of ctRSD circuits compared to DNA-based circuits ... 42**
8.1 Cost analysis ... 42
**9. Experimental conditions, transcription rate calibration, and experimental variability ... 44**
9.1 Experimental conditions ... 45
9.2 Transcription rate calibration ... 47
9.3 Experimental variability ... 49

**DISCLAIMER**

Certain commercial entities, equipment, or materials may be identified in this document to describe an experimental procedure or concept adequately. Such identification is not intended to imply recommendation or endorsement by the National Institute of Standards and Technology, nor is it intended to imply that the entities, materials, or equipment are necessarily the best available for the purpose. Official contribution of the National Institute of Standards and Technology; not subject to copyright in the United States.

### 1. Sequences, schematics, and control transcripts

**Supplementary Table 1:** DNA sequences used in the study. All transcription templates were ordered as gBlock gene fragments from IDT. All primers were ordered without purification from IDT. For the input and fuel templates the last 30 lower case bases were added to bring the sequence above 125 bases to order as gBlocks. The PCR product resulting from the T7fwd and T7rev primers does not include this sequence. The T7 RNAP promoter sequence is underlined in all sequences. Black highlighted bases indicate bases that were mutated from a C to a T to render the HDV ribozyme catalytically inactive.

| Sequence name | Sequence |
|---|---|
| Inputs / Fuels | |
| Io | TTCTAATACGACTCACTATAGGGAGATTCGTCTCCCA<br>ATCAATAACACACATA<br>CTATAACCCCTTGGGGCCTCTAAACGGGTCTTGAGGGGTTTTTTG<br>ctgaaaggaggaactatatccggatatccc |
| I1 | TTCTAATACGACTCACTATAGGGAGATTCGTCTCCCA<br>TCACTTCACAACATCA<br>CTATAACCCCTTGGGGCCTCTAAACGGGTCTTGAGGGGTTTTTTG<br>ctgaaaggaggaactatatccggatatccc |
| I3 | TTCTAATACGACTCACTATAGGGAGATTCGTCTCCCA<br>CACATTCAATACATCA<br>CTATAACCCCTTGGGGCCTCTAAACGGGTCTTGAGGGGTTTTTTG<br>ctgaaaggaggaactatatccggatatccc |
| I4 | TTCTAATACGACTCACTATAGGGAGATTCGTCTCCCA<br>ACCACTAAACACATCA<br>CTATAACCCCTTGGGGCCTCTAAACGGGTCTTGAGGGGTTTTTTG<br>ctgaaaggaggaactatatccggatatccc |
| I5 | TTCTAATACGACTCACTATAGGGAGATTCGTCTCCCT<br>ACCATACTACACATCA<br>CTATAACCCCTTGGGGCCTCTAAACGGGTCTTGAGGGGTTTTTTG<br>ctgaaaggaggaactatatccggatatccc |
| F1 | TTCTAATACGACTCACTATAGGGAGATTCGTCTCCCA<br>ATTAACTCACTTCACA<br>CTTTAACCCCTTGGGGCCTCTAAACGGGTCTTGAGGGGTTTTTTG<br>ctgaaaggaggaactatatccggatatccc |
| I1 (4 base toehold) | TTCTAATACGACTCACTATAGGGAGATTCGTCTCCCA<br>TCACTTCACAACAT<br>ATATAACCCCTTGGGGCCTCTAAACGGGTCTTGAGGGGTTTTTTG<br>ctgaaaggaggaactatatccggatatccc |
| I1 (8 base toehold) | TTCTAATACGACTCACTATAGGGAGATTCGTCTCCCA<br>TCACTTCACAACATCATA<br>ATATAACCCCTTGGGGCCTCTAAACGGGTCTTGAGGGGTTTTTTG<br>ctgaaaggaggaactatatccggatatccc |
| I1 (10 base toehold) | TTCTAATACGACTCACTATAGGGAGATTCGTCTCCCA<br>TCACTTCACAACATCATACT<br>ATATAACCCCTTGGGGCCTCTAAACGGGTCTTGAGGGGTTTTTTG<br>ctgaaaggaggaactatatccggatatccc |

| Gates | |
|---|---|
| unfolded xRz gate | TTCTAATACGACTCACTATAGGGAGATTCGTCTCCCA<br>ACATTACATAATCAAATACATACATATTCGGGTCGGCA<br>TGGCATCTCCACCTCCTCGCGGTCCGACCTGGGCTACTT<br>CGGTAGGTTAAGGGAGTGATGTTGTGAAGTTAGTTAAT<br>CTATAACCCCTTGGGGCCTCTAAACGGGTCTTGAGGGGTTTTTTG |
| 1_2 xRz gate | TTCTAATACGACTCACTATAGGGAGATTCGTCTCCCA<br>CCACATACTAATTAACTTACTTCACATTCGGGTCGGCA<br>TGGCATCTCCACCTCCTCGCGGTCCGACCTGGGCTACTT<br>CGGTAGGTTAAGGGAGTGATGTTGTGAAGT**T**AGTTAAT<br>CTATAACCCCTTGGGGCCTCTAAACGGGTCTTGAGGGGTTTTTTG |
| 1_2r xRz gate | TTCTAATACGACTCACTATAGGGAGATTCGTCTCCCA<br>CTACATCCACATACTAATTAACTTACTTCACATTCGGGTCGGCA<br>TGGCATCTCCACCTCCTCGCGGTCCGACCTGGGCTACTT<br>CGGTAGGTTAAGGGAGTGATGTTGTGAAGT**T**AGTTAAT<br>CTATAACCCCTTGGGGCCTCTAAACGGGTCTTGAGGGGTTTTTTG |
| 1_2 HDV cau | TTCTAATACGACTCACTATAGGGAGATTCGTCTCCCA<br>CCACATACTAATTAACTTACTTCACATTCGGGTCGGCA<br>TGGCATCTCCACCTCCTCGCGGTCCGACCTGGGCTACTT<br>CGGTAGGCTAAGGGAGATACACTCACTATCACTACTCAT<br>TCATAACCCCTTGGGGCCTCTAAACGGGTCTTGAGGGGTTTTTTG |
| 1_2r HDV cau | TTCTAATACGACTCACTATAGGGAGATTCGTCTCCCA<br>CTACATCCACATACTAATTAACTTACTTCACATTCGGGTCGGCA<br>TGGCATCTCCACCTCCTCGCGGTCCGACCTGGGCTACTT<br>CGGTAGGCTAAGGGAGATACACTCACTATCACTACTCAT<br>TCATAACCCCTTGGGGCCTCTAAACGGGTCTTGAGGGGTTTTTTG |
| 1_2 gate HP Rz | TTCTAATACGACTCACTATAGGGAGATTCGTCTCCCA<br>CCACATACTAATTAACTTACTTCACATACCCTCACAGT<br>CCAGTGATTCGTCACTGAGAAGTGAACCAGAGAAACACA<br>CGTTGTGGTATATTACCTGGTCTGATGTTGTGAAGT**G**AGTTAAT<br>CTATAACCCCTTGGGGCCTCTAAACGGGTCTTGAGGGGTTTTTTG |
| gate' | TTCTAATACGACTCACTATAGGGAGATTCGTCTCCCA<br>TTCGGGTCGGCATGGCATCTCCACCTCCTCGCGGTCCGACCTGGGCTACTT<br>CGGTAGGCTAAGGGAGTGATGTTGTGAAGT**G**AGTTAAT<br>CTATAACCCCTTGGGGCCTCTAAACGGGTCTTGAGGGGTTTTTTG |
| 1_2 gate | TTCTAATACGACTCACTATAGGGAGATTCGTCTCCCA<br>CCACATACTAATTAACTTACTTCACATTCGGGTCGGCA<br>TGGCATCTCCACCTCCTCGCGGTCCGACCTGGGCTACTT<br>CGGTAGGCTAAGGGAGTGATGTTGTGAAGT**G**AGTTAAT<br>CTATAACCCCTTGGGGCCTCTAAACGGGTCTTGAGGGGTTTTTTG |
| 1_2r gate | TTCTAATACGACTCACTATAGGGAGATTCGTCTCCCA<br>CTACATCCACATACTAATTAACTTACTTCACATTCGGGTCGGCA<br>TGGCATCTCCACCTCCTCGCGGTCCGACCTGGGCTACTT<br>CGGTAGGCTAAGGGAGTGATGTTGTGAAGT**G**AGTTAAT<br>CTATAACCCCTTGGGGCCTCTAAACGGGTCTTGAGGGGTTTTTTG |
| 3_2 gate | TTCTAATACGACTCACTATAGGGAGATTCGTCTCCCA<br>CCACATACTAATTAACCATATTCAATTTCGGGTCGGCA<br>TGGCATCTCCACCTCCTCGCGGTCCGACCTGGGCTACTT<br>CGGTAGGCTAAGGGAGTGATGTATTGAAT**G**TGGTTAAT<br>CTATAACCCCTTGGGGCCTCTAAACGGGTCTTGAGGGGTTTTTTG |

| | |
|---|---|
| 3_2r gate | TTCTAATACGACTCACTATAGGGAGATTCGTCTCCCA<br>CTACATCCACATACTAATTAACCATATTCAATTTCGGGTCGGCA<br>TGGCATCTCCACCTCCTCGCGGTCCGACCTGGGCTACTT<br>CGGTAGGCTAAGGGAGTGATGTATTGAAT**G**TGGTTAAT<br>CTATAACCCCTTGGGGCCTCTAAACGGGTCTTGAGGGGTTTTTTG |
| 4_2r gate | TTCTAATACGACTCACTATAGGGAGATTCGTCTCCCA<br>CTACATCCACATACTAATTAACACTACTAAACTTC<br>GGGTCGGCATGGCATCTCCACCTCCTCGCGGT<br>CCGACCTGGGCTACTTCGGTAGGCTAAGGGAG<br>TGATGTGTTTAGTGGTGTTAAT<br>CTATAACCCCTTGGGGCCTCTAAACGGGTCTTGAGGGGTTTTTTG |
| 5_2r gate | TTCTAATACGACTCACTATAGGGAGATTCGTCTCCCA<br>CTACATCCACATACTAATTAACACTATACTACTTC<br>GGGTCGGCATGGCATCTCCACCTCCTCGCGGT<br>CCGACCTGGGCTACTTCGGTAGGCTAAGGGAG<br>TGATGTGTAGTATGGTGTTAAT<br>CTATAACCCCTTGGGGCCTCTAAACGGGTCTTGAGGGGTTTTTTG |
| 4_1 gate | TTCTAATACGACTCACTATAGGGAGATTCGTCTCCCA<br>TCACTTCACAACATCAACTACTAAACTTC<br>GGGTCGGCATGGCATCTCCACCTCCTCGCGGT<br>CCGACCTGGGCTACTTCGGTAGGCTAAGGGAG<br>TGATGTGTTTAGTGGTTGATGT<br>CTATAACCCCTTGGGGCCTCTAAACGGGTCTTGAGGGGTTTTTTG |
| 5_1 gate | TTCTAATACGACTCACTATAGGGAGATTCGTCTCCCA<br>TCACTTCACAACATCAACTATACTACTTC<br>GGGTCGGCATGGCATCTCCACCTCCTCGCGGT<br>CCGACCTGGGCTACTTCGGTAGGCTAAGGGAG<br>TGATGTGTAGTATGGTTGATGT<br>CTATAACCCCTTGGGGCCTCTAAACGGGTCTTGAGGGGTTTTTTG |
| 4_5 gate | TTCTAATACGACTCACTATAGGGAGATTCGTCTCCCA<br>ACCATACTACACATCAACTACTAAACTTC<br>GGGTCGGCATGGCATCTCCACCTCCTCGCGGT<br>CCGACCTGGGCTACTTCGGTAGGCTAAGGGAG<br>TGATGTGTTTAGTGGTTGATGT<br>AATTAACCCCTTGGGGCCTCTAAACGGGTCTTGAGGGGTTTTTTG |
| 3_4 gate | TTCTAATACGACTCACTATAGGGAGATTCGTCTCCCA<br>ACCACTAAACACATCACATATTCAATTTC<br>GGGTCGGCATGGCATCTCCACCTCCTCGCGGT<br>CCGACCTGGGCTACTTCGGTAGGCTAAGGGAG<br>TGATGTATTGAAT**G**TGTGATGT<br>AATTAACCCCTTGGGGCCTCTAAACGGGTCTTGAGGGGTTTTTTG |
| 3&1_2r gate | TTCTAATACGACTCACTATAGGGAGATTCGTCTCCCA<br>CTACATCCACATACTAATTAACTTACTTCACATAATACATTCAATTTC<br>GGGTCGGCATGGCATCTCCACCTCCTCGCGGT<br>CCGACCTGGGCTACTTCGGTAGGCTAAGGGAG<br>TGATGTATTGAATGTGTGATGTTGTGAAGTGAGTTAAT<br>CTATAACCCCTTGGGGCCTCTAAACGGGTCTTGAGGGGTTTTTTG |
| gate' (3&1_2) | TTCTAATACGACTCACTATAGGGAGATTCGTCTCCCA<br>TTCGGGTCGGCATGGCATCTCCACCTCCTCGCGGT<br>CCGACCTGGGCTACTTCGGTAGGCTAAGGGAG<br>TGATGTATTGAATGTGTGATGTTGTGAAGTGAGTTAAT<br>CTATAACCCCTTGGGGCCTCTAAACGGGTCTTGAGGGGTTTTTTG |

| | |
|---|---|
| 5&4_1 gate | TTCTAATACGACTCACTATAGGGAGATTCGTCTCCCA<br>TCACTTCACAACATCAACTACTAAACTAAACTATACTACTTC<br>GGGTCGGCATGGCATCTCCACCTCCTCGCGGT<br>CCGACCTGGGCTACTTCGGTAGGCTAAGGGAG<br>TGATGTGTAGTATGGTTGATGTGTTTAGTGGTTGATGT<br>CTATAACCCCTTGGGGCCTCTAAACGGGTCTTGAGGGGTTTTTTG |
| 5&4_2r gate | TTCTAATACGACTCACTATAGGGAGATTCGTCTCCCA<br>CTACATCCACATACTAATTAACACTACTAAACTAAACTATACTACTTC<br>GGGTCGGCATGGCATCTCCACCTCCTCGCGGT<br>CCGACCTGGGCTACTTCGGTAGGCTAAGGGAG<br>TGATGTGTAGTATGGTTGATGTGTTTAGTGGTGTTAAT<br>CTATAACCCCTTGGGGCCTCTAAACGGGTCTTGAGGGGTTTTTTG |
| 3&1_2 (3bp a'-loop) | TTCTAATACGACTCACTATAGGGAGATTCGTCTCCCA<br>CCACATACTAATTAACTTACTTCACATTCACATATTCAATTTC<br>GGGTCGGCATGGCATCTCCACCTCCTCGCGGT<br>CCGACCTGGGCTACTTCGGTAGGCTAAGGGAG<br>TGATGTATTGAATGTGTGATGTTGTGAAGTGAGTTAAT<br>CTATAACCCCTTGGGGCCTCTAAACGGGTCTTGAGGGGTTTTTTG |
| 3&1_2 (4bp a'-loop) | TTCTAATACGACTCACTATAGGGAGATTCGTCTCCCA<br>CCACATACTAATTAACTTACTTCACACCATACATTCAATTTC<br>GGGTCGGCATGGCATCTCCACCTCCTCGCGGT<br>CCGACCTGGGCTACTTCGGTAGGCTAAGGGAG<br>TGATGTATTGAATGTGTGATGTTGTGAAGTGAGTTAAT<br>CTATAACCCCTTGGGGCCTCTAAACGGGTCTTGAGGGGTTTTTTG |
| 3&1_2 (5bp a'-loop) | TTCTAATACGACTCACTATAGGGAGATTCGTCTCCCA<br>CCACATACTAATTAACTTACTTCACATAATACATTCAATTTC<br>GGGTCGGCATGGCATCTCCACCTCCTCGCGGT<br>CCGACCTGGGCTACTTCGGTAGGCTAAGGGAG<br>TGATGTATTGAATGTGTGATGTTGTGAAGTGAGTTAAT<br>CTATAACCCCTTGGGGCCTCTAAACGGGTCTTGAGGGGTTTTTTG |
| 3&1_2 (6bp a'-loop) | TTCTAATACGACTCACTATAGGGAGATTCGTCTCCCA<br>CCACATACTAATTAACTTACTTCACACCATATTCAATTTC<br>GGGTCGGCATGGCATCTCCACCTCCTCGCGGT<br>CCGACCTGGGCTACTTCGGTAGGCTAAGGGAG<br>TGATGTATTGAATGTGTGATGTTGTGAAGTGAGTTAAT<br>CTATAACCCCTTGGGGCCTCTAAACGGGTCTTGAGGGGTTTTTTG |
| 5_1 HDV cau | TTCTAATACGACTCACTATAGGGAGATTCGTCTCCCA<br>TCACTTCACAACATCAACTATACTACTTC<br>GGGTCGGCATGGCATCTCCACCTCCTCGCGGT<br>CCGACCTGGGCTACTTCGGTAGGCTAAGGGAG<br>TACACTCACTATCACTACTCATT<br>CATAACCCCTTGGGGCCTCTAAACGGGTCTTGAGGGGTTTTTTG |
| 5_1 leak hairpin | TTCTAATACGACTCACTATAGGGAGATTCGTCTCCCA<br>TCACTTCACAACATCAACTATACTACTTC<br>GGGTCGGCATGGCATCTCCACCTCCTCGCGGT<br>CCGACCTGGGCTACTTCGGTAGGTTAAGGGAG<br>TGATGTGTAGTATGGT<br>CTATAACCCCTTGGGGCCTCTAAACGGGTCTTGAGGGGTTTTTTG |
| 1_2r gate<br>(8 base toehold) | TTCTAATACGACTCACTATAGGGAGATTCGTCTCCCA<br>CTACATCCACATACTAATTAACTTACTTCACATTC<br>GGGTCGGCATGGCATCTCCACCTCCTCGCGGT<br>CCGACCTGGGCTACTTCGGTAGGCTAAGGGAG<br>TATGATGTTGTGAAGTGAGTTAAT<br>CTATAACCCCTTGGGGCCTCTAAACGGGTCTTGAGGGGTTTTTTG |

| | |
|---|---|
| 1_2r gate (10 base toehold) | TTCTAATACGACTCACTATAGGGAGATTCGTCTCCCA CTACATCCACATACTAATTAACTTACTTCACATTC GGGTCGGCATGGCATCTCCACCTCCTCGCGGT CCGACCTGGGCTACTTCGGTAGGCTAAGGGAG AGTATGATGTTGTGAAGTGAGTTAAT CTATAACCCCTTGGGGCCTCTAAACGGGTCTTGAGGGGTTTTTTG |
| 1_2r gate (12 base toehold) | TTCTAATACGACTCACTATAGGGAGATTCGTCTCCCA CTACATCCACATACTAATTAACTTACTTCACATTC GGGTCGGCATGGCATCTCCACCTCCTCGCGGT CCGACCTGGGCTACTTCGGTAGGCTAAGGGAG AGAGTATGATGTTGTGAAGTGAGTTAAT CTATAACCCCTTGGGGCCTCTAAACGGGTCTTGAGGGGTTTTTTG |
| **DNA oligos** | |
| T7fwd | TTCTAATACGACTCACTATAGGGAG |
| T7rev | CAAAAAACCCCTCAAGACCCGTTTAG |
| O2-nt | TTCTAATACGACTCACTATAGGGAGATTCGTCTCCCA CCACATACTAATTAACTTACTTCACATTC |
| O2-t | GAATGTGAAGTAAGTTAATTAGTATGTGGT GGGAGACGAATCTCCCTATAGTGAGTCGTATTAGAA |
| O2r-nt | TTCTAATACGACTCACTATAGGGAGATTCGTCTCCCA CTACATCCACATACTAATTAACTTACTTCACATTC |
| O2r-t | GAATGTGAAGTAAGTTAATTAGTATGTGGATGTAGT GGGAGACGAATCTCCCTATAGTGAGTCGTATTAGAA |
| DNA reporter F | /5HEX/CTACATCCACATACTA |
| DNA reporter Q | GTTAATTAGTATGTGGATGTAG/3IABRQSp/ |

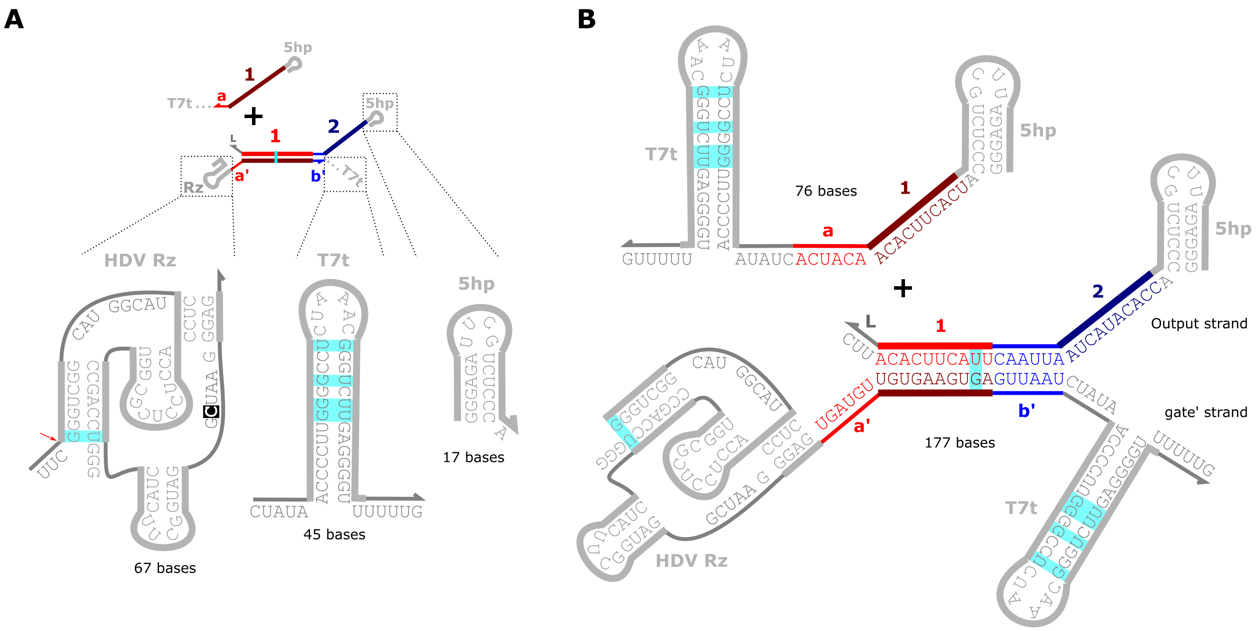


**Supplementary Figure 1:** RNA sequence motifs used to co-transcriptionally encode stand displacement circuit components. (**A**) The HDV ribozyme (Rz), an internal self-cleaving ribozyme, was added between the output and gate' strands of ctRSD gates. The red arrow adjacent to the HDV ribozyme indicates the cleavage site. The cytosine highlighted in black is required for catalytic activity and was mutated to uracil in inactive mutants used in this study. Additionally, a T7 RNAP terminator (T7t) sequence was appended to the 3' end of the gate' strand of ctRSD gates and to the 3' end of input sequences. Further, all transcripts were designed with a hairpin motif at their 5' end (5hp). This motif possesses the consensus initiation sequence (5′GGGAGA) for T7 RNAP (*1*). Finally, a G-U wobble base pair was introduced within the double stranded branch migration domain (cyan) to reduce secondary structure in the DNA template, which can interfere with synthesis (*2*), and to provide a thermodynamic driving force for strand displacement with inputs that convert the G-U wobble pair to a G-C pair (*3*). G-U wobble base pairs are also present in the HDV Rz and T7t sequences. (**B**) The full sequence schematics of the I1 and the 1_2 gate transcripts.

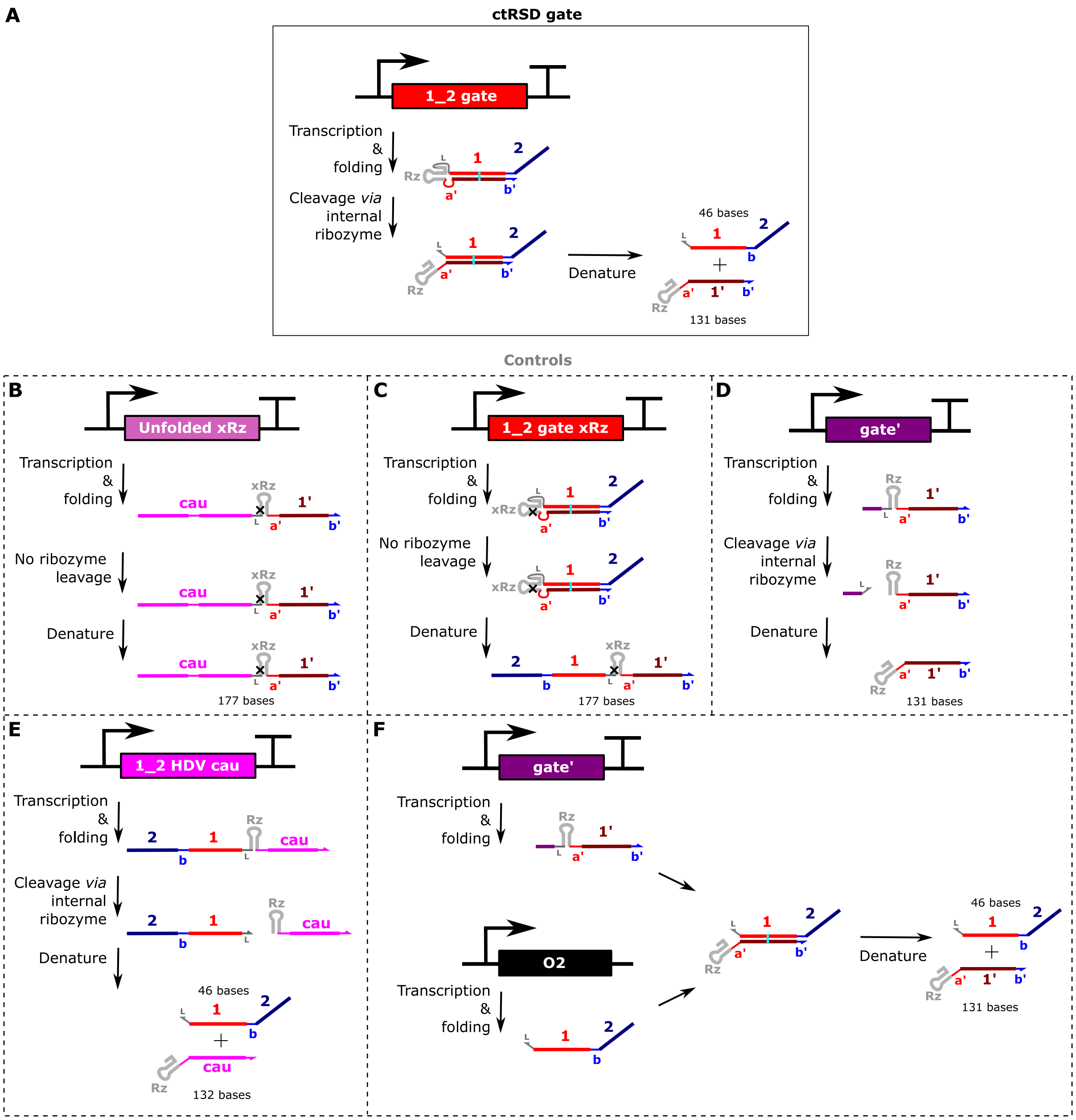


**Supplementary Figure 2:** Schematics of control RNA transcripts used throughout the study. A black X in the Rz domains indicates a C → U mutation at the 75$^{th}$ position of the genomic HDV ribozyme that prevents cleavage from occurring (xRz) (*4*). Pink cau domains are designed without guanine bases in their sequence. (**A**) Schematic of the 1_2 ctRSD gate investigated in this study. (**B**) A control transcript corresponding to the full length of the desired 1_2 gate transcript. In this control, an inactive HDV mutant was used to prevent cleavage, and the 1-, b-, and 2-domains were mutated to prevent the transcript from folding into a hairpin. (**C**) A control transcript that is the 1_2 gate with an inactive HDV mutant to prevent self-cleavage. In denaturing conditions, this transcript should be the same length as the control transcript in (B). (**D**) A control transcript corresponding to the gate' strand of the 1_2 gate. (**E**) A control transcript possessing the output sequence of the 1_2 gate, followed by the HDV ribozyme and 22 C, A, or U bases. This control produces an HDV ribozyme cleaved output strand identical to the output strand the 1_2 gate in (A). The product of HDV ribozyme cleavage (Rz- and cau-domains) should also migrate similarly to the control transcript in (D) in agarose gel electrophoresis experiments. (**F**) A control in which the two strands of the 1_2 gate are produced from separate transcription templates. The O2 transcription template was prepared from two oligos (O2-nt and O2-t in Supplementary Table 1) that were mixed in a 1:1 molar ratio in transcription buffer, heated to 90 °C for 5 min, and then cooled to 20 °C at a rate of 1 °C/min. To obtain a 1:1 concentration of O2 RNA and gate' RNA, 25 nmol/L of the O2 strand template and 18.75 nmol/L of the gate' strand template were used (Supplementary Figure 9).

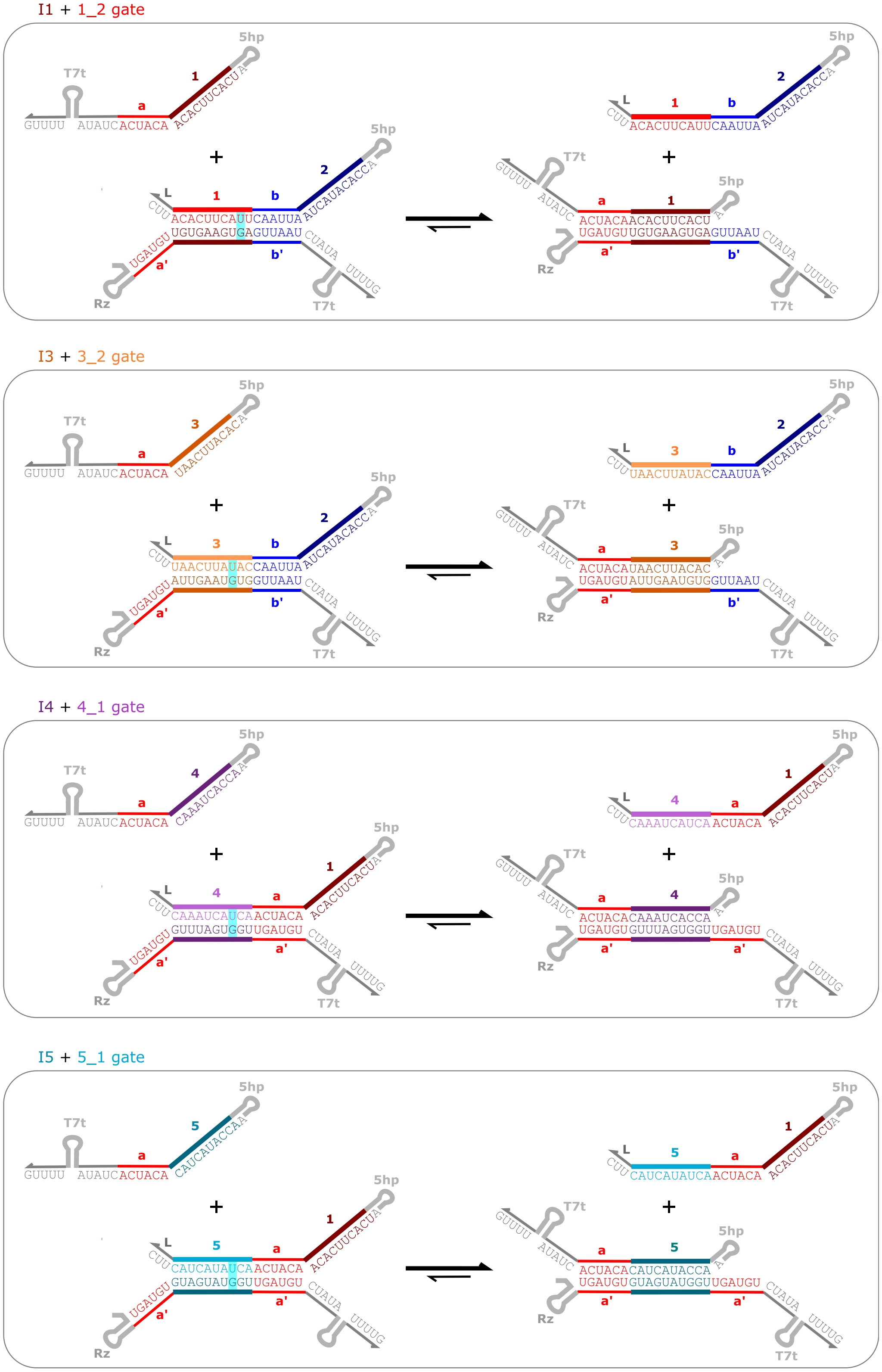


**Supplementary Figure 3:** Sequence schematics of ctRSD reactions between I1, I3, I4, or I5 and corresponding gates. G-U wobble base pairs are highlighted in cyan. An input reacting with its designed gate turns a G-U wobble pairing to a G-C pairing, providing a thermodynamic driving force for the forward reaction (*3*).

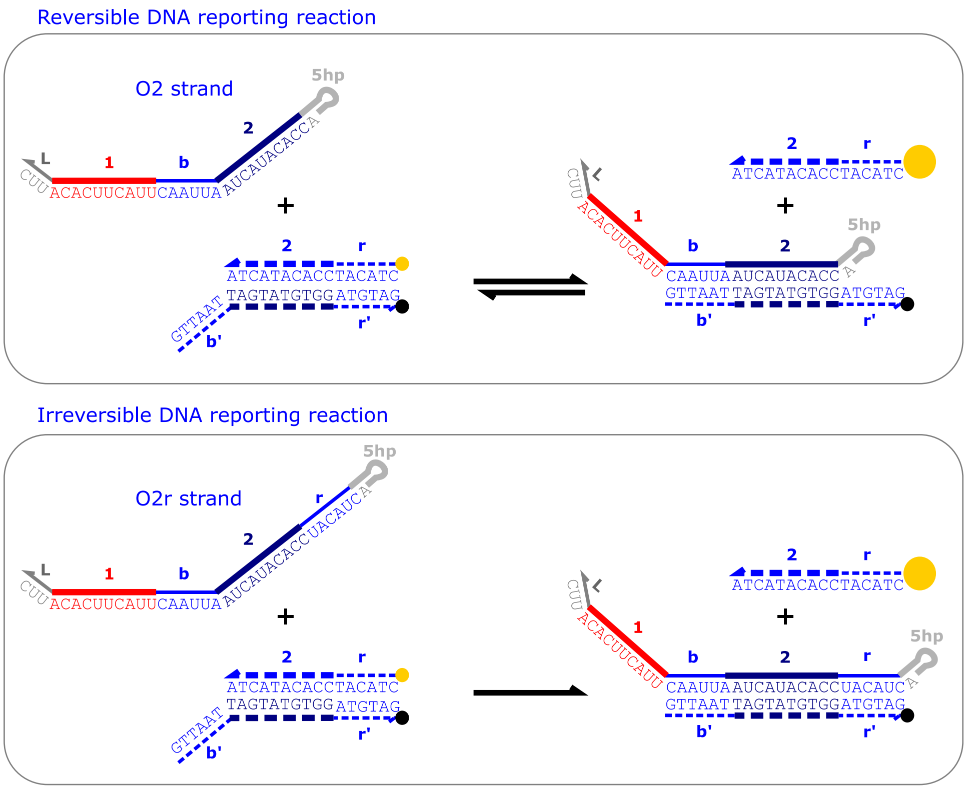


**Supplementary Figure 4:** Sequence schematics of the DNA reporter reacting with the O2 strand (top) or the O2r strand, which is extended to contain the reversible domain of the reporter output (bottom). To be stable at 37 °C, the DNA reporter needed a double stranded region of >15 bases, which prompted addition of the r-domain.

I3 + I1 + 3&1_2 gate

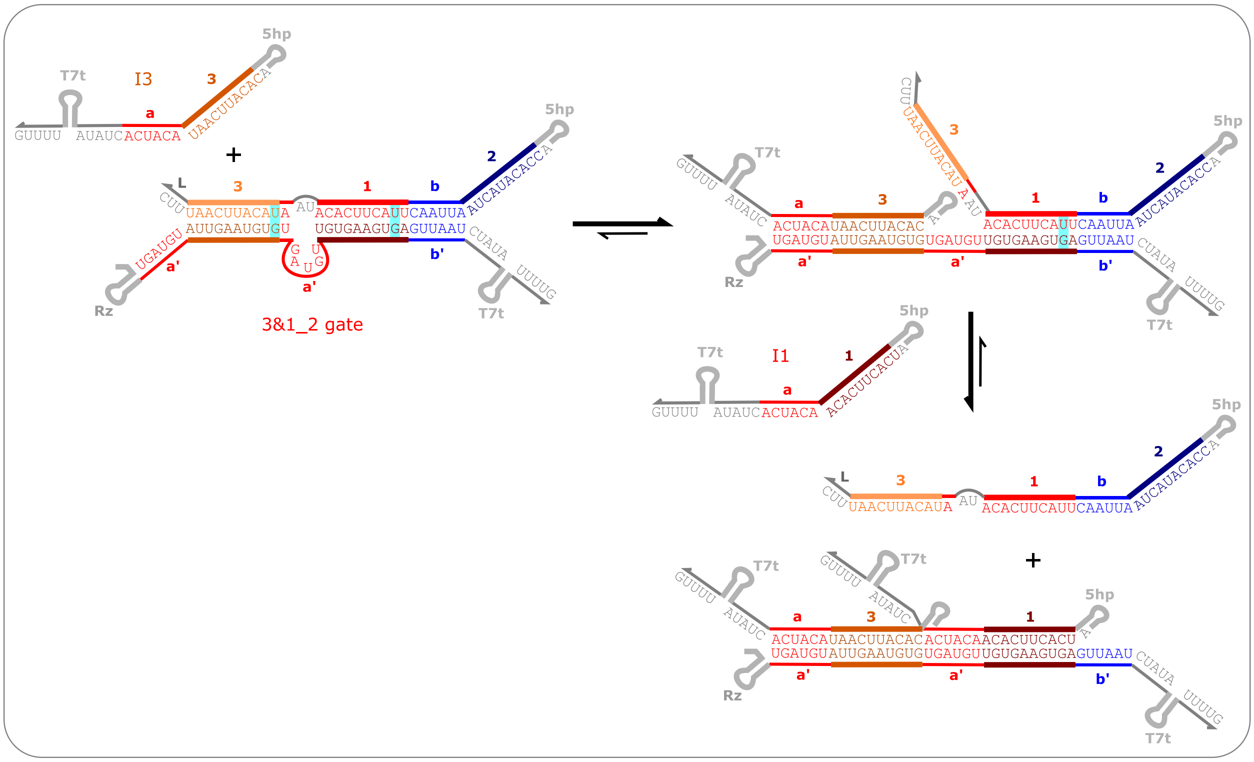


I5 + I4 + 5&4_2 gate

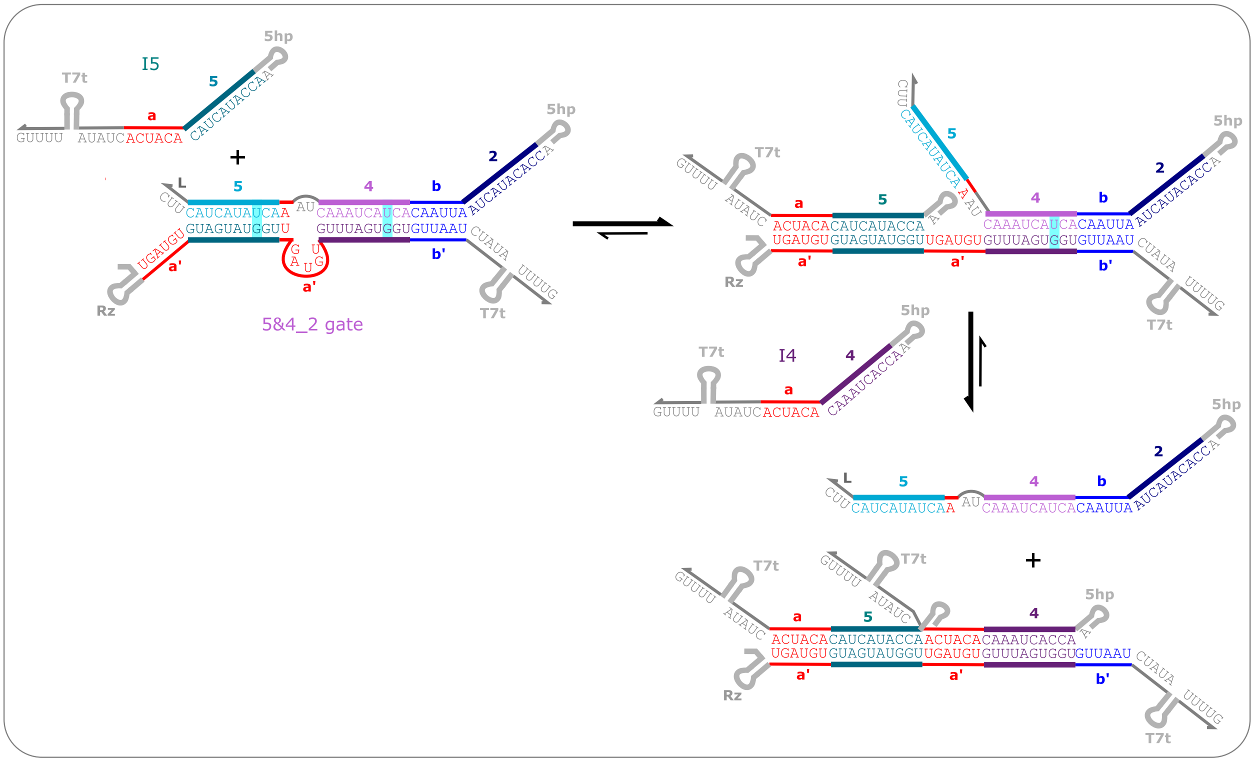


**Supplementary Figure 5:** Sequence schematics of ctRSD for 3&1_2 and 5&4_2 gates. G-U wobble base pairs are highlighted in cyan. An input reacting with its designed gate turns a G-U wobble pairing to a G-C pairing, providing a thermodynamic driving force for the forward reaction (*3*).

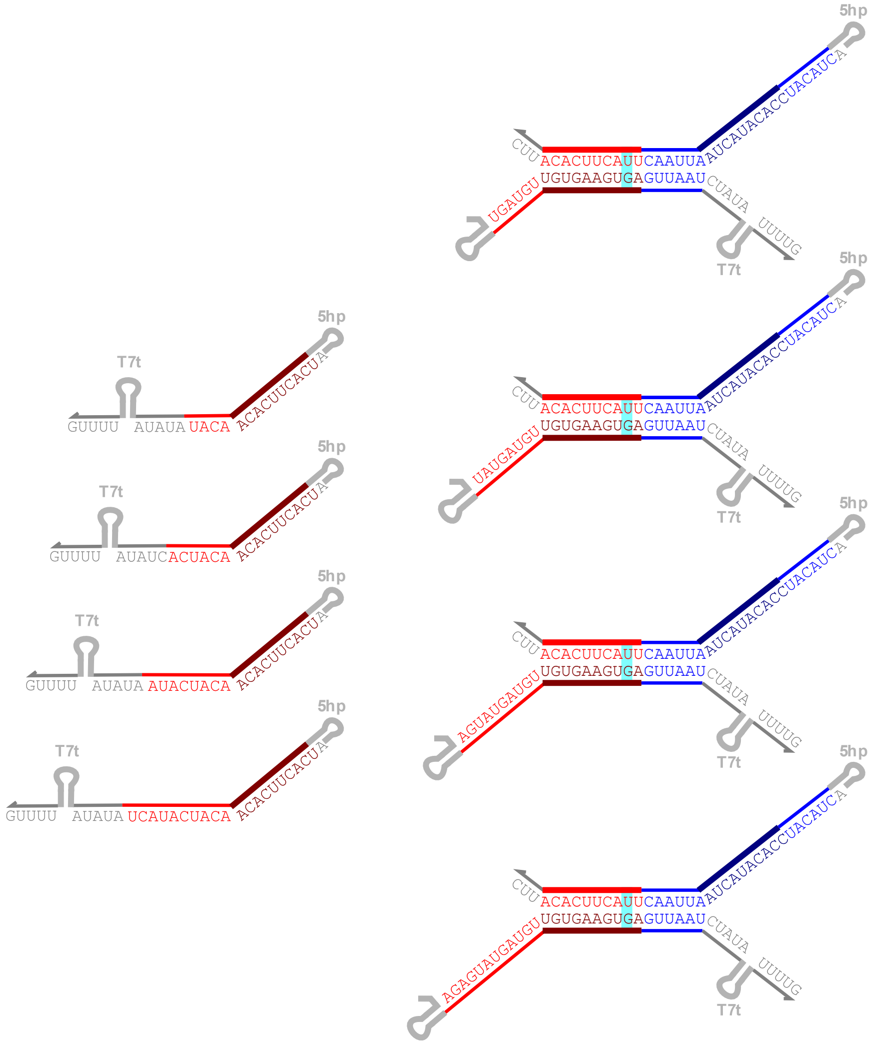


**Supplementary Figure 6:** Sequence schematics of ctRSD gates and inputs with variable toehold lengths. G-U wobble base pairs are highlighted in cyan. An input reacting with its designed gate turns a G-U wobble pairing to a G-C pairing, providing a thermodynamic driving force for the forward reaction (*3*).

### 2. ctRSD design considerations

This section details the different design considerations analyzed during development of the ctRSD gates and describes the motivation for specific design choices. Supplementary Section 2.1 compares two methods to transcriptionally encode RNA strand displacement gates: transcription of the two gate strands from separate transcription templates or transcription of an RNA hairpin with a ribozyme that cleaves the hairpin after folding to produce a dsRNA gate. The former method introduces a significant downstream leak reaction (Supplementary Section 2.1) and was not used. Supplementary Section 2.2 analyzes four different transcription paths for producing ctRSD gates. In principle, these different transcription paths are conceptually equivalent but depend on the selected toehold directionality (5′ vs 3′) and the position of the ribozyme within the transcript. Supplementary Section 2.3 analyzes three different self-cleaving ribozyme options for the ctRSD gates.

### 2.1 Separate transcription of gate strands results in downstream leak

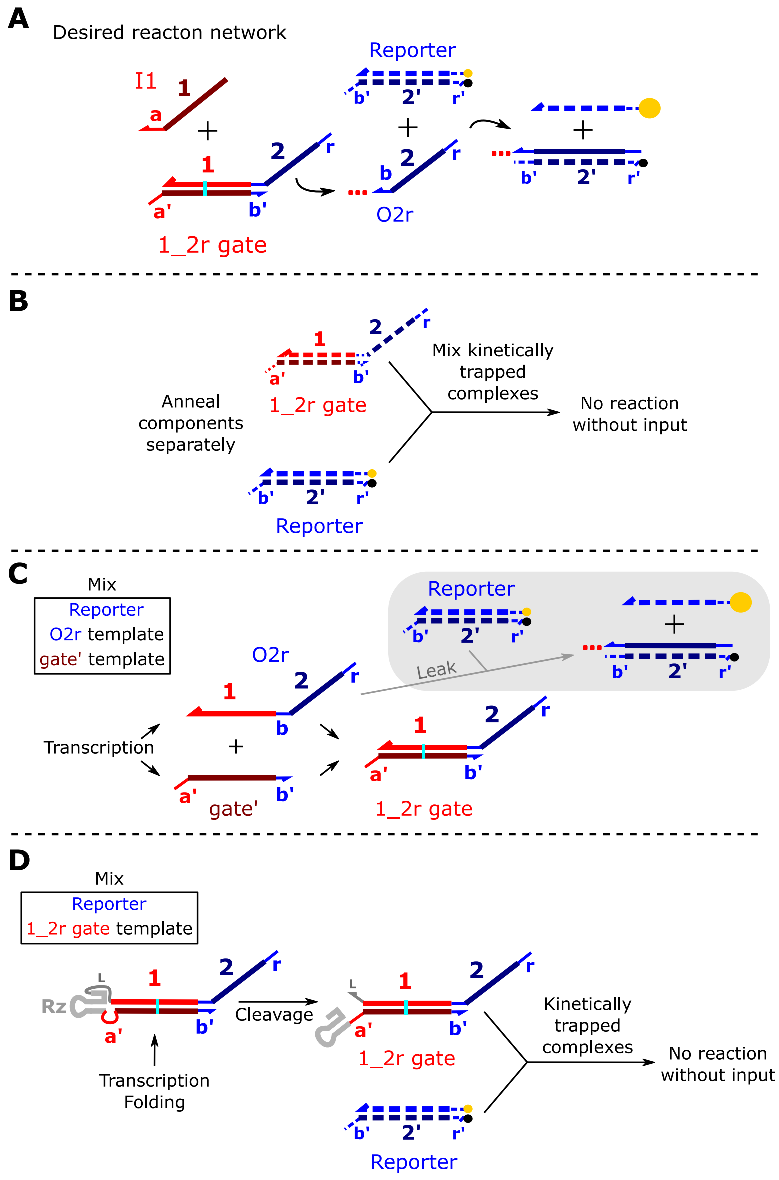


**Supplementary Figure 7:** Overview of options for preparing gates for TMSD circuits. (**A**) The desired reaction network. I1 releases an output from the 1_2r gate that in turn reacts with a DNA reporter complex to produce a fluorescent signal. The 1_2r gate should not react with the DNA reporter on its own. (**B**) The typical method for preparing DNA gates for TMSD circuits. The 1_2r gate and the DNA reporter are prepared in separate test tubes and then mixed to make the circuit. Separate preparation ensures the two complexes are kinetically precluded from reacting in the absence of input. (**C**) A method for preparing ctRSD gates in which separate transcription templates produce the output and gate' strands of the gate. Here, the two RNA strands should hybridize after transcription to form the gate. However, if this is done alongside the DNA reporter, or any other downstream gates that take O2r as their input, the output strand can also react downstream before it hybridizes to form the gate. This introduces significant leak in the circuit (Supplementary Figure 8). (**D**) A method for preparing ctRSD gates in which a gate is encoded as a hairpin that cleaves into a dsRNA product *via* an internal self-cleaving ribozyme. In this method, the hairpin rapidly folds during transcription to prevent any downstream reaction in the absence of input. This method for preparing ctRSD gates was used in this study.

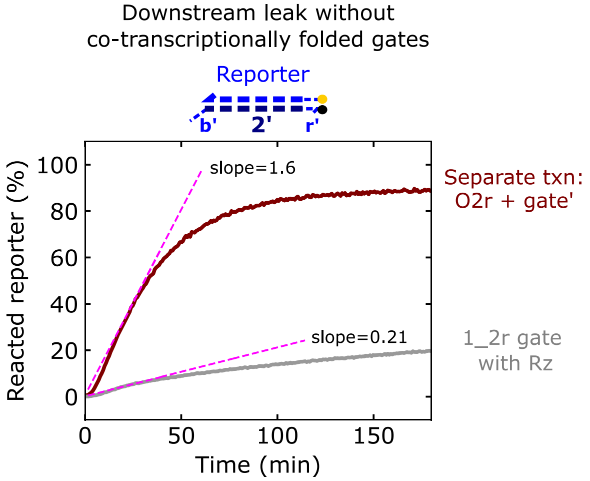


**Supplementary Figure 8:** Measuring downstream leak during transcription of the output (O2r) and gate′ strands of the 1_2r gate (Supplementary Figure 7C) or transcription of the self-cleaving 1_2r gate hairpin (Supplementary Figure 7D). The leak rate was determined by fitting a line (pink dotted lines) to the first 40 min of each reaction. Compared to separate transcription of the two gate strands, transcription of the 1_2r gate with the self-cleaving ribozyme reduces the leak 7.6-fold. To facilitate an irreversible reporting reaction, a 1_2 gate variant with a 6 base 5′ extension (domain r) was used (Supplementary Figure 4). Reactions were conducted with 500 nmol/L DNA reporter, 1 U/µL of T7 RNAP, and 25 nmol/L 1_2r gate and O2r templates. For a 1:1 concentration of O2r RNA and gate′ RNA, 18.75 nmol/L of the gate′ strand template was used (Supplementary Figure 9).

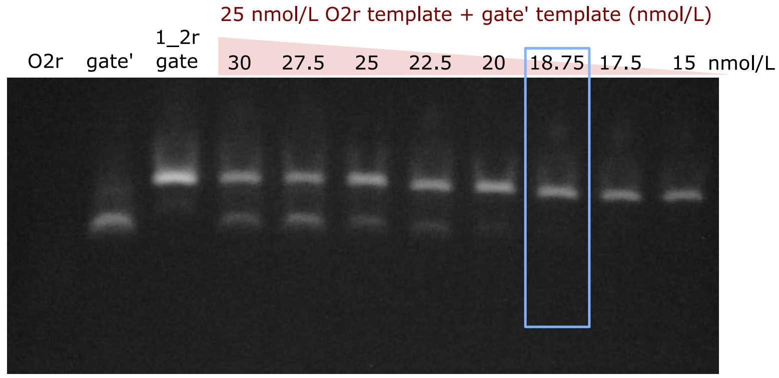


**Supplementary Figure 9:** Titrating the gate′ strand template concentration to achieve equal concentrations of O2r and gate′ transcripts. 25 nmol/L O2r template and 18.75 nmol/L gate′ template yield a 1:1 concentration (blue rectangle). Increasing the gate′ strand template concentration results in excess gate′ transcript. The O2r strand did not stain well. Transcription and native gel electrophoresis were conducted as described in the Methods of the main text. After transcription, samples were incubated with DNase I for 1 h before gel electrophoresis.

### 2.2 The co-transcriptional folding pathway

Considering the directionality of the single-stranded RNA (ssRNA) toehold that facilitates strand displacement and the placement of the self-cleaving ribozyme within the RNA transcript, there are four possible designs for ctRSD gates (Supplementary Figure 10). Previous work indicates that 5′ toeholds on RNA strand displacement gates perform better than 3′ toeholds (*5*, *6*), so we focused on designs with 5′ toeholds (Supplementary Figure 10A,B). The placement of the self-cleaving ribozyme influences which domains of the gate are transcribed first. For example, placing the ribozyme adjacent to the 5′ toehold results in transcription of the output region (2-, b-, and 1-domains) of the gate first, while placing the ribozyme on the opposite side of the transcript results in transcription of the gate′ strand first. In many DNA strand displacement circuits the output sequences of the DNA gates are constrained to a 3 letter code (C, A, or, T). This constraint reduces the possibility of unwanted secondary structure from forming and preventing output strands in larger circuits from hybridizing with each other (crosstalk) (*7*, *8*). We adopted the same sequence constraints in our RNA gate designs, limiting the gate output sequences (2-, b-, and 1-domains) to only C, A, or U bases. This constraint is particularly important for RNA circuits because G-U wobble base pairings are more energetically favored in RNA than G-T wobble pairings in DNA (*9*). Thus, even output sequences constrained to G, A, U bases could fold into undesired secondary structures. In the ctRSD gate design in which the self-cleaving ribozyme is placed opposite of the gate toehold (Supplementary Figure 10B), the gate′ strand (a′-, 1′-, and b′- domains), whose sequence would be composed of G, A, U bases, would be transcribed first. Given that co-transcriptional folding of RNA is much faster than transcription (*10*), transcription of the a′-, 1′-, and b′-domains first could result in undesired secondary structure in the transcript before the complementary 1- and b-domains are transcribed, hindering the correct gate formation. For these reasons, we chose the RNA gate design in which the 2-, b-, and 1-domains (composed of only C, A, U bases) are transcribed first (Supplementary Figure 10A).

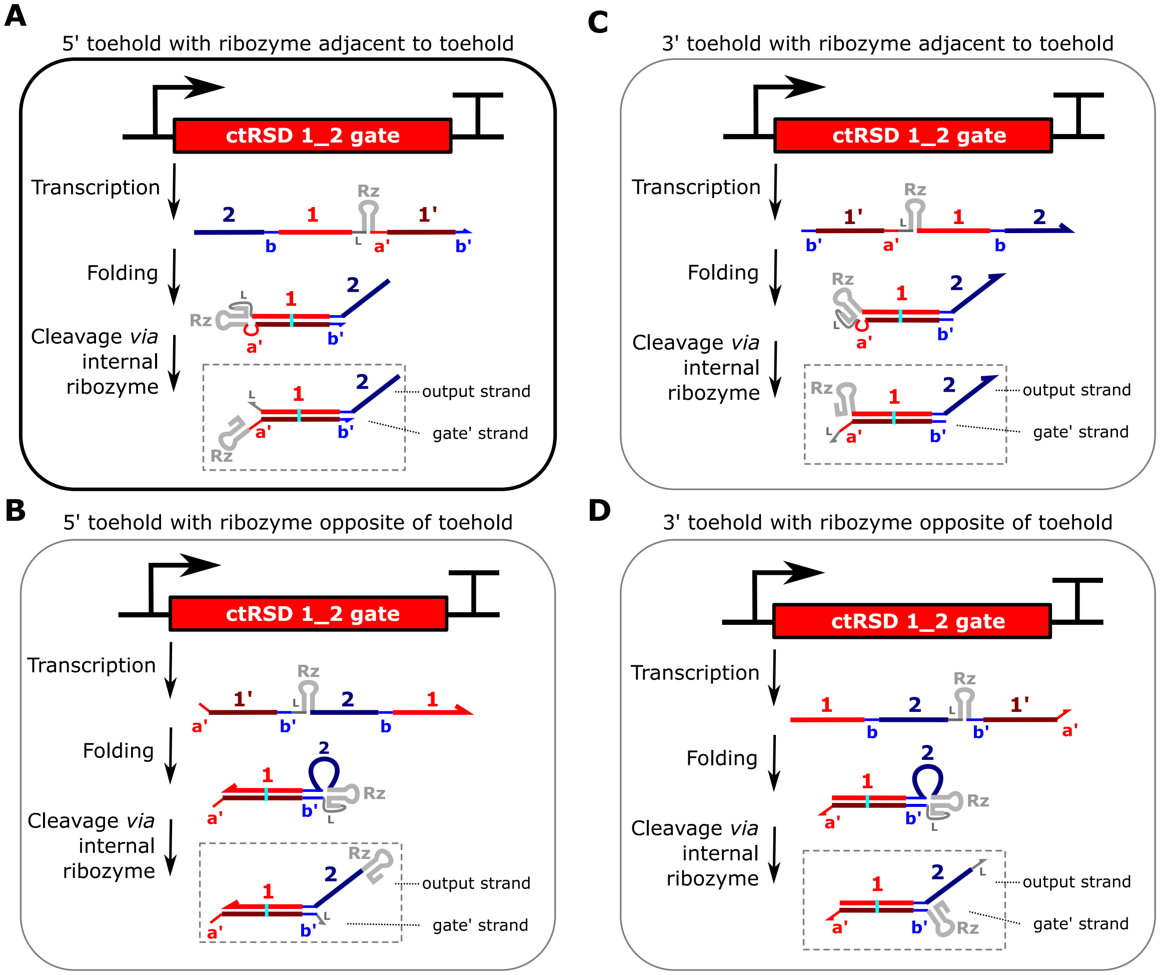


**Supplementary Figure 10:** The four possible folding pathways for designing ctRSD gates. The two major design choices are the placement of the ssRNA toehold—5′ (**A** and **B**) or 3′ (**C** and **D**)—and the placement of the self-cleaving ribozyme (Rz)—adjacent (**A** and **C**) or opposite (**B** and **D**) of the ssRNA toehold. The design in (A) was selected.

### 2.3 Self-cleaving ribozyme selection

Three well characterized ribozymes were considered: the hammerhead ribozyme, the hairpin ribozyme, and the hepatitis delta virus (HDV) ribozyme. The HDV ribozyme has several advantages over the hammerhead and hairpin ribozymes. First, the HDV ribozyme folds quickly into a stable structure (*11*, *12*), likely making it resistant to misfolding across different flanking sequences. Second, the rate constant for HDV ribozyme cleavage has been reported as 52 $min^{-1}$ in certain settings (*13*), compared to 1 $min^{-1}$ for the hammerhead (*13*, *14*) or 0.5 $min^{-1}$ to 0.05 $min^{-1}$ for the hairpin ribozymes (*14*, *15*). Lastly, the HDV ribozyme has little sequence preference upstream of the cleavage site (*16*). Both the hammerhead (*17*) and hairpin (*15*) ribozymes have cleavage site sequence constraints and their cleavage sites are flanked by RNA duplexes thus requiring a dissociation step following cleavage to separate the two strands. This dissociation step is particularly problematic in our ctRSD designs, in which the ssRNA toehold for strand displacement must be exposed after cleavage. In our designs, the hammerhead and hairpin ribozymes require 6 and 4 bases, respectively, to dissociate after cleavage to expose the toehold for strand displacement (Supplementary Figure 11). In the case of the hammerhead ribozyme, these 6 bases are likely to remain hybridized most of the time after cleavage, impeding RNA strand displacement. The HDV ribozyme does not suffer from these sequence limitations, driving this choice for our designs. We found the HDV ribozyme (sequence adopted from Ref (*16*)) resulted in the desired efficient and rapid cleavage in our RNA gates (Figure 1 and Supplementary Figure 12). We also tested a ctRSD gate with the hairpin ribozyme, but much less cleavage was observed than with the HDV ribozyme (Supplementary Figure 13).

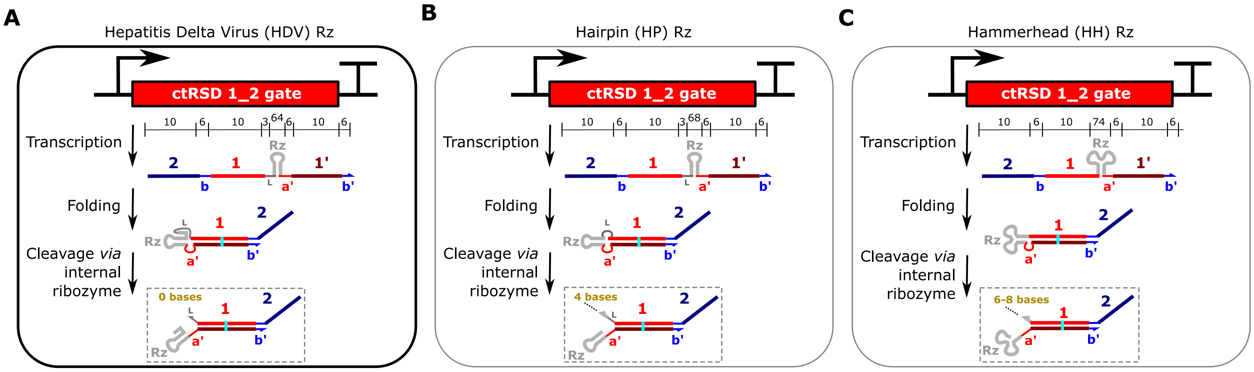


**Supplementary Figure 11:** Designs of ctRSD gates with different self-cleaving ribozymes. Numbers above transcripts indicate domain lengths in nucleotides. In the final cleavage products, the number of bases that must dissociate to expose the toehold are written in gold lettering. The HDV ribozyme does not require any bases to unhybridized after cleavage, making it the best candidate for our ctRSD gate design.

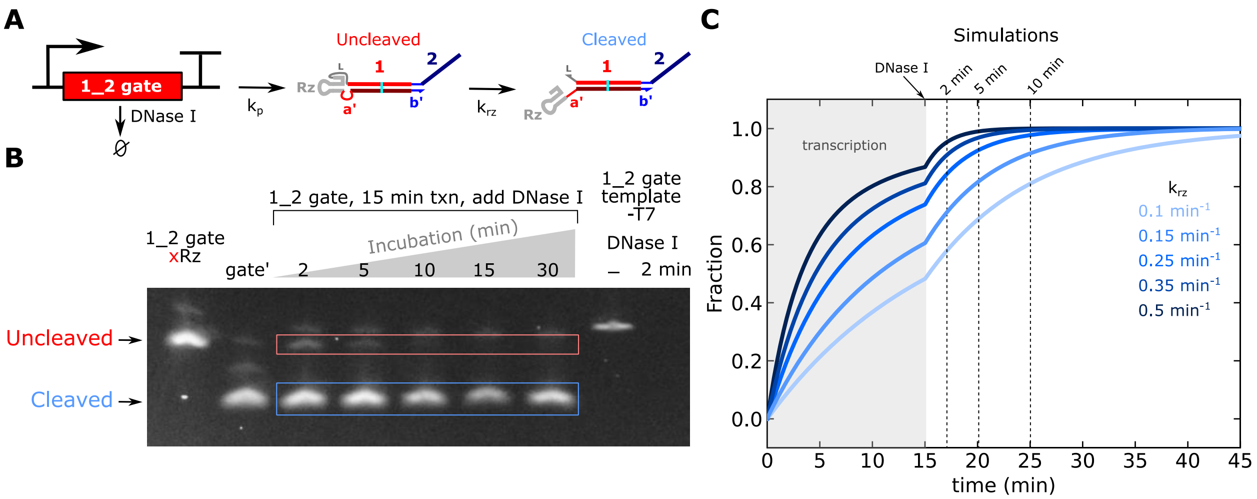


**Supplementary Figure 12:** Rate and efficiency of internal HDV ribozyme cleavage in ctRSD gates. (**A**) Schematic of experimental and simulation setup. A DNA template (25 nmol/L) is transcribed to produce the uncleaved ctRSD gate. $k_p$ is the first order transcription rate constant. The HDV ribozyme can then convert the ctRSD gate into the cleaved product with a first order rate constant $k_{rz}$. Transcription proceeded for 15 min. DNase I (final concentration 0.23 U/µL, 1.43 mmol/L $CaCl_2$) was then added to the samples, and the samples were incubated for varying times prior to the addition of denaturing solution and heating to 90 °C for 5 min. Following denaturing, the samples were immediately run on a 4 % EX agarose E-gel (Invitrogen) for 25 min and imaged. For denaturing, the denaturing solution (100 % formamide, 36 mmol/L EDTA) was mixed 1:1 by volume with the samples. (**B**) Denaturing gel results for the experiment described in (A). The first two lanes are controls showing the uncleaved and cleaved RNA products. These samples were incubated with DNase I for 30 min following transcription. The last gel lane shows that all the DNA template is degraded after 2 min with DNase I, suggesting degradation is rapid in our experimental conditions. Almost no detectable uncleaved transcript was left after a 10 min incubation with DNase I. (**C**) The fraction of cleaved ctRSD gate predicted from a simulation of the experiment depicted in (A). Mass action kinetic equations were used with a $k_p$ of 0.025 $s^{-1}$ and different values for $k_{rz}$. The value of $k_p$ does not influence the faction of cleaved RNA in simulation. The gray shaded region in the plot indicates the initial 15 min of transcription. At 15 min in the simulations, the DNA template concentration is changed to 0 nmol/L to simulate the addition of DNase I. Dashed lines indicate the 2 min, 5 min, and 10 min timepoints that were run on the gel in (B). Based on this analysis, we estimate $k_{rz}$ as approximately between (0.25 and 0.35) $min^{-1}$. This estimated rate is within a factor of 2 of previously published rates for HDV ribozyme self-cleavage *in vitro* (*14*).

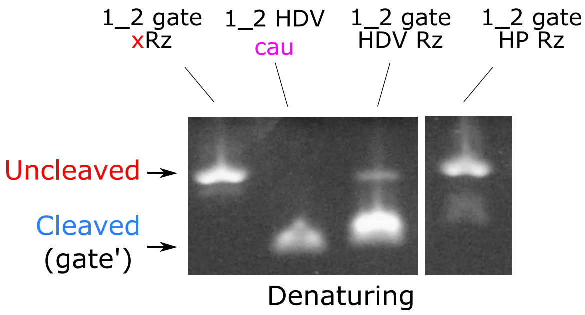


**Supplementary Figure 13:** Denaturing gel of the ctRSD 1_2 gate with the hepatitis delta virus (HDV) or the hairpin (HP) ribozyme. Most of the HP ribozyme transcript does not cleave. *In vitro* transcription reactions were conducted for 1 h, followed by DNase I digestion (Methods). Samples were mixed with an equal volume of Gel Loading Buffer II (Invitrogen), heated to 80 °C for 5 min, and subsequently run on 4 % EX agarose E-gels for 30 min. The 1_2 gate xRz and 1_2 HDV cau controls, which represent the full length uncleaved and cleaved products, respectively, are depicted in Supplementary Figure 2. All samples were analyzed on the same gel but lanes between the HDV Rz and HP Rz samples were removed because they were not related to this experiment.

### 3. Equilibrium analysis with NUPACK

NUPACK 3.2.2 (*18*) was used for equilibrium analysis of RNA complexes. We used the default NUPACK parameters for RNA (1.0 mol/L Na+ and 0 mol/L Mg++, dangles: some). Although there is 6 mmol/L $MgCl_2$ in our transcription buffer, there is a total of 8 mmol/L NTPs, which will sequester $MgCl_2$, so the concentration of free Mg++ is unknown. For RNA analysis, the default salt conditions are the only options. Unless otherwise state, analysis was conducted at 37 $^{o}$C with 1 µmol/L of each RNA species. Changing the equimolar concentration of the RNA species between 10 nmol/L and 100 µmol/L does not change the predicted equilibrium concentrations.

For analysis of the reaction I1 + 1_2 gate ↔ I1:gate′ + O2 the strands supplied to NUPACK are shown below:

```
I1: 5′GGGAGAUUCGUCUCCCAUCACUUCACAACAUCAAUAUAACCCCUUGGGGCCUCUAAACGGGUCUUGAGGGGUUUUUUG

1_2 strand(O2): 5′GGGAGAUUCGUCUCCCACCACAUACUAAUUAACUUACUUCACAUUC

1_2 gate′: 5′GGGUCGGCAUGGCAUCUCCACCUCCUCGCGGUCCGACCUGGGCUACUUCGGUAGGCUAAGGGAG
             UGAUGUUGUGAAGUGAGUUAAUCUAUAACCCCUUGGGGCCUCUAAACGGGUCUUGAGGGGUUUUUUG
```

The 1_2 gate′ sequence contains the HDV ribozyme sequence. However, the HDV ribozymes structure contains a pseudoknot, which NUPACK is incapable of predicting. Thus, the secondary structure of the HDV ribozyme in NUPACK does not represent its real structure. We found that the first two 5′ bases of the T7 RNAP terminator sequence n I1 (5′ CU) o were predicted to hybridize to part of the HDV ribozyme sequence on the 1_2 gate. However, this region of the ribozyme sequence is expected to be double stranded in the true ribozyme structure. To remove the influence of these spurious bases from the equilibrium analysis in NUPACK, the first C of the T7 RNAP terminator sequence was changed to an A (highlighted in yellow in the sequence above). This was done for all input sequences when analyzing these sequences in NUPACK.

### 4. Modeling ctRSD reactions

#### 4.1 Model assumptions and reactions

RNA strand displacement reactions were modeled using ordinary differential equations derived from mass action kinetics. All modeled reactions are shown in Supplementary Figure 14. In our model transcription was simplified to a first order process, whereby transcription rate is linearly proportional to the template concentration ($k_p$*[template]). This assumption ignores transcriptional loading effects (*19*) that arise when the concentration of polymerase is not in excess of the total concentration of total transcription templates. Thus, in our model, the first order transcription rate constant ($k_p$) depends both on the concentration of T7 RNAP and the total concentration of templates, *i.e.* transcriptional load (Supplementary Figure 28). To account for these dependencies, we used an experimental calibration sample to obtain $k_p$ values for a given T7 RNAP concentration and total template concentration. The $k_p$ value obtained from this calibration sample was then used to simulate experimental samples with the same conditions (Supplementary Figure 29). Because all of the transcripts in this study possess the same 5′ sequence, we assumed $k_p$ was the same for all transcripts (*20*). Because co-transcriptional folding is 10-fold faster than transcription (*10*), we assume that the gates fold instantaneously upon transcription, unless otherwise stated. Supplementary Section 4.2 discusses the rate constant values used in simulations.

Supplementary Section 4.3 describes the characterization of a leak reaction in the ctRSD system. This leak reaction was modeled by assuming that a small fraction of each ctRSD gate produced is as reactive as the designed output of the gate. Thus, a leak term was introduced in which an output is directly produced from its ctRSD gate template ($k_{pL}$*[ctRSD template]). $k_{pL}$ is the first order leak transcription rate constant. For single input ctRSD gates, we found a $k_{pL}$ that was 3 % of $k_p$ recapitulated our experimental observations. This 3 % leak transcription was used for all single input gates. We found that a 3 % transcriptional leak for AND gates resulted in less leak than we observed in experiments. We reasoned this might be because each AND gate possesses two dsRNA domains. If we assume that each dsRNA stem has a 97 % chance of being transcribed and folded correctly, we expect the chances an AND gate is correctly produced to be $(0.97)^2$ = 94.1 %. Based on this analysis, we assume a 6 % transcriptional leak ($k_{pLA}$) for all AND gates in the study. We also assumed the reactions between AND gates and their first inputs were irreversible because the reverse reaction is facilitated by a one base toehold. The reverse reaction between an AND gate and its final output was included in the model.

Beyond the leak reaction described above, our model ignores other potential side reactions that are not expected to significantly influence dynamics. First, any gate possessing an output complementary to another gate could react *via* a 0 base strand displacement mechanism. This reaction was not included in the model because it occurs two to three orders of magnitude slower (*21*) than the designed RNA strand displacement reactions. Second, an input can react with an RNA strand displacement gate prior to ribozyme cleavage. However, a mutant ctRSD gate that could not cleave reacted much slower with input than the self-cleaving ctRSD gate (Supplementary Figure 15). We assumed this side reaction would not greatly influence the observed kinetics at the low concentrations expected for the uncleaved ctRSD gate.

Our model implementation pools output strands from gates with different input domains. For example, if both a 4_1 and 5_1 gate are present in a simulation the model only tracks the total O1 produced and does not explicitly track O1 released from the 4_1 gate and the O1 released from the 5_1 gate

(Supplementary Figure 14). In most of our simulations, this issue does not arise as there is only a single gate releasing a given output. However, for circuits with OR elements, the reverse reaction for each gate will be overestimated. For example, for a 4_1 and 5_1 OR element the reverse rate for the 4_1 gate is $k_{rev}$*[I4:RSDg4]*[O1], but O1 can come from both the 4_1 and the 5_1 gate, and only O1 from the 4_1 gate can participate in this reverse reaction. In each OR gate experiment, the input template concentrations were equal and the gate template concentrations were equal. Thus, the overcounting of outputs would only change the reverse reaction rate 2-fold. For single-layer OR gates, the largest circuits simulated in which two gates produced the same output, a significant change in kinetics is not observed until the reverse rate constants are increased 25-fold (Supplementary Figure 16A,B). Based on these results, the overcounting in OR gate reverse reaction rates should not influence the results for the networks we simulated in this study. A more rigorous model that tracks which gates the outputs come from could become necessary as ctRSD circuits expand.

Finally, the model does not consider any loss of T7 RNAP activity or depletion of NTPs during transcription. Thus, the model is likely to become inaccurate when simulating experimental times > (4 to 5) h, as T7 RNAP activity will have decreased significantly (*22*). For slow reactions that are limited by transcription (*i.e.* transcription of leak products), the loss of T7 RNAP activity will eventually result in a plateau in output. The model will not capture this.

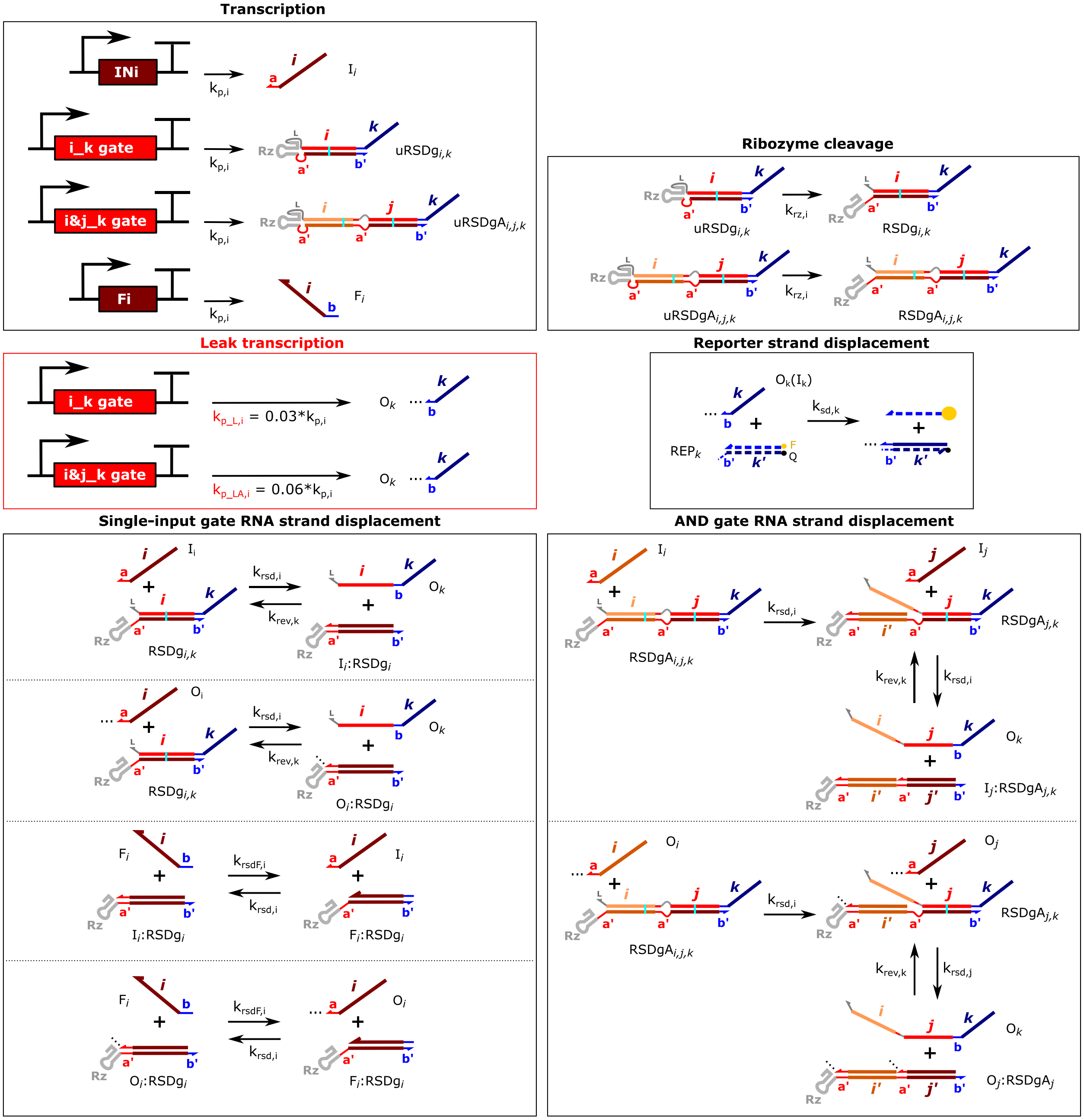


**Supplementary Figure 14:** Schematics of all reactions in the ctRSD kinetic model. Output templates can also be included in the model to simulate the transcription rate calibration samples (Supplementary Section 9).

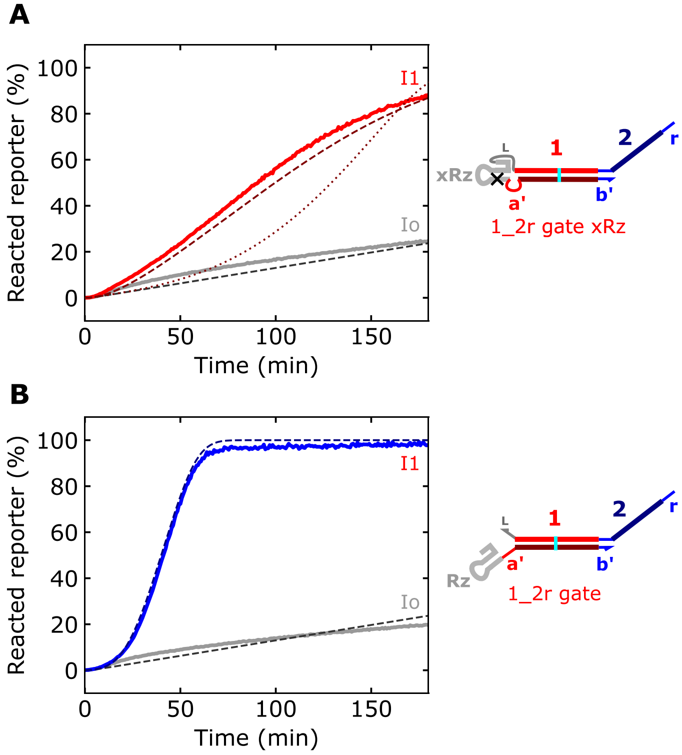


**Supplementary Figure 15:** An uncleaved ctRSD gate (**A**) produces output >3-fold slower than a cleaved ctRSD gate (**B**). 25 nmol/L of gate templates were used with 25 nmol/L of I1 or Io, 500 nmol/L of DNA reporter, and 1 U/µL of T7 RNAP. Experiments were otherwise conducted as described in the Methods of the main text. For simulation results (dashed lines), $k_p = 0.014\ s^{-1}$. In (A), two simulation results are shown. The dotted line is from a model in which $k_{rsd}$ was lowered to $1x10^1\ L\ mol^{-1}\ s^{-1}$ to simulate a slower forward strand displacement reaction. The dashed line is from a model in which $k_{rev}$ was increased to $1x10^6\ L\ mol^{-1}\ s^{-1}$ to simulate a fast reverse reaction due to a high effective concentration of the output strand. The Io simulation results are the same for both models. These experiments were conducted along side the experiments presented in Supplementary Figure 8 and the 1_2r gate + Io results in (B) are also shown there.

### 4.2 Kinetic parameters used to model co-transcriptional RNA strand displacement

In toehold mediated DNA strand displacement (DSD), the rate of the strand displacement reaction is correlated to the binding energy of the toehold. As binding energy increases with increasing toehold length, the same trend between toehold length and strand displacement rate enhancement is predicted for RSD as for DSD (*5*). Because rate enhancement is related to toehold binding energy, toehold sequence can also greatly influence the observed rate. For example, a strong 6 base toehold with high G-C content can result in rate constants near $10^6$ L mol$^{-1}$ s$^{-1}$, while weaker 6 base toehold sequences can result in rate constants closer to $10^4$ L mol$^{-1}$ s$^{-1}$ (*21*). The toehold on the DNA reporter contains five A or T bases and a single G, making it a weak toehold. Thus, a rate constant of $10^4$ L mol$^{-1}$ s$^{-1}$ was used to model the reaction between the reporter and the 1_2r strand ($k_{sd}$). All reporting reactions are considered to be irreversible.

For the rate constant of the reaction between an input strand and its corresponding ctRSD gate complex ($k_{rsd}$), we found a value of $10^3$ L mol$^{-1}$ s$^{-1}$ best recapitulated our experimental results. This value is at least two orders of magnitude lower than expected for a 6 base toehold with moderate GC content (*21*). There is some evidence that RSD reaction rate constants can be an order of magnitude lower than DSD reaction rates for short toeholds (*23*). Additionally, the presence of the bulky HDV ribozyme structure directly upstream of the toehold on the ctRSD gate could lower the observed reaction rate (Supplementary Figure 1). This bulky structure could sterically clash with the terminator hairpin at the 3′ end of the input strand and effectively decrease the strand displacement rate (*24*). In support of this hypothesis, we found the introduction of a 4 base single-stranded spacer between the HDV ribozyme motif and the toehold increased $k_{rsd}$ to $\geq 10^5$ L mol$^{-1}$ s$^{-1}$ (Supplementary Section 7). We assumed the $k_{rsd}$ value was the same for all ctRSD reactions, including for both toeholds of the AND gates. For the reaction between a fuel species and an input:gate′ complex, we assumed the same reaction rate constant as between the input and the ctRSD gate.

All the RNA strand displacement reactions in this study are reversible (Supplementary Figure 3). Estimation of the reverse reaction rate constants based on toehold length and sequence alone is confounded by reverse reactions replacing a G-C pair with a G-U wobble in the first two to three bases of branch migration (Supplementary Figure 3). This will reduce the rate of strand displacement, but the amount of this reduction is unknown. In DNA strand displacement, introduction of a mismatch at a similar position during branch migration can reduce the reaction rate by two to three orders of magnitude; presumably a G-U wobble would have a slightly less pronounced effect. Because we did not find an estimate for a comparable system in the literature, we estimated the reverse reaction rate constants from an equilibrium analysis of the strand displacement products. We used NUPACK 3.2.2 (*18*) to calculate the equilibrium constant (Keq) for each complementary gate and input sequence (Supplementary Section 3). The reverse reaction rate constant ($k_{rev}$) was determined from the equilibrium constant as $k_{rev} = k_{rsd}/Keq$. Based on this analysis, we found gates with outputs possessing the b-toehold have reverse rate constants nearly three orders of magnitude lower than the forward rate constants. Gates with outputs possessing the a-toehold have reverse reaction rate constants only one order of magnitude lower than the forward rates (Supplementary Table 2). To simplify the model, we assumed a single reverse rate constant for gates with b-toeholds (5 L mol$^{-1}$ s$^{-1}$) and for gates with a-toeholds (270 L mol$^{-1}$ s$^{-1}$). The reaction rate constants must be 10-fold larger than these values to begin to influence model predictions (Supplementary Figure 16).

The HDV ribozyme cleavage rate constant was estimated as 0.25 $min^{-1}$ (Supplementary Figure 12), consistent with previously reported *in vitro* values (*14*).

All the rate constants used in simulations are presented in Supplementary Table 3.

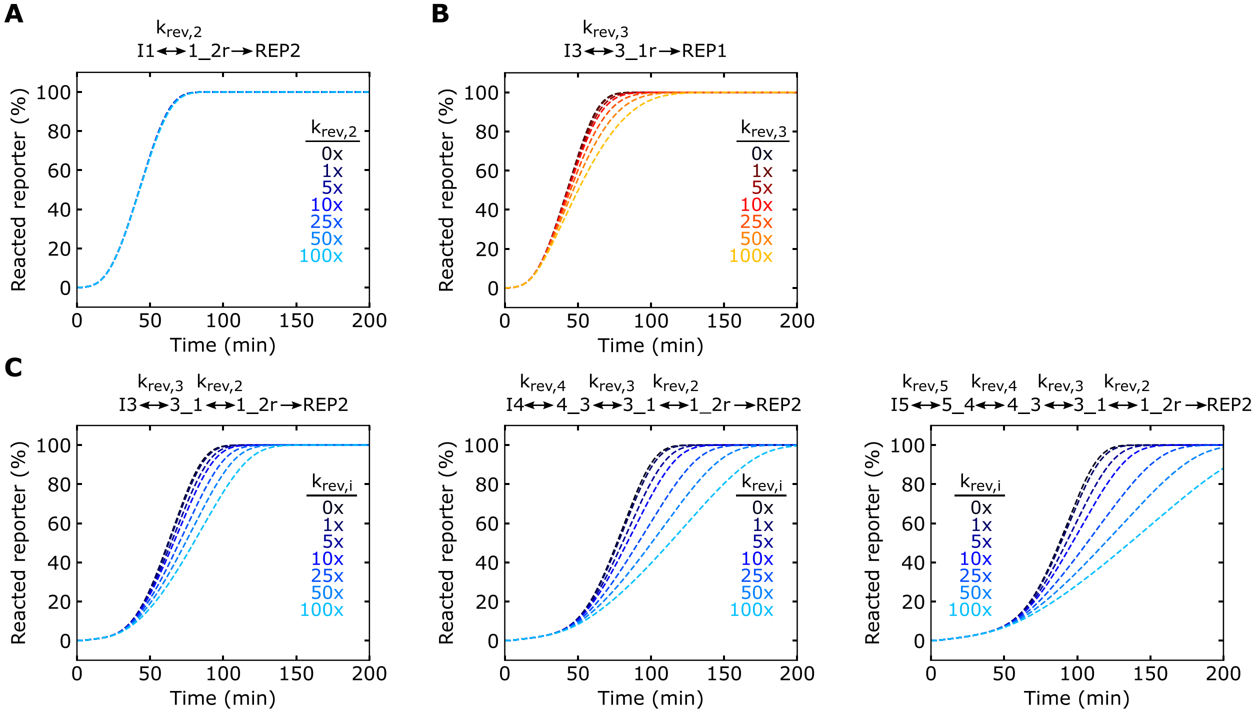


**Supplementary Figure 16:** The influence of reverse reaction rates on simulation results. The reverse rate must be 10-fold higher to significantly influence the results. (**A**) Simulations of a one-layer cascade producing O2r. Here, O2r initiates the reverse reaction with the weak b-toehold, which was modeled at 1x as 5 L $mol^{-1}$ $s^{-1}$. (**B**) Simulations of a one-layer cascade producing O1r. Here, O1r initiates the reverse with the a-toehold, which was modeled at 1x as 270 L $mol^{-1}$ $s^{-1}$. (**C**) Simulations of two-, three-, and four-layer cascades producing O2r as the final output.

**Supplementary Table 2:** Equilibrium analysis of ctRSD reactions across different gates and inputs. See Supplementary Section 3 for NUPACK analysis details.

| Gate | I + gate ↔ I:gate + Out (Keq) | $k_{rev}$ (L mol$^{-1}$ s$^{-1}$) |
|---|---|---|
| 1_2 gate | $(0.93)^2/(0.07)^2$ = 176.51 | 5.67 |
| 3_2 gate | $(0.905)^2/(0.095)^2$ = 90.75 | 11.01 |
| 4_2 gate | $(0.942)^2/(0.058)^2$ = 263.78 | 3.79 |
| 5_2 gate | $(0.939)^2/(0.061)^2$ = 236.36 | 4.22 |
| 1_4 gate | $(0.65)^2/(0.35)^2$ = 3.45 | 289.86 |
| 3_4 gate | $(0.62)^2/(0.38)^2$ = 2.66 | 375.94 |
| 4_1 gate | $(0.72)^2/(0.28)^2$ = 6.61 | 151.29 |
| 5_1 gate | $(0.66)^2/(0.34)^2$ = 3.77 | 265.25 |

**Supplementary Table 3:** Kinetic rate constants used in simulations. The last two rate constants in red text are rate constants used to model the leak transcription reaction for single input gates (kpL) and AND gates (kpLA).

| Parameter | Value | Source |
|---|---|---|
| $k_p$ (s$^{-1}$) | 0.0075 to 0.019 | Calibrated for transcriptional load |
| $k_{rz}$ (s$^{-1}$) | 4.167 x $10^{-3}$ | Supplementary Figure 12 |
| $k_{sd}$ (L mol$^{-1}$ s$^{-1}$) | 1 x $10^4$ | Ref (*21*) |
| $k_{rsd}$ (L mol$^{-1}$ s$^{-1}$) | 1 x $10^3$ | Supplementary Section 7 |
| $k_{rsdF}$ (L mol$^{-1}$ s$^{-1}$) | 1 x $10^3$ | Supplementary Section 7 |
| $k_{rev,a}$ (L mol$^{-1}$ s$^{-1}$) | 2.7 x $10^2$ | Supplementary Table 2 |
| $k_{rev,b}$ (L mol$^{-1}$ s$^{-1}$) | 5 | Supplementary Table 2 |
| $k_{pL}$ (s$^{-1}$) | 3 % of $k_p$ | Calibrated to data |
| $k_{pLA}$ (s$^{-1}$) | 6 % of $k_p$ | Calibrated to data |

### 4.3 Modeling and characterizing leak

In our experiments, we observed a leak in which transcription of the 1_2r gate template in the absence of the I1 template resulted in a slow increase in DNA reporter signal. This leak reaction increased with increasing concentrations of T7 RNAP, *i.e.* the leak increased with increasing transcription rate (Supplementary Figure 17). The initial ctRSD model did not include terms capable of producing this leak (Supplementary Figure 18A,B). To include this observed leak in the model, we investigated three potential models of leak the pathway: Models 1, 2, and 3 in Supplementary Figure 18C. Both Model 2 and Model 3 can recapitulate the experimental data, but only Model 3 is consistent with the experimental results presented in Supplementary Figure 18. Unless otherwise stated, all other simulations included the leak reactions depicted in Model 3.

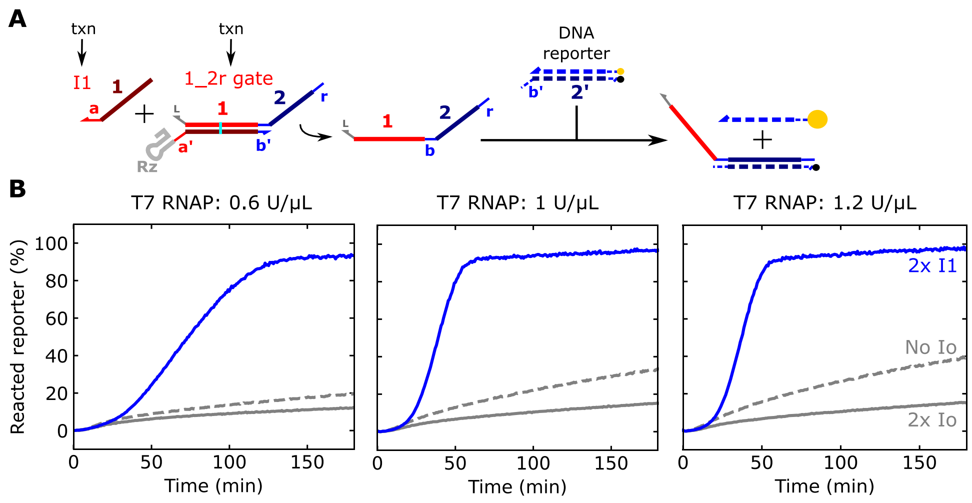


**Supplementary Figure 17:** ctRSD leak depends on polymerase concentration and transcriptional load. (**A**) RNA strand displacement reactions coupled with fluorescence reporting. (**B**) Normalized reporter signal during transcription of 25 nmol/L of the 1_2r gate template with either 50 nmol/L I1 template (blue line), 50 nmol/L Io template (gray line), or no input template (gray dashed line). From left to right, the concentration of T7 RNAP is increased. Increasing the polymerase concentration increased the rate of output production both with and without input. The transcriptional load in the sample with 50 nmol/L I1 and 25 nmol/L 1_2r gate is higher than the sample with 25 nmol/L 1_2r gate alone. To ensure the transcriptional load between the samples with and without I1 is the same, we tested another sample with the Io template (which produces an input RNA that does not react with the 1_2r gate) instead of the I1 template. Inclusion of the Io template reduces the leak from the 1_2r gate and ensures the same transcriptional load across samples for comparison. 500 nmol/L of DNA reporter was used in all samples.

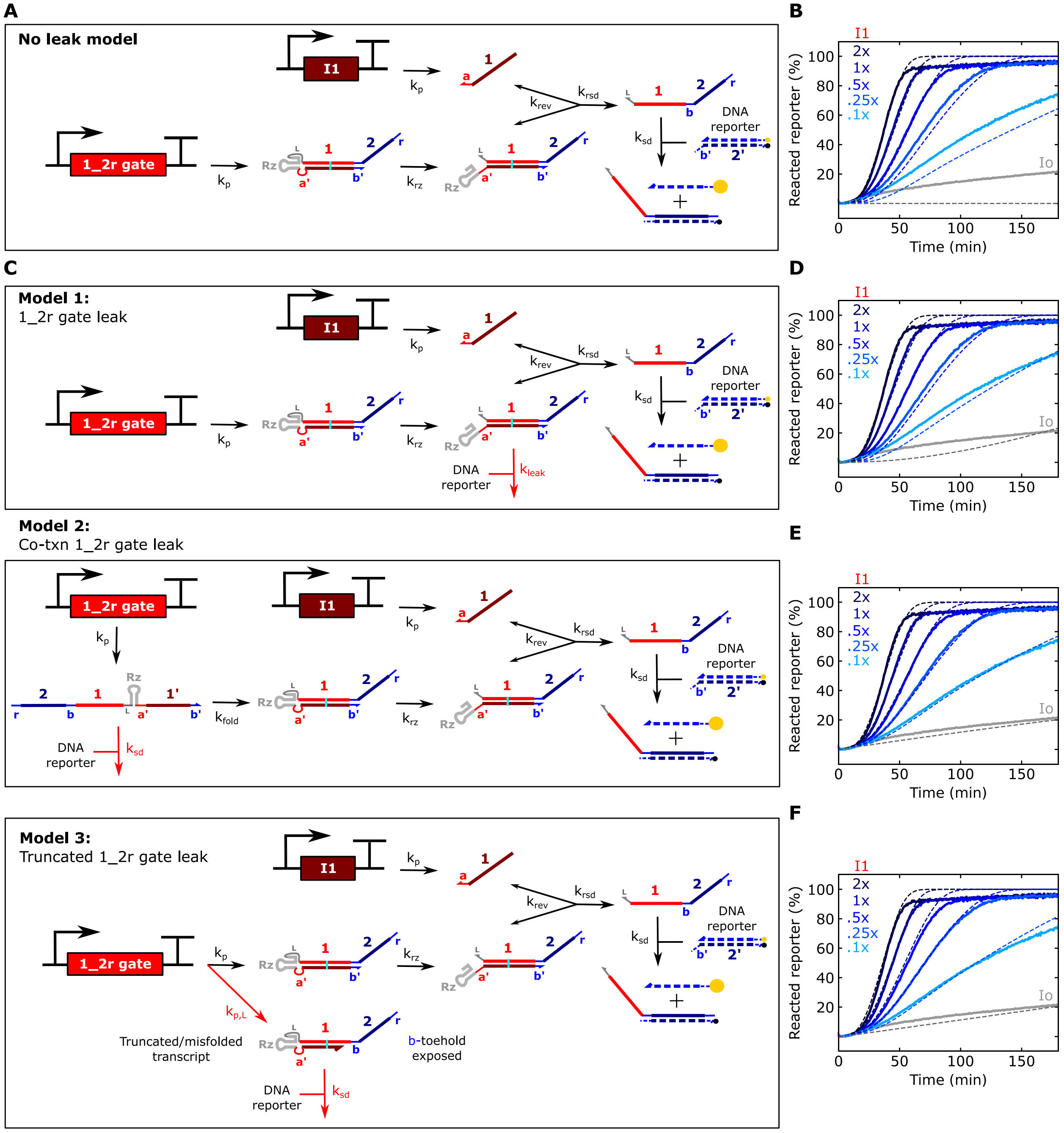


**Supplementary Figure 18:** Four different ctRSD models considered in this study. (**A**) Schematic of the model without any leak reactions. (**B**) Simulation results (dashed lines) for the model in (A) compared to experimental results (solid lines). (**C**) Schematics of the different leak models investigated. Each model includes a different pathway for a leak reaction between the 1_2r gate and the DNA reporter (red reaction lines). In Model 1, the leak is modeled as a 0 base toehold reaction (*25*) between the cleaved 1_2r gate and the DNA reporter. In Model 2, an additional folding step for the 1_2r gate transcript was introduced into the model. This model assumes the 1_2r gate RNA can react with the DNA reporter complex before the gate has folded. Because such a reaction would use the 6 base b-toehold—as does the designed reaction between the 1_2r strand—the leak was assumed to have the same rate constant as the designed reporting reaction ($k_{sd}$). In Model 3, it is assumed that a certain percentage of the 1_2r gate transcripts produced are truncated or misfolded. In the case of a truncated product, the b-toehold would be exposed, and thus the truncated product could react with the DNA reporter complex with the same rate constant as the designed reporting reaction.

Caption continued below…

(**D**) Simulation results (dashed lines) for Model 1 compared to experimental results (solid lines). In the simulations, a $k_{leak}$ of 15 L $mol^{-1}$ $s^{-1}$ was used for the 0 base toehold reaction between the 1_2r gate complex and the DNA reporter. This is an order of magnitude higher than reported previously (*21*, *25*). (**E**) Simulation results (dashed lines) for Model 2 compared to experimental results (solid lines). In the simulations, a $k_{fold}$ of 0.15 $s^{-1}$ was used. Considering that co-transcriptional folding occurs much faster than transcription (*10*), the $k_{fold}$ parameter may be taken as the time required to produce the transcript, during which the nascent transcript could react with the DNA reporter. A $k_{fold}$ of 0.15 $s^{-1}$ corresponds to a transcript produced every 6.67 s, and this corresponds to the transcription rate of ≈27 nt/s for the 183 nt 1_2r gate transcript. This transcription rate is within a factor of 1.5 to 3.5 of previously reported transcription rates for T7 RNAP (*26*, *27*), supporting the feasibility of the $k_{fold}$ parameter that recapitulates the experimental data. (**F**) Simulation results (dashed lines) for Model 3 compared to experimental results (solid lines). In the simulations, a production rate of truncated 1_2r gate products ($k_{p,L}$) that was 3 % of the production rate of correct products ($k_p$) was used. The reaction between the DNA reporter and the truncated 1_2r gate product was assumed to have the same rate constant ($k_{sd}$) as the reaction between the DNA reporter and the 1_2r strand. All other rate constants are in Supplementary Table 3. The experimental results are also presented in Figure 2 of the main text.

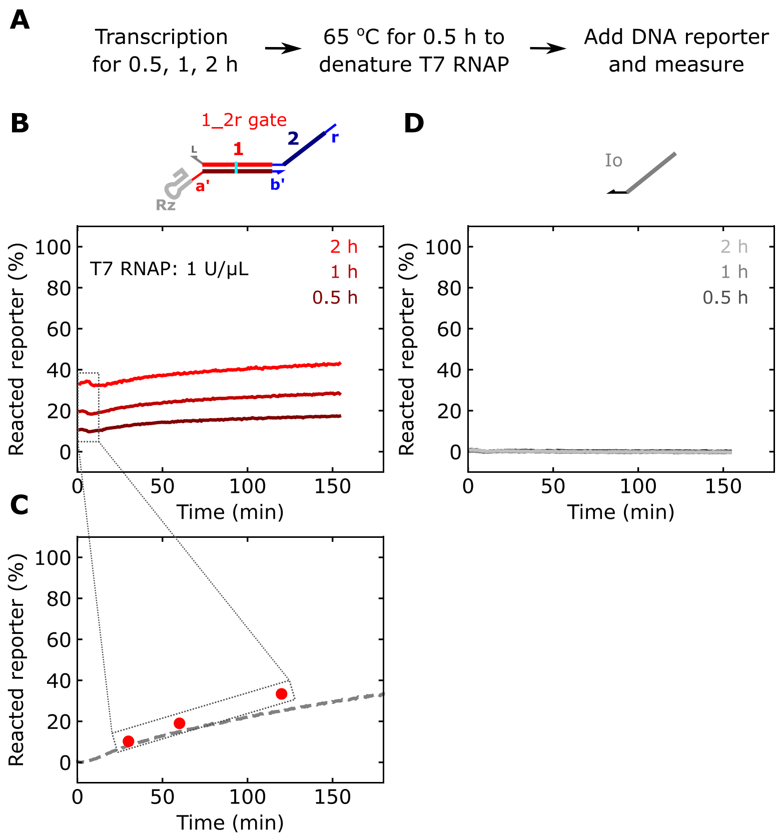


**Supplementary Figure 19:** Experimentally testing the leak pathways in Models 1, 2, and 3. (**A**) Experimental set up. The 1_2r gate template or the Io template (negative control) were transcribed with 1 U/µL T7 RNAP for (0.5, 1, or 2) h. After each transcription timepoint, the samples were heated to 65 °C for 30 min to denature T7 RNAP. After the heat denaturing step, the samples were returned to 37 °C until the end of the experiment. Once all timepoints had been denatured, the DNA reporter was added to each sample, and fluorescence measurements were begun immediately in the plate reader. If Model 1 described the leak, the reporter signal should start at 0 and slowly increase. If Model 2 described the leak, the reporter signal should start at 0 and remain there indefinitely because the 1_2r gate is folded and no transcription is occurring. If Model 3 described the leak, the reporter signal should immediately increase above 0. The longer the transcription time before heat denaturing, the higher this initial increase. (**B**) Normalized reporter signal starting right after 500 nmol/L of the DNA reporter was added to the samples containing the 1_2r gate transcripts. (**C**) The average normalized reporter signal values from the first 10 min of the data in (B) superimposed over leak of the 1_2r gate transcribed alongside the DNA reporter. (**D**) Normalized reporter signal from a negative control, in which the Io template was used instead of the 1_2r gate template in the experiment described in (A). This control demonstrates that the assay itself does not influence DNA reporter signal. The results in (B) and (C) support the leak pathway of Model 3 in Supplementary Figure 18 and are inconsistent with the pathways of Model 1 and Model 2.

### 5. Design of additional ctRSD sequence domains and circuit elements

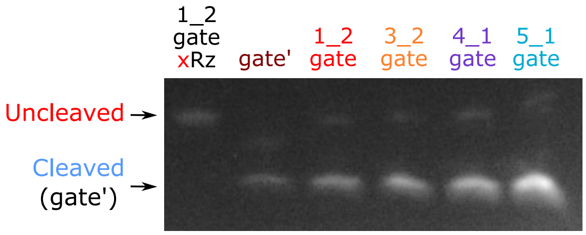


**Supplementary Figure 20:** Denaturing gel of ctRSD gates with 1, 3, 4, and 5 input domains. All gates self-cleave as designed. Transcription and gel electrophoresis were conducted as described in the Methods of the main text.

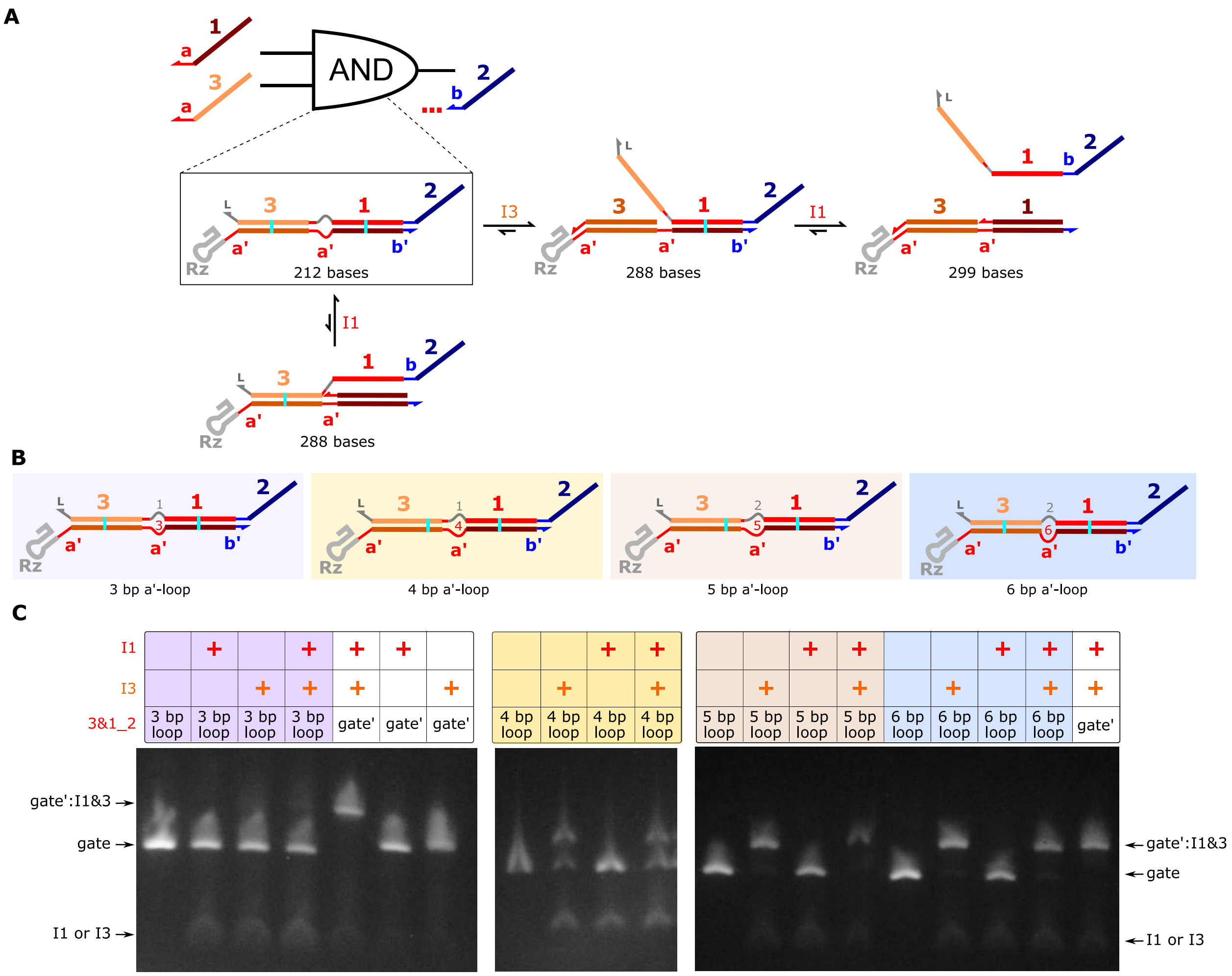


**Supplementary Figure 21:** Native gels of 3&1_2 AND gates with different sized a' internal loops between input domains 3 and 1. (**A**) Schematics of the desired AND gate design. The 3&1_2 gate is designed so that I3 must first react with the gate to expose the toehold for I1 to react and release the output. To accomplish this, we introduced an internal loop in the a:a'-domain between the 3 and 1 duplexes on the AND gate. The gray domains in the internal loops are short linker domains added to reduce strain between the 3 and 1 duplexes. A tradeoff to consider when introducing these internal loops pertains to the following: the a'-loop is the toehold for I1. The longer the a'-loop, the higher the rate at which I1 can react with the AND gate in the absence of I3. We designed AND gates with (3, 4, 5, and 6) base a'-loops to find a design that favored the reaction with I3, while disfavoring the reaction with I1 alone. (**B**) Schematics of the different AND gate designs tested. (**C**) Native gel shift assay results for the AND gate variants in (B). In these experiments, a higher molecular weight product should appear both when I3 is present and when I1 and I3 are present. Additionally, I1 by itself should not produce a higher molecular weight product. The AND gate with a 3 base a'-loop does not react with I3 or I1+I3. The AND gate with a 4 base a'-loop reacts ≈50 % with I3 and I1+I3. The AND gates with 5 base or 6 base a'-loops fully react with I3 and I1+I3. The larger the a'-loop, the more likely I1 will react with the AND gate in the absence of input. Thus, we selected the 5 base a'-loop design. The samples were run on the gels 2 h after the DNA templates were degraded with DNase I. Experiments were otherwise conducted as described in the Methods of the main text. The three images were taken from three different gels.

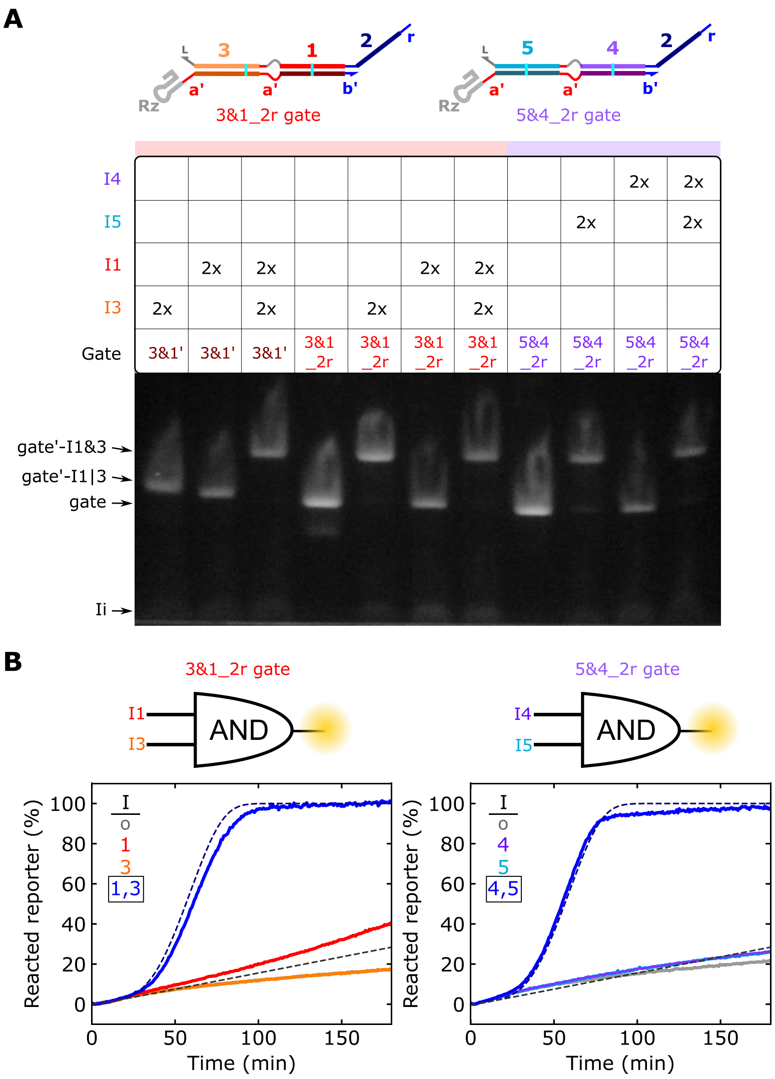


**Supplementary Figure 22:** AND gate characterization with the native RNA gel shift assay (**A**) and the DNA reporter assay (**B**). The 5&4_2r experiment was conducted using the same gate, input(s), and T7 RNAP concentrations as the 3&1_2r circuit element (Supplementary Table 4). For the gel electrophoresis results, gate and input templates were at 25 nmol/L and 50 nmol/L (2x), respectively. Electrophoresis was conducted 1 h after DNase I addition. The gate' strand is from the 3&1_2r gate. The first seven lanes of the gel are also presented in Figure 4E of the main text. The DNA reporter 3&1_2r results are also presented in Figure 4F of the main text.

### 6. Analysis of deviations between experiments and simulations

Across our experiments there were two minor deviations from simulation predictions. Deviation 1: There was lower leak than predicted between ctRSD gates, which could be the result of steric hinderance between leak products and gates (Supplementary Figure 23). Deviation 2: The gates that take I4 as an input reacted slower than the gates that take other inputs. The 4_2r gate was noticably slower than the three other single input gates tested (Figure 3a of the main text), and the 4_1 gate was slower than the 5_1 gate in a two-layer cascade (Supplementary Figure 24B). The 4_1 gate was also slower in a logic cascade than the 5_1 gate (Supplementary 24D). We hypothesized that the strand displacement rate constant for gates that take I4 as an input could be lower than the other domains due to the high UA content at the start of branch migration (Supplementary Figure 3). Similar sequences have been shown to significantly decrease overall strand displacement kinetics (*28*). Decreasing $k_{rsd}$ 4-fold in our simulations for just the gates that take I4 as an input resulted in better model agreement across experiments (Supplementary Figure 24).

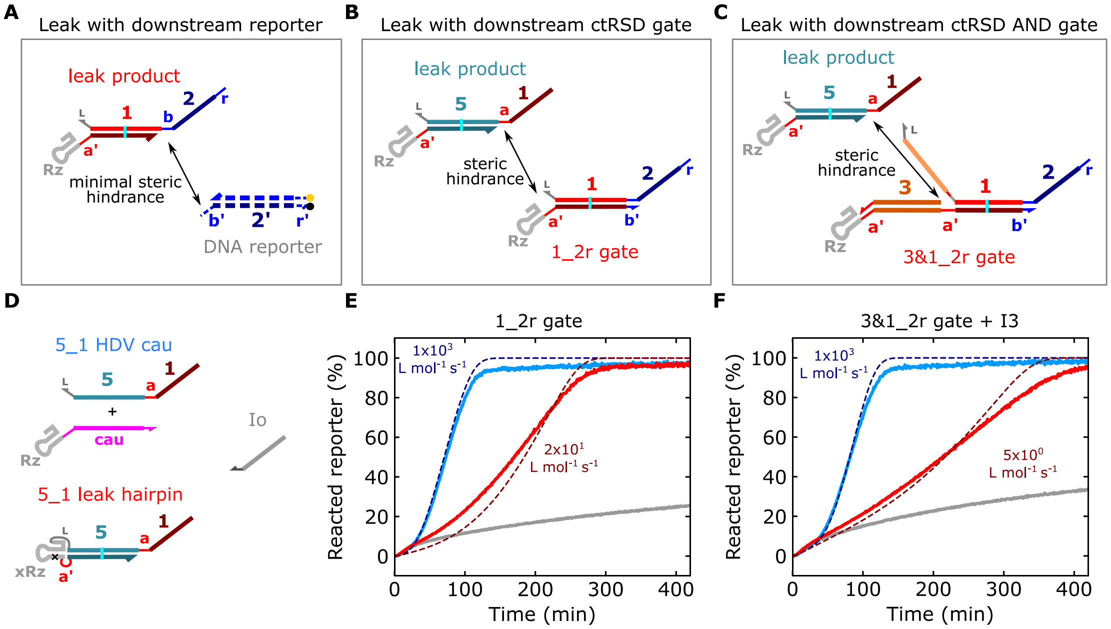


**Supplementary Figure 23:** Steric hinderance between the leak products and ctRSD gates reduces the leak observed in experiments. (**A**) Nothing upstream of the 5′ toehold of the DNA reporter sterically hinders a bulky leak product. Thus, the leak product likely reacts with the DNA reporter at a similar rate as a single-stranded output. (**B** and **C**) Single-input ctRSD gates (B) and ctRSD AND gates (C) have duplexes upstream of their 5′ toehold, which could sterically hinder a reaction with a bulky leak product. Note the leak products shown here are hypothetical but are representative of the true leak products, which are likely bulkier than their ssRNA output strands. (**D**) Schematics of the single-stranded 5_1 output strand and the 5_1 leak hairpin product used in the experiments in (E) and (F). The 5_1 HDV cau transcript contained the output strand of the 5_1 gate, followed by the HDV ribozyme and a 22 base sequence composed of only C, A, U bases. Upon transcription, this transcript cleaves to produce the output from the 5_1 gate. The 5_1 leak hairpin transcript was truncated such that the a-toehold of the 5_1 gate output was exposed. Further, an inactive ribozyme variant (xRz) was used to ensure the transcript remained in a hairpin structure. (**E**) Experimental (solid) and simulated (dashed) reporter signal during co-transcription of the 1_2r gate and the RNA products in (D). (**F**) Experimental (solid) and simulated (dashed) reporter signal during co-transcription of the 3&1_2r gate and I3 and the RNA products in (D). The color of the experimental data corresponds to the color of the transcript names in (D). [1_2r gate template] = [3&1_2r gate template] = [I3 template] = 25 nmol/L. [5_1 HDV cau template] = [5_1 leak hairpin template] = 10 nmol/L. Io was added to bring the total template concentration up to 75 nmol/L in each sample. [T7 RNAP] = 1 U/µL. For the simulations, a $k_p$ of 0.01 $s^{-1}$ was used, and the RNA strand displacement rate constant between the 5_1 leak product and the gates was varied to recapitulate the experimental kinetics. The reaction rate constant with the leak reaction was approximately two orders of magnitude lower than the rate constant for the reaction with the single-stranded 5_1 output. The rate constants are given inside the plots.

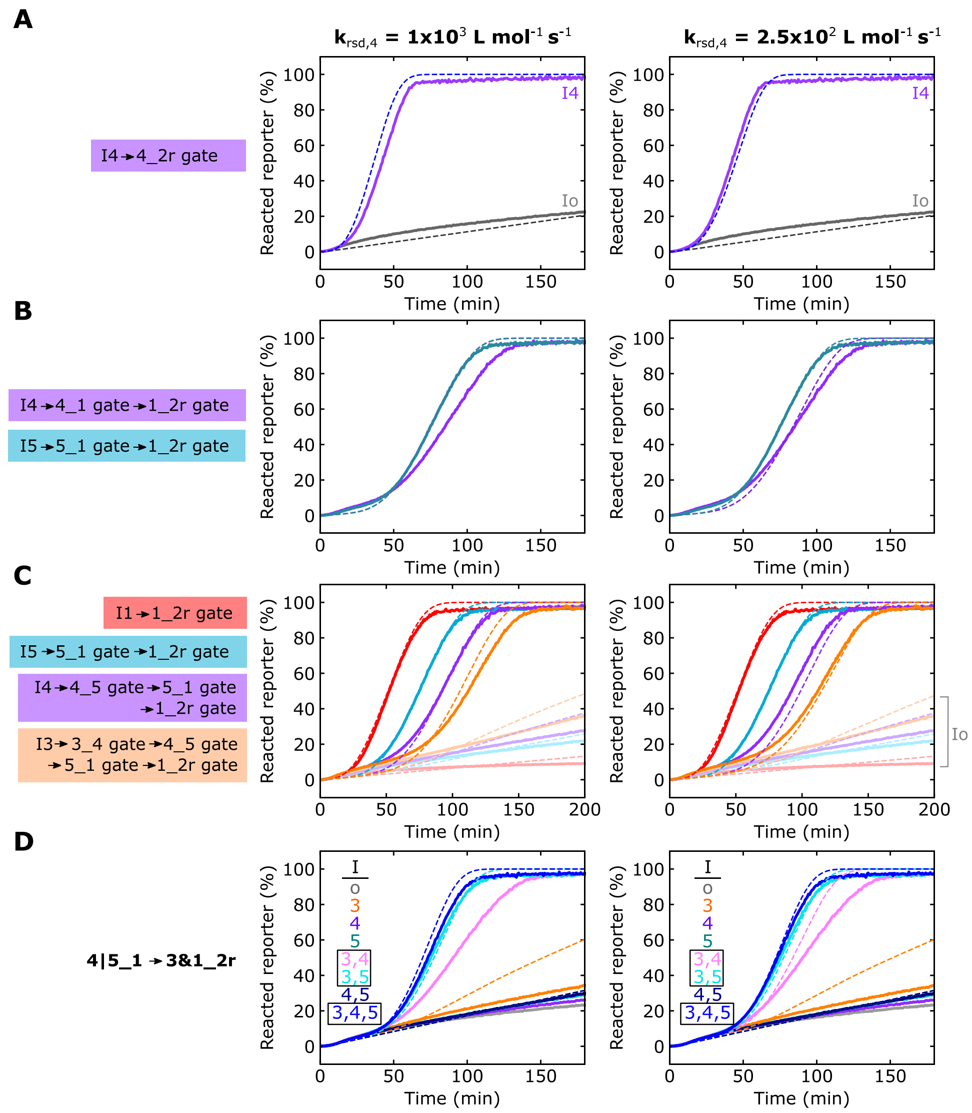


**Supplementary Figure 24:** Lowering $k_{rsd}$ for I4 4-fold better recapitulates experimental results. (**A**) The single-layer cascade of I4 to 4_2r from Figure 3A of the main text with simulation results using $k_{rsd}$ = 1x10$^3$ L mol$^{-1}$ s$^{-1}$ (left) or 2.5x10$^2$ L mol$^{-1}$ s$^{-1}$ (right). (**B**) A two-layer cascade of I4 to 4_1 to 1_2r (purple) or I5 to 5_1 to 1_2r (teal) with simulation results using $k_{rsd}$ = 1x10$^3$ L mol$^{-1}$ s$^{-1}$ (left) or 2.5x10$^2$ L mol$^{-1}$ s$^{-1}$ (right). (**C**) The four-layer cascade from Figure 5B of the main text with simulation results using $k_{rsd}$ = 1x10$^3$ L mol$^{-1}$ s$^{-1}$ (left) or 2.5x10$^2$ L mol$^{-1}$ s$^{-1}$ (right). (**D**) The logic cascade containing the 4_1 gate from Figure 5E with simulation results using $k_{rsd}$ = 1x10$^3$ L mol$^{-1}$ s$^{-1}$ (left) or 2.5x10$^2$ L mol$^{-1}$ s$^{-1}$ (right). All other rate constants are presented in Supplementary Table 3.

### 7. 1_2r gates with different toehold lengths

The kinetics of toehold-mediated strand displacement reactions can be controlled by toehold length. Here, we explore how toehold length influenced ctRSD kinetics. The initial design for the 1_2r gate included a 6 base single-stranded input toehold, which we would expect to result in a rate constant near the maximum theoretical limit ($10^6$ L $mol^{-1}$ $s^{-1}$) (*6*, *21*). However, our simulations indicated that the forward strand displacement rate constant between the 1_2r gate and I1 was only $10^3$ L $mol^{-1}$ $s^{-1}$. We theorized steric hinderance between the ribozyme and the input strand could result in slower strand displacement because the 6 base toehold is directly adjacent to the bulky HDV ribozyme motif (Supplementary Figure 1). Thus, in addition exploring the influence of toehold length on ctRSD kinetics, we also explored the influence of including a single-stranded spacer sequence between the ribozyme motif and the toehold. To do this, we designed 1_2r gates with (6, 8, 10, and 12) base toeholds and I1 variants possessing (4, 6, 8, or 10) base toeholds and combinatorially tested all gate and input combinations. Schematics with sequences are presented in Supplementary Figure 6. We first confirmed increasing toehold length did not influence gate folding and/or cleavage. Supplementary Figure 25 demonstrates that the 1_2r gate toehold variants cleaved as designed. Supplementary Figure 26 shows that increasing toehold length did not increase leak with the DNA reporter, indicating proper folding.

We next evaluated ctRSD kinetics for all gate and input toehold length combinations. These experiments encompassed toehold lengths of 4 bases to 10 bases with spacer lengths varying from (0 to 8) bases depending on the input toehold length (Supplementary Figure 27A,B). In these experiments, we were not able to resolve reaction rate constants greater than $10^5$ L $mol^{-1}$ $s^{-1}$ (Supplementary Figure 27C). When the strand displacement reaction rate gets this high, the overall rate of output release becomes limited by transcription and gate cleavage, rather than strand displacement. Thus, we report all reaction rate constants near this $10^5$ L $mol^{-1}$ $s^{-1}$ limit as ≥$10^5$ L $mol^{-1}$ $s^{-1}$ (Supplementary Figure 27B).

Supplementary Figure 27D show the kinetic traces for each gate and input toehold combination, highlighting the influence of spacer length on ctRSD kinetics for each input toehold length. Inclusion of a spacer generally increases reaction rate, but the spacer length that saturates the reaction rate decreases as input toehold length increases. An explanation for this observation could be: the weaker the input binding energy, the greater the influence of steric hinderance on the reaction rate. For example, the input with a 4 base toehold binds weakly to the 1_2r gate toehold, so a long spacer is required to completely remove any effect of steric hinderance. Conversely, for the input with a 10 base toehold, the same kinetics are observed for a 0 base and 2 base spacer. In this case, the input with the 10 base toehold can be viewed as an input with a 6 base toehold binding to a gate with a 4 base spacer, or an input with an 8 base toehold binding to a gate with a 2 base spacer. Both those reactions occur at a rate near the maximum value. Put another way, once the input toehold is long enough, increasing the length of the a-toehold (input) and a'-toehold (gate) together has almost the same effect as simply increasing the spacer length, *i.e.* increasing the a'-toehold (gate) without increasing the a-toehold (input). In support of this hypothesis, the 8 base a-toehold (input) and 8 base a'-toehold (gate) reaction rate constant is close to the 6 base a-toehold (input) and 8 base a'-toehold (gate) reaction rate constant (Supplementary Figure 27B).

Supplementary Figure 27E shows the kinetic traces for each gate and input toehold combination, highlighting the influence of toehold length on ctRSD kinetics for each spacer length. With the exception of the input with a 4 base a-toehold, most of the changes in kinetics observed across toehold length can be attributed to the increase in a'-toehold (spacer) length. For example, with long enough spacers, inputs with (6, 8, and 10) base toeholds exhibit strand displacement constants close to the maximum value (≥$10^5$ L $mol^{-1}$ $s^{-1}$).

How do these results compare to previous studies of toehold-mediated strand displacement kinetics? For traditional DNA (*21*) and RNA strand displacement (*6*), in which double-stranded complexes are pre-annealed and gate toeholds have no secondary structure upstream, toeholds ≥6 bases should result in reaction rate constants at the theoretical maximum of ≈$10^6$ L $mol^{-1}$ $s^{-1}$. We found similar results for ctRSD when using a long enough spacer between the HDV ribozyme and the toehold. Regarding the input with a 4 base a-toehold, the reaction between this input and any of the 1_2r gates has a much lower thermodynamic driving force than the other input toeholds tested. This is because the 1_2r gates all possess a 6 base reverse toehold, *i.e.* completion of the forward strand displacement reaction results in a net loss of two base pairs compared to the intact 1_2r gate. The rate constant for a DNA strand displacement reaction between an input with a 4 base toehold and a gate with a 6 base reverse toehold (b-toehold) was measured to be between ($10^2$ and $10^3$) L $mol^{-1}$ $s^{-1}$. This aligns with our estimated rate constant of 2x$10^2$ L $mol^{-1}$ $s^{-1}$ for ctRSD between the 4 base input toehold variant and gates with either a 6 base or 8 base spacers (Supplementary Figure 27B). Together, these results suggest that ctRSD should possess the same kinetic control of traditional toehold-mediated strand displacement, provided appropriate spacers are used.

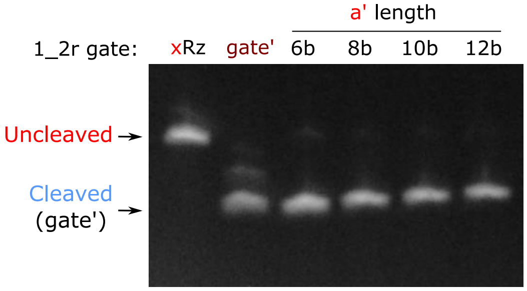


**Supplementary Figure 25:** Denaturing gel of 1_2r gates with (6, 8, 10, and 12) base a'-toeholds. All gates self-cleave as designed. Transcription and gel electrophoresis were conducted as described in the Methods of the main text.

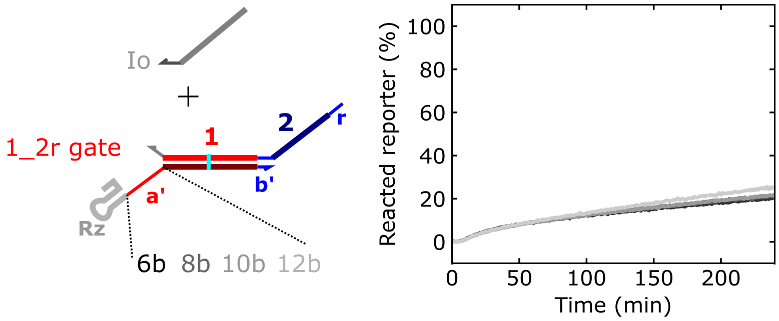


**Supplementary Figure 26:** Normalized reporter signal during co-transcription of Io and 1_2r gate variants with differing a'-toehold lengths. Increasing toehold length did not increase the observed amount of downstream reporter leak. [1_2r gate templates] = 5 nmol/L, [Io template] = 40 nmol/L, [DNA reporter] = 150 nmol/L, and [T7 RNAP] = 1 U/µL.

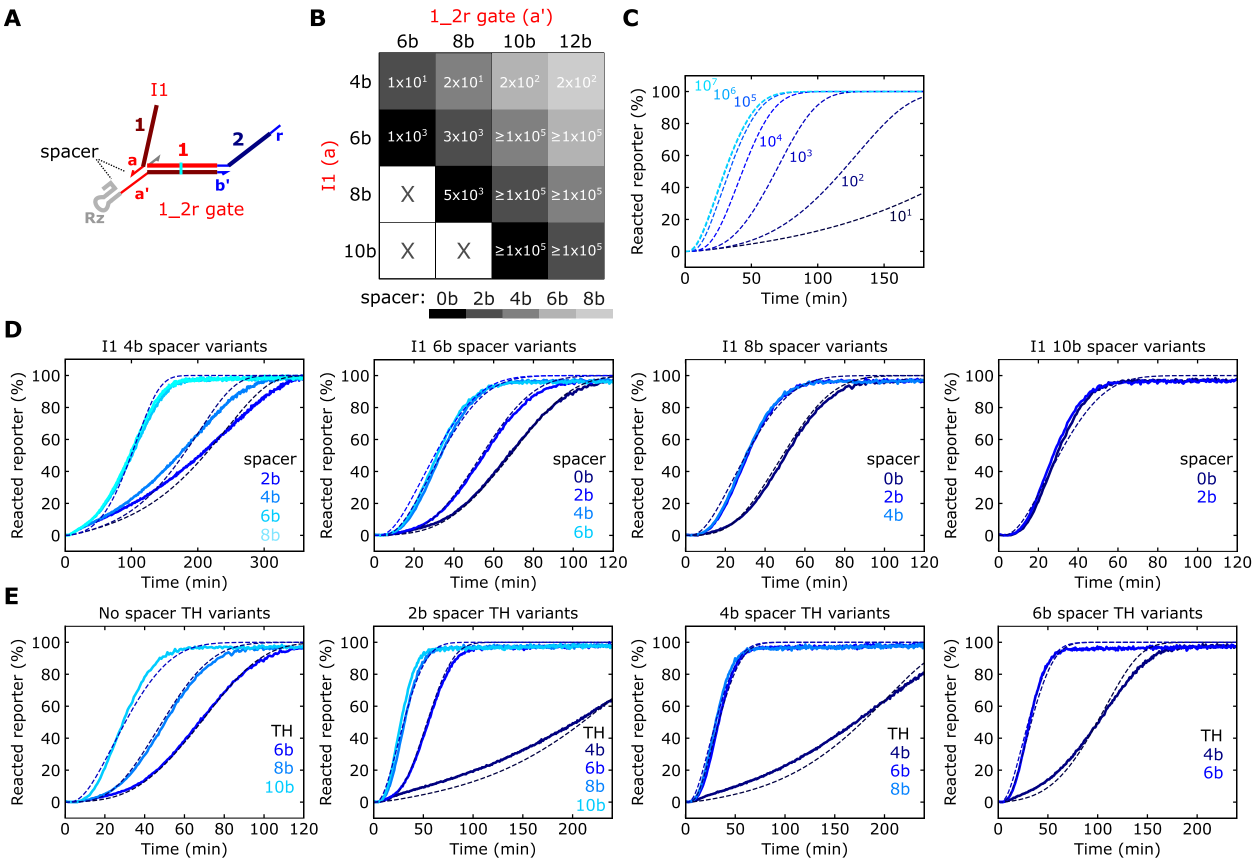


**Supplementary Figure 27:** ctRSD kinetics depend on toehold length and the length of a single-stranded spacer after the self-cleaving ribozyme. (**A**) Schematic of I1 bound to a 1_2r gate with a single-stranded spacer. (**B**) Grid representing the combination of I1 a-toehold lengths and 1_2r gate a'-toehold lengths tested in experiments. Box shading represents the length of the spacer for each combination. The numbers inside the boxes are the $k_{rsd}$ reaction rate constants, in units of L mol$^{-1}$ s$^{-1}$, that recapitulated the experimental data for each combination. Under the experimental conditions, estimates of reaction rate constants >1x10$^5$ L mol$^{-1}$ s$^{-1}$ could not be distinguished (see panel C) and are reported as ≥ 1x10$^5$ L mol$^{-1}$ s$^{-1}$. (**C**) Simulations of the ctRSD reaction with $k_{rsd}$ rate constants ranging six orders of magnitude. These simulations show rate constants > 1x10$^5$ L mol$^{-1}$ s$^{-1}$ result in similar kinetics and are indistinguishable in this assay. (**D**) Normalized reporter signal for the I1 and 1_2r gate combinations in (B). Here, each plot shows a single I1 variant with different 1_2r gate variants to explore the effect of spacer length on ctRSD kinetics. Each plot represents a single row from the grid in (B). (**E**) Normalized reporter signal for the I1 and 1_2r gate combinations in (B). Here, each plot shows I1 and 1_2r gate variant combinations with constant spacer lengths to explore the effect of toehold length on ctRSD kinetics. Each plot represents a left to right diagonal of equal shading from the grid in (B). [1_2r gate templates] = 5 nmol/L, [I1 templates] = 5 nmol/L, [DNA reporter] = 150 nmol/L, and [T7 RNAP] = 1 U/µL. Io was added to bring the total template concentration up to 45 nmol/L in each sample. For the simulations, $k_p$ of 0.019 s$^{-1}$ was used, and the $k_{rsd}$ reaction rate constants were calibrated to the experimental kinetics. The simulation results in (D and E) used the $k_{rsd}$ values tabulated in (B). Schematics with sequences of the input and gate variants are presented in Supplementary Figure 6.

### 8. Potential advantages of ctRSD circuits compared to DNA-based circuits

Should ctRSD circuits continue to prove as predictable and programmable as DNA-based circuits, ctRSD could serve as a more versatile alternative to DNA computing. Such a shift could be justified given the high fidelity and decreasing price of gene synthesis. Integrated DNA Technologies currently reports ≈80 % of 30 base DNA oligonucleotides are the correct product compared to ≈100 % for gBlocks of >125 bases (*29*). The low fidelity of DNA oligonucleotide synthesis requires the strands to be purified with gel electrophoresis and many DNA computing papers report the purification of individual dsDNA circuit complexes to obtain desired circuit function (*8*, *30*). For ctRSD the high-fidelity gBlock synthesis is followed by a high-fidelity PCR step (<0.25 % error (*31*)) and high-fidelity transcription–T7 RNAP's nucleotide substitution rate is less than 1 in 17,000 bases (*32*). Further, encoding the dsRNA complex as a single transcript ensures the proper stoichiometry between the two gate strands, reducing leak pathways (*33*). Thus, ctRSD removes the need for purification of circuit components before operation, greatly simplifying the workflow. Further, the per nanomole cost of a ctRSD gate template can be reduced to nearly that of analogous DNA gates with a few modifications to the protocol presented here (Supplementary Section 8.1).

Another advantage of using transcriptionally encoded circuits over DNA strand displacement circuits is the long-term stability of long linear DNA templates and DNA plasmids. For example, in many biosensor (*34*) and diagnostic (*35*) applications, circuit components are freeze dried for long-term storage and ease of transportation. These freeze-dried circuits are then activated by adding a liquid sample at the point of need. Both linear DNA templates on the order of 300 bases and DNA plasmids have been shown to remain stable for months after freeze drying. Short DNA strand displacement duplexes show significant decrease in performance only one week after freeze drying (*36*).

#### 8.1 Cost analysis

<u>Cost of preparing DNA templates for ctRSD:</u>

In this study, the gBlocks were price fixed from (125 to 500) bases and were $89 USD each for 250 ng. The gBlocks were resuspended in 25 µL of Buffer EB. For PCR, 1.5 µL of gBlock DNA was used in a 75 µL reaction and typically yielded 50 µL of 600 nmol/L product after purification. This is 30 pmol of DNA template per reaction, and 16.67 reactions can be conducted using the 25 µL of gBlock DNA, resulting in 0.5 nmol of total DNA template for a given gBlock order. This comes to $178 per nmol of template. The Phusion Master-Mix used for gBlock PCRs cost $782 for 500 reactions, which is $1.56 per reaction and $26 to produce all 0.5 nmol of DNA. This results in an additional $52 per nmol of DNA. Finally, the cost of the DNA primers used in the PCR must be added. These 25 base oligos cost $9.25 for 180 µg and 0.5 µmol/L (0.31 µg) were used in each reaction, which is $0.016 per reaction and $0.26 to produce the total 0.5 nmol of product ($0.52 per nmol). In total, the ctRSD DNA templates cost: $178 per nmol for the gBlock DNA, $52 per nmol of DNA for the Phusion Master-Mix, and $0.52 per nmol for the DNA primers = **$230.52 per nmol**.

Note that the main cost is the initial purchase of the gBlock DNA. This cost could be significantly reduced by purchasing the DNA in bulk as eBlocks from IDT. eBlocks require an order of at least 24 sequences but cost only $0.06 per base, which is $18 for a 300 base template (the minimum length for ordering). This would drop the cost to **$88.52 per nmol** of DNA template. Further, a gBlock or eBlock template only needs to be ordered a single time from IDT, and once the DNA from that order has been exhausted, the

PCR products of the gBlock DNA can themselves be PCR amplified to produce more template. A single gBlock PCR could be used to conduct 250 more PCR experiments, which yields 7.5 nmol of DNA template. If PCR of a PCR amplified gBlock is conducted just once, the cost for a DNA template is dropped to $11.87 per nmol for gBlocks and $2.4 per nmol for eBlocks. This amounts to **$64.39 and $54.94**, respectively. Here the primary cost is from PCR reagents.

<u>Cost of preparing a DNA gates for DNA strand displacement:</u>

For DNA oligos, IDT charges $0.70 for oligos at the 100 nmol scale, which is required for purification. If these oligos are ordered with polyacrylamide gel electrophoresis (PAGE) purification, the added cost is $60. PAGE purification has a guaranteed yield of 2 nmol of final product.

A PAGE purified 26 base oligo that serves as the output strand of a DNA gate thus costs $78.2 for a guaranteed 2 nmol of DNA or $39.10 per nmol. A 22 base gate' strand is $75.40 for 2 nmol ($37.7 per nmol). Thus, at the guaranteed yield, the cost is **$76.80 per nmol of gate**.

### 9. Experimental conditions, transcription rate calibration, and experimental variability

Supplementary Table 4 contains transcription template and T7 RNAP concentrations used in DNA reporter assays. In our experiments, the transcription rate was dependent on the total transcription template and T7 RNAP concentrations (Supplementary Figure 28). To ensure the same transcription load across different samples for the same ctRSD element or circuit, a template producing an unreactive input (Io) was added so that all samples had the same total template concentration. The total concentration of templates for each experiment is also presented in Supplementary Table 4. Because the transcription rate differed across many experiments, the first order rate constant ($k_p$) used to model transcription had to be calibrated for a given T7 RNAP and total template concentration (Supplementary Figure 29). The first order rate constant ($k_p$) calibrated for each experiment is presented in Supplementary Table 4. This transcription rate calibration should also calibrate for batch to batch variation in T7 RNAP stocks. Other than the differences arising from different experimental conditions or T7 RNAP batches, replicate measurements of circuit kinetics varied between 2 % to 5 % (Supplementary Section 9.3).

### 9.1 Experimental conditions

**Supplementary Table 4:** Transcription template and T7 RNAP concentrations used in DNA reporter experiments. 500 nmol/L of DNA reporter was used in each experiment. For experiments in which multiple input template concentrations or input template combinations were tested, each column of values represents the nominal concentrations of each input template for each experiment.

| Figure 2d (I1 titration) | Values |
|---|---|
| [1_2r gate] (nmol/L) | 25 |
| [I1] (nmol/L) | 0, 2.5, 5,12.5,25,50 |
| [Io] (nmol/L) | 50,47.5,45,37.5,25, 0 |
| [T7 RNAP] (U/µL) | 1.0 |
| *Total [templates]* (nmol/L) | *75* |
| $k_p$ ($s^{-1}$) | *0.013* |
| Figure 3a (orthogonal sequences) | Values |
| [Gates] (nmol/L) | 25 |
| [Ii] (nmol/L) | 0,50 |
| [Io] (nmol/L) | 50,0 |
| [T7 RNAP] (U/µL) | 1.0 |
| *Total [templates]* (nmol/L) | *75* |
| $k_p$ ($s^{-1}$) | *0.013* |
| Figure 4c (1\|3_2r) | Values |
| [1_2r gate] (nmol/L) | 25 |
| [3_2r gate] (nmol/L) | 25 |
| [I1] (nmol/L) | 0,50, 0,50 |
| [I3] (nmol/L) | 0, 0,50,50 |
| [Io] (nmol/L) | 100,50,50, 0 |
| [T7 RNAP] (U/µL) | 1.2 |
| *Total [templates]* (nmol/L) | *150* |
| $k_p$ ($s^{-1}$) | *0.008* |
| Figure 4f (3&1_2r) | Values |
| [3&1_2r gate] (nmol/L) | 25 |
| [I1] (nmol/L) | 0,50, 0,50 |
| [I3] (nmol/L) | 0, 0,50,50 |
| [Io] (nmol/L) | 100,50,50, 0 |
| [T7 RNAP] (U/µL) | 1.0 |
| *Total [template]* (nmol/L) | *125* |
| $k_p$ ($s^{-1}$) | *0.009* |
| Figure 4i (Catalytic Amp) | Values |
| [1_2r gate] (nmol/L) | 25 |
| [I1] (nmol/L) | 1.25,2.50,1.25,2.50 |
| [F1] (nmol/L) | 0, 0, 25, 25 |
| [Io] (nmol/L) | 26.25, 25,1.25, 0 |
| [T7 RNAP] (U/µL) | 1.2 |
| *Total [template]* (nmol/L) | *52.5* |
| $k_p$ ($s^{-1}$) | *0.010* |
| Figure 5b (4-layer cascade) | Values |
| [1_2r gate] (nmol/L) | 25, 25, 25,25,25,25,25,25 |
| [5_1 gate] (nmol/L) | 0, 25, 25,25,0, 25,25,25 |
| [4_5 gate] (nmol/L) | 0, 0, 25,25,0, 0, 25,25 |
| [3_4 gate] (nmol/L) | 0, 0, 0, 25,0, 0, 0, 25 |
| [I1] (nmol/L) | 0, 0, 0, 0, 50,0, 0, 0 |
| [I3] (nmol/L) | 0, 0, 0, 0, 0, 0, 0, 50 |
| [I4] (nmol/L) | 0, 0, 0, 0, 0, 0, 50,0 |
| [I5] (nmol/L) | 0, 0, 0, 0, 0, 50,0, 0 |
| [Io] (nmol/L) | 125,100,75,50,75,50,25,0 |
| [T7 RNAP] (U/µL) | 1.5 |
| *Total [template]* (nmol/L) | *150* |
| $k_p$ ($s^{-1}$) | *0.0075* |

| Figure 5c (1\|3\|4\|5_2r) | Values |
|---|---|
| [1_2r gate] (nmol/L) | 12.5 |
| [3_2r gate] (nmol/L) | 12.5 |
| [4_2r gate] (nmol/L) | 12.5 |
| [5_2r gate] (nmol/L) | 12.5 |
| [I1] (nmol/L) | 0,50, 0, 0,0 |
| [I3] (nmol/L) | 0, 0, 50, 0,0 |
| [I4] (nmol/L) | 0, 0, 0,50,0 |
| [I5] (nmol/L) | 0, 0, 0, 0,50 |
| [Io] (nmol/L) | 50, 0, 0, 0,0 |
| [T7 RNAP] (U/µL) | 1.2 |
| *Total [template]* (nmol/L) | *100* |
| $k_p$ ($s^{-1}$) | *0.015* |
| **Figure 5d (5&4_1 to 3&1_2r)** | **Values** |
| [5&4_2r gate] (nmol/L) | 25 |
| [3&1_2r gate] (nmol/L) | 25 |
| [I3] (nmol/L) | 0,25, 0, 0,25,25, 0,25 |
| [I4] (nmol/L) | 0, 0,25, 0,25, 0,25,25 |
| [I5] (nmol/L) | 0, 0, 0,25, 0,25,25,25 |
| [Io] (nmol/L) | 75,50,50,50,25,25,25, 0 |
| [T7 RNAP] (U/µL) | 1.2 |
| *Total [template]* (nmol/L) | *125* |
| $k_p$ ($s^{-1}$) | *0.0075* |
| **Figure 5e (4\|5_1 to 3&1_2r)** | **Values** |
| [4_1 gate] (nmol/L) | 25 |
| [5_1 gate] (nmol/L) | 25 |
| [3&1_2r gate] (nmol/L) | 25 |
| [I3] (nmol/L) | 0,25, 0, 0,25,25, 0,25 |
| [I4] (nmol/L) | 0, 0,25, 0,25, 0,25,25 |
| [I5] (nmol/L) | 0, 0, 0,25, 0,25,25,25 |
| [Io] (nmol/L) | 75,50,50,50,25,25,25, 0 |
| [T7 RNAP] (U/µL) | 1.5 |
| *Total [template]* (nmol/L) | *150* |
| $k_p$ ($s^{-1}$) | *0.010* |
| **Figure 5f (5&4_1 to 1\|3_2r)** | **Values** |
| [5&4_2r gate] (nmol/L) | 25 |
| [1_2r gate] (*nmol/L*) | 25 |
| [3_2r gate] (*nmol/L*) | 25 |
| [I3] (*nmol/L*) | 0,25, 0, 0,25,25, 0,25 |
| [I4] (*nmol/L*) | 0, 0,25, 0,25, 0,25,25 |
| [I5] (*nmol/L*) | 0, 0, 0,25, 0,25,25,25 |
| [Io] (*nmol/L*) | 75,50,50,50,25,25,25, 0 |
| [T7 RNAP] (U/µL) | 1.5 |
| *Total [template] (nmol/L)* | *150* |
| $k_p$ ($s^{-1}$) | *0.010* |

### 9.2 Transcription rate calibration

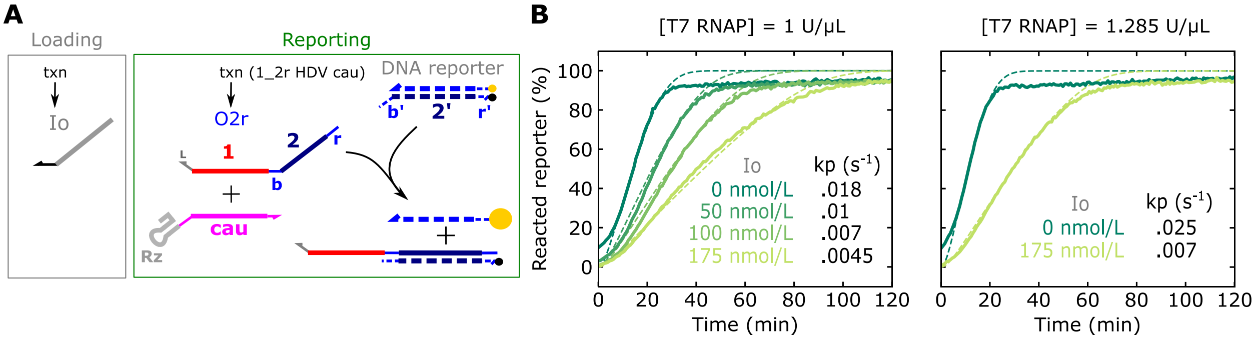


**Supplementary Figure 28:** Total template concentration and T7 RNAP concentration influence transcriptional load. (**A**) Schematic of the loading and reporting reactions. The Io template was added at different concentrations to change the total transcriptional load, and the O2r strand was constitutively produced from the 1_2r HDV cau template. The 1_2r HDV cau template was chosen to match the total number of bases in the 1_2r gate template. (**B**) Experimental (solid lines) and simulated (dashed lines) reporter signal during transcription of the 1_2r HDV cau RNA with different Io template concentrations and T7 RNAP concentrations. The $k_p$ values used in the simulations are tabulated for the different Io template concentrations and T7 RNAP concentrations. Increasing the Io template concentration (transcriptional load) decreases the effective $k_p$ value and increasing the T7 RNAP concentration increases the effective $k_p$ value. 25 nmol/L of the 1_2r HDV cau template and 500 nmol/L of the DNA reporter were used in each experiment. Reactions were otherwise conducted as described in the Methods of the main text.

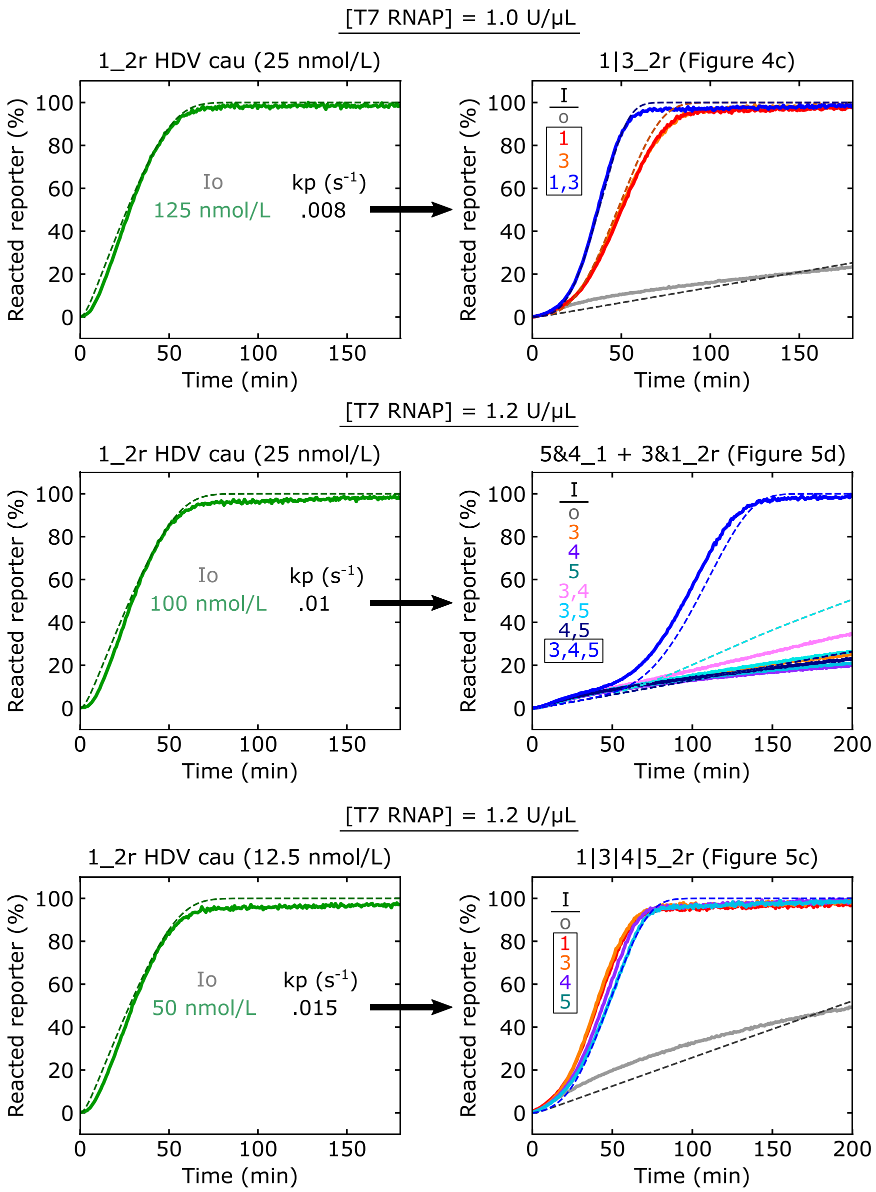


**Supplementary Figure 29:** Representative examples showing calibration of the transcription rate constant for experiments with different transcriptional loads and/or T7 RNAP concentrations. The 1_2r HDV cau template, which constitutively produces the O2r strand, was used as a reference sample for each experiment. The Io template was added to bring the total template concentration in the reference sample to the total concentration used in the experimental samples. The reference sample was then used to calibrate the first order transcription rate constant, $k_p$, for simulation of the experimental samples.

### 9.3 Experimental variability

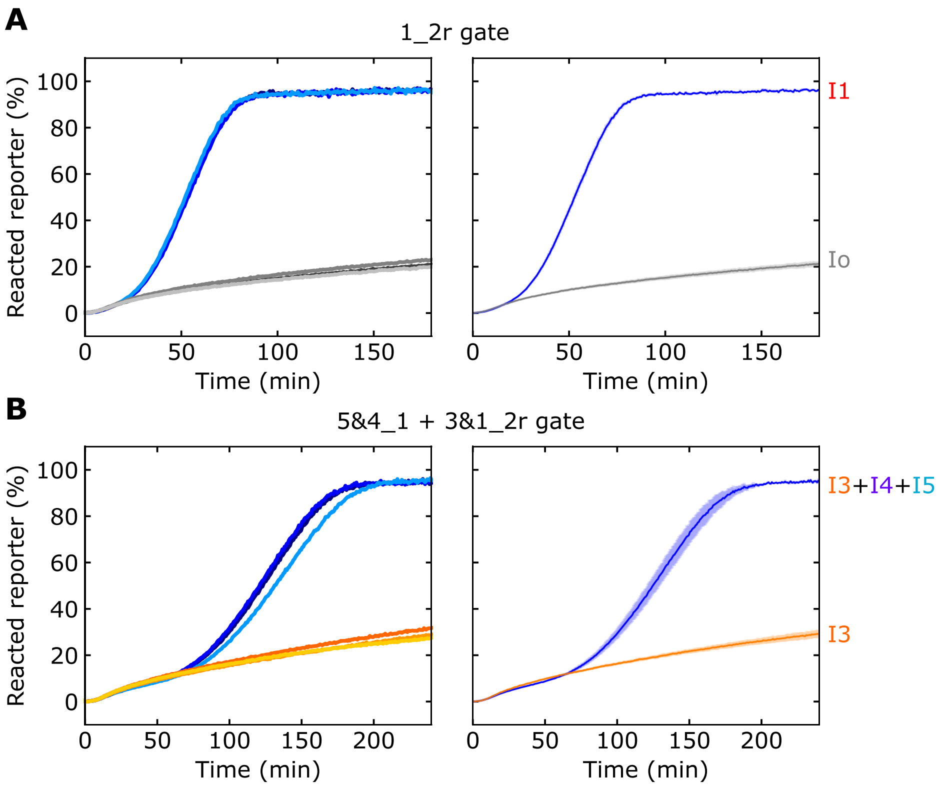


**Supplementary Figure 30:** Variation across independently prepared technical replicates for ctRSD reactions using the 1_2r gate (**A**) or the 5&4_1+3&1_2r gates (**B**). Left plots show data for three independently prepared replicates of the same ctRSD reaction. The replicates were prepared from the same stock solutions on the same day. The right plots show the mean of the three replicates; error bars represent one standard deviation. The standard deviation was < 1.5 % from the mean value at each time point for (A) and < 5 % from the mean value at each time point for (B). In (A), reactions included 25 nmol/L of the 1_2r gate template, 12.5 nmol/L of the I1 template, and Io template to bring the total template to 50 nmol/L in both reactions. In (B), reactions included 25 nmol/L of the 5&4_1, 25 nmol/L of the 3&1_2r gate, and 25 nmol/L of the input templates. For the samples containing I3 only, 50 nmol/L of the Io template was added to bring the total template concentration to 75 nmol/L. In all experiments, 500 nmol/L of the DNA reporter and 2 U/µL of T7 RNAP was used.

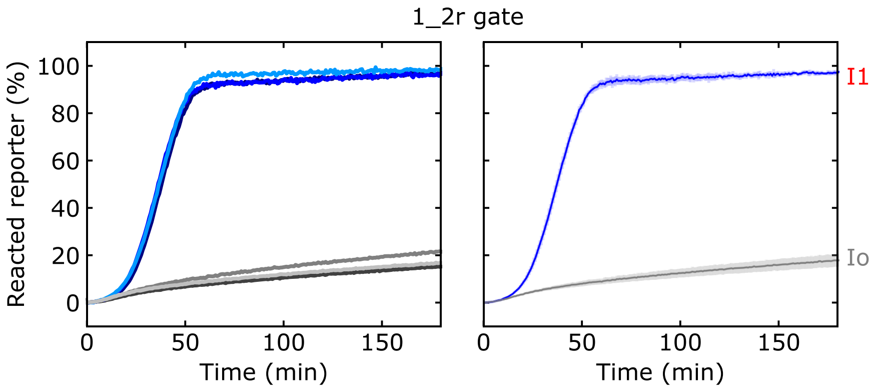


**Supplementary Figure 31:** Variation across experiments performed on different days for the same set of conditions. Left plots show data for three independent replicates of the same ctRSD reaction. Reactions included 25 nmol/L of the 1_2r gate template, 500 nmol/L of the DNA reporter, 1 U/µL of T7 RNAP, and 50 nmol/L of the I1 or Io templates. The three individual replicates were prepared independently and tested on separate days, with the second and third replicates conducted 5 and 19 days after the first replicate, respectively. Dark colored lines represent the oldest replicate; data also presented in Supplementary Figure 17B, middle panel. Medium colored lines represent the second oldest replicate; data also presented in Figure 2D, 2x I1, of the main text. Light colored lines represent the newest replicate; data also presented in Supplementary Figure 15B. The right plot shows the mean of the three replicates; error bars represent one standard deviation. The standard deviation is < 3 % from the mean value at each time point.
