## Supplementary material for "Co-transcriptional RNA strand displacement circuits": Fluorescence data: figure2D_fluorescence_data_plotting.py

```
import numpy as np
import scipy.integrate as spi
import matplotlib as mpl
import matplotlib.pyplot as plt
from matplotlib.ticker import MaxNLocator
from matplotlib import cm
import openpyxl as opxl
import xlrd
    
'''
##############################################################################
Loading data (Normalized)
##############################################################################
'''
# In these experiments, the maximum fluorescence value for normalization was 
# taken from each individual sample, except for the Io sample which did not
# reach a maximum
# For the Io sample, the maximum fluorescence value from the 25 nmol/L I1 
# sample was used

save_loc = 'C:\\Users\\sws5\\OneDrive - National Institute of Standards and Technology (NIST)\\Presentations and write ups\\Papers\\RSD paper\\data upload\\fluorescence data\\'
save_name = 'ctRSD_data_upload'

wb = xlrd.open_workbook(save_loc+save_name+'.xlsx') # saved data file

ws = wb.sheet_by_name('Fig. 2D') # sheet to import data from

# importing normalized data
t = np.zeros(ws.nrows-3)
for r in range(3,ws.nrows):
    t[r-3] = ws.cell_value(r,0)
    
DDn = np.zeros([ws.nrows-3,int(ws.ncols/2)-1])

for c in range(1,int(ws.ncols/2)):
    for r in range(3,ws.nrows):
        DDn[r-3,c-1] = ws.cell_value(r,c)
        
# importing raw data    
DDnR = np.zeros([ws.nrows-3,int(ws.ncols/2)-1])

for c in range(8,int(ws.ncols)):
    for r in range(3,ws.nrows):
        DDnR[r-3,c-8] = ws.cell_value(r,c)
        
'''
###############################################################################
PLOTTING
###############################################################################
'''

fs = 12 # font size for plotting
ts = 44 # shifting time to when T7 RNAP was added to samples

# Normalized fluorescence
plt.figure(1)
plt.rcParams.update({'font.sans-serif':'Verdana'})
plt.subplot(3,5,1)
plt.plot(t-ts,DDn[:,0]*100,color=[0.6,0.6,0.6],linewidth=2)
plt.plot(t-ts,DDn[:,1]*100,color=[0,0,0.33],linewidth=2)
plt.plot(t-ts,DDn[:,2]*100,color=[0,0,0.66],linewidth=2)
plt.plot(t-ts,DDn[:,3]*100,color=[0,0,1],linewidth=2)
plt.plot(t-ts,DDn[:,4]*100,color=[0,0.33,1],linewidth=2)
plt.plot(t-ts,DDn[:,5]*100,color=[0,0.66,1],linewidth=2)
plt.xticks(fontsize=fs)
plt.yticks(fontsize=fs)
plt.ylim(-10,110)
plt.xlim(0,180)
ax1 = plt.gca()
ax1.xaxis.set_tick_params(which='both',size=3,width=1,direction='in',top='on')
#ax1.xaxis.set_major_locator(MaxNLocator(integer=True))
ax1.yaxis.set_tick_params(which='both',size=3,width=1,direction='in',right='on')
plt.xlabel('Time (min)',fontsize=fs)
plt.ylabel('Reacted reporter (%)',fontsize=fs)

# Raw fluorescence
plt.figure(1)
plt.rcParams.update({'font.sans-serif':'Verdana'})
plt.subplot(3,5,3)
plt.plot(t-ts,DDnR[:,0],color=[0.6,0.6,0.6],linewidth=2)
plt.plot(t-ts,DDnR[:,1],color=[0,0,0.33],linewidth=2)
plt.plot(t-ts,DDnR[:,2],color=[0,0,0.66],linewidth=2)
plt.plot(t-ts,DDnR[:,3],color=[0,0,1],linewidth=2)
plt.plot(t-ts,DDnR[:,4],color=[0,0.33,1],linewidth=2)
plt.plot(t-ts,DDnR[:,5],color=[0,0.66,1],linewidth=2)
plt.xticks(fontsize=fs)
plt.yticks(fontsize=fs)
#plt.ylim(-10,110)
plt.xlim(0,180)
ax1 = plt.gca()
ax1.xaxis.set_tick_params(which='both',size=3,width=1,direction='in',top='on')
#ax1.xaxis.set_major_locator(MaxNLocator(integer=True))
ax1.yaxis.set_tick_params(which='both',size=3,width=1,direction='in',right='on')
plt.xlabel('Time (min)',fontsize=fs)
plt.ylabel('Fluorescence (A.U.)',fontsize=fs)
```
