## Supplementary material for "Co-transcriptional RNA strand displacement circuits": Fluorescence data: notes.txt

The ctRSD_data_upload.xlsx file contains both the raw and normalized fluorescence data
from: Schaffter, SW and Strychalski, EA. Co-transcriptional RNA strand displacement circuits (2021)

In the Excel file, rows highlighted in green indicate the times when T7 RNAP was added to the samples
Plots of data in the manuscript designate this time as time = 0

The figure2D_fluorescence_data_plotting.py file is an example showing how the data was imported from Excel
and plotted in the manuscript
In the script, the save_loc variable will need to be changed to the path pointing
to where the ctRSD_data_upload.xlsx file is saved on a users local computer
